# Digitize your Biology! Modeling multicellular systems through interpretable cell behavior

**DOI:** 10.1101/2023.09.17.557982

**Authors:** Jeanette A.I. Johnson, Genevieve L. Stein-O’Brien, Max Booth, Randy Heiland, Furkan Kurtoglu, Daniel R. Bergman, Elmar Bucher, Atul Deshpande, André Forjaz, Michael Getz, Ines Godet, Melissa Lyman, John Metzcar, Jacob Mitchell, Andrew Raddatz, Heber Rocha, Jacobo Solorzano, Aneequa Sundus, Yafei Wang, Danielle Gilkes, Luciane T. Kagohara, Ashley L. Kiemen, Elizabeth D. Thompson, Denis Wirtz, Pei-Hsun Wu, Neeha Zaidi, Lei Zheng, Jacquelyn W. Zimmerman, Elizabeth M. Jaffee, Young Hwan Chang, Lisa M. Coussens, Joe W. Gray, Laura M. Heiser, Elana J. Fertig, Paul Macklin

## Abstract

Cells are fundamental units of life, constantly interacting and evolving as dynamical systems. While recent spatial multi-omics can quantitate individual cells’ characteristics and regulatory programs, forecasting their evolution ultimately requires mathematical modeling. We develop a conceptual framework—a cell behavior hypothesis grammar—that uses natural language statements (cell rules) to create mathematical models. This allows us to systematically integrate biological knowledge and multi-omics data to make them computable. We can then perform virtual “thought experiments” that challenge and extend our understanding of multicellular systems, and ultimately generate new testable hypotheses. In this paper, we motivate and describe the grammar, provide a reference implementation, and demonstrate its potential through a series of examples in tumor biology and immunotherapy. Altogether, this approach provides a bridge between biological, clinical, and systems biology researchers for mathematical modeling of biological systems at scale, allowing the community to extrapolate from single-cell characterization to emergent multicellular behavior.

## INTRODUCTION AND BACKGROUND

Every cell has an identity and internal state defined by its DNA, epigenetic profile, and molecular signaling pathways. Quantifying those internal state variables and how they change due to signals received from their environment has enabled the classification of cell types and behaviors^1–4^. However, the necessity of destroying the cell to obtain those measurements results in static snapshots of a single moment in time. Thus, the ability to connect those snapshots into cellular movies remains an area of strong community interest and an open computational challenge^5–7^. The large number of cellular and molecular interactions within biological systems make temporal predictions impossible to perform solely through experimentation. Techniques such as trajectory inference and RNA velocity can estimate transitions of cellular phenotypes^8^, but they cannot account for more complex temporal changes throughout the diverse cellular ecosystems of biological systems. While machine learning from single-cell datasets can make these predictions for individual cell types^9,10^, they generally cannot account for changes resulting from cell-cell interactions. Moreover, these predictive methods are data-driven and unable to predict cellular states from prior biological knowledge or mechanism alone. Beyond filling the gaps between measurement times, computational techniques for temporal analysis of cellular states provide the potential to predict the future state of multicellular systems. More advanced computational tools are needed to model unseen emergent behaviors in complex biological systems comprised of multiple cell types.

Mechanistic mathematical modeling can address the key challenge of extending from static high-resolution data to forecasting the dynamics of multicellular systems. However, mathematical modeling lacks the language to *directly* connect to the vast accumulated knowledge of the biological community, nor to easily trans-form data into equations. In this paper, we develop a conceptual framework—a cell behavior hypothesis grammar—that bridges the divide between biology and mathematical modeling, by embracing well-defined human language hypotheses on cell behavior as a logical model that is fully equivalent to a mathematical model. This one-for-one relationship between human language and mathematics allows us to systematically curate and integrate both biological knowledge and high-throughput data, and makes them *computable.* This, in turn, enables virtual “thought experiments”^11^ that challenge and extend our understanding of multicellular systems, and ultimately generate new testable hypotheses.

Agent-based modeling is a powerful mathematical modeling technique that enables prediction of emergent complex behaviors by populations of individual “agents” in a system as they follow predefined rules based on their identity, state, and nearby conditions^12^. Agent-based models (ABMs) are well-suited to studying the dynamics of multicellular biology, as they can map a cell type to their set of internal states and rules of behavior^13–16^. At each time increment, each agent evaluates its surroundings and internal state variables and considers this information to calculate its next action. When applied to individual cells, each agent’s rules can represent hypotheses of single-cell behaviors, including their actions upon or in response to nearby cells (i.e., cell-cell interactions). By encoding the rules and relationships governing the stochastic behavior of cellular agents in complex systems, ABMs empower *in silico* experimentation and modeling of multiscale dynamic knowledge representations even in the absence of temporal measurements^14,17^.

ABMs can thus be used to test hypotheses in human development and disease where comprehensive, subject based experimentation is not possible. Applications to understanding human disease and in particular cancer have already been proven particularly powerful^18–33^. By predicting the future state of cells and the impact of perturbations, ABMs provide a powerful toolset to generate digital twins and virtual clinical trials^7,15,34–43^. Furthermore, the ability to run simulations at scale across diverse biological conditions with ABMs can refine biological understanding and predict future cellular behaviors in these complex systems. As a result, they provide a computational means to prioritize bench experiments or clinical trials to address the costs and practical constraints of real-world experimentation.

Widespread application of ABMs for modeling biological systems currently has two major limitations; the highly technical nature of most software implementations, and the limited ability to integrate molecular data to ground simulations in the real-world. The former issue gatekeeps ABM away from those without significant mathematical and computer-programming experience. This technical requirement limits widespread application, including by many potential users who have extensive working knowledge of the biological system they are investigating. Even for users with advanced computational expertise, the custom coding required can limit reproducibility. Disease etiology and operating biological hypotheses are often hidden deep in source code, obscuring the assumptions about the system and, thus, the full set of hypotheses that are being simulated and tested. These technical limitations to ABM modeling also limit the ability to embed molecular datasets, which are often too high-dimensional to manually encode into equations of agents and rules. A conceptual framing that can generally abstract cellular phenotypes and their interactions—combined with a simplifying coding infrastructure—is essential for the integration of molecular measurements to personalize model predictions.

Here, we sought a way for an expert’s knowledge base to be condensed and standardized into a set of clear biological statements that can be readily mapped onto biophysically motivated ABM parameters that describe their biological system, with or without molecular and cellular measurements. Facilitating this requires an intuitive language to concisely express expert knowledge as plain text descriptions of the rules of cell interactions that “encode” a system of study, and software to translate these plain text descriptions into mathematical expressions and executable models for immediate exploration of a digitized copy of the biological system^44^. While we use PhysiCell^44^ as a reference implementation, the grammar can independently represent expert knowledge for further analysis or import into other simulation frameworks^45–50^. Thus, the grammar allows for broad application of ABMs and addresses the need to be reproducible, modular, and extensible. We demonstrate how this grammar can encode expert biological knowledge through sample models in cancer biology spanning tumor cell growth, invasion, and response to immunotherapy. We note that because this language focuses on fundamental cell behavioral responses to biophysical signals, it can be applied far beyond cancer biology such as to immunology, developmental biology, tissue engineering, and related areas.

Building on the successful efforts by the community to compile unified definitions and ontologies of cell behavior^2,5,51–55^, this work represents the first major integration of those ontologies with high-throughput single cell and spatial omics data. Our grammar defines the components of ABMs based on annotated cell types and behaviors, and empowers integration of cellular phenotypes estimated from genomics data. Rules can be both knowledge-driven (e.g., based on expert statements that observe changes in cell behavior in response to chemical stimuli) or data-driven (e.g., observing changes in expression of immune cells in the tumor microenvironment). We demonstrate how the data-driven rules enable for the first time direct transfer of bulk and single-cell RNA-seq and spatial transcriptomics into ABMs for personalized predictions. Assimilating multi-omics and spatial molecular data into agent-based models can ground simulations in the real-world in such a way that ABMs can serve as an abstract digital copy of the biological system as it changes over time.

## RESULTS

Agent-based simulation frameworks^13^ like PhysiCell^44^ model individual cells as software agents with independent states (e.g., position, volume, cycle status) and processes (e.g., volume regulation, motility, secretion); see **Fig. 1A**. Each cell agent respond to stimuli (signals) in their microenvironment that effect changes in their behaviors (**Fig. 1B**). Cell hypotheses relating cell behavioral responses to signals are written in a grammar that can be translated into mathematics and executable code. (**Fig. 1C**). Hypotheses can be drawn from a variety of sources, including domain expertise, natural language processing / mining of prior literature, and analysis of transcriptomic and other data. Due to our uniform knowledge representation, all these rules can be compatibly integrated into a growing library of cell hypotheses. (**Fig. 1D**). We implement the grammar using the PhysiCell agent-based modeling framework^44^, however it can be translated to other agent-based modeling systems^45,47–50,56–58^.

**Figure 1:**
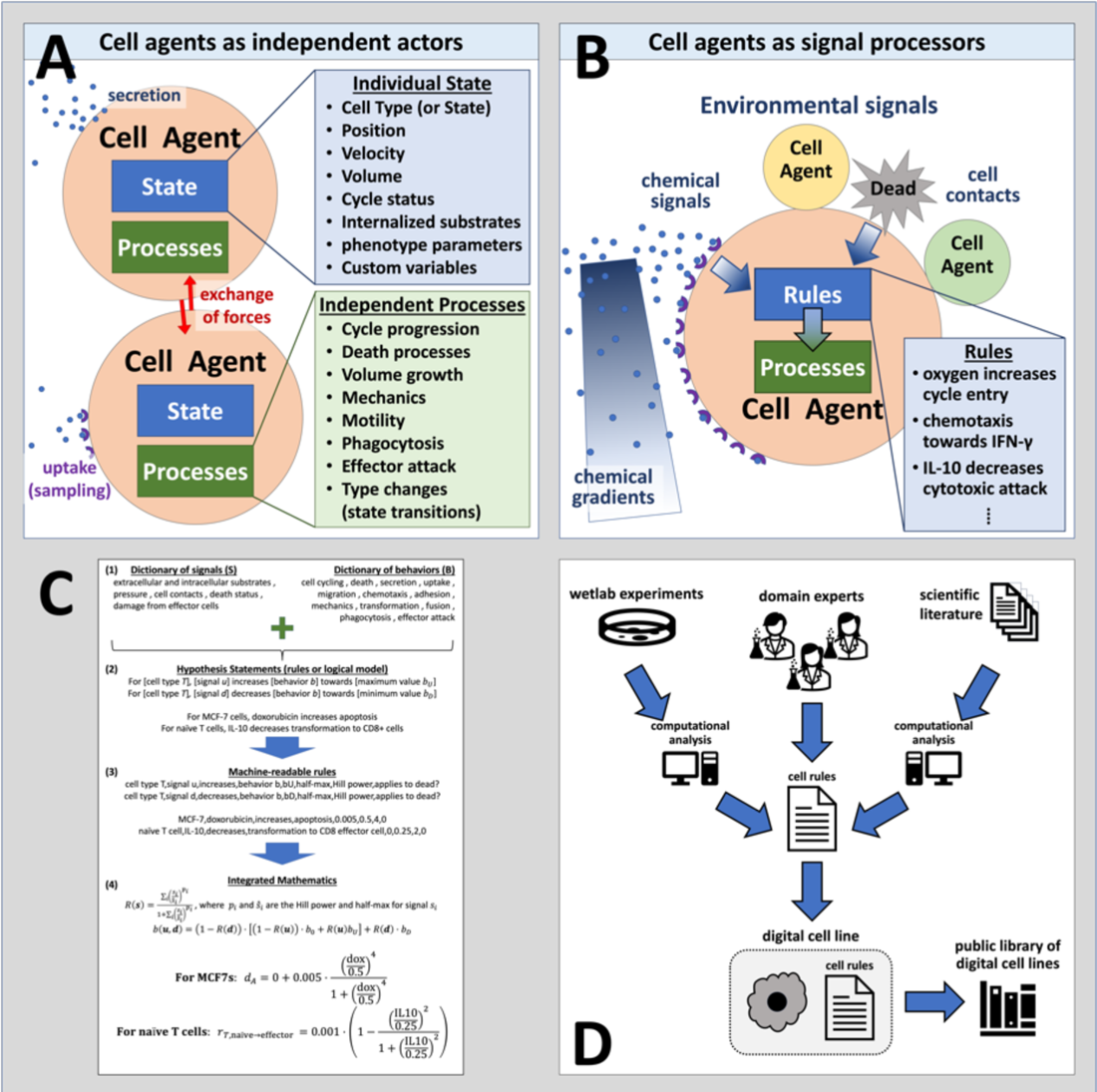
Using agent-based models to digitize cell knowledge. (A) Agent-based models simulate cells as individual objects with separate states and processes. (B) Cells agents use rules that process biophysical signals in their microenvironment—including other cells—to drive changes in their behaviors. These rules are based on our biological hypotheses. (C) The cell behavior grammar combines signals and behaviors from well-defined dictionaries (1) to create interpretable hypothesis statements (2) that can be automatically transformed into computer-readable code (3) and mathematical models (4). (D) Rules can integrate knowledge gained from novel experiments, domain expertise, and literature mining to create digital cell lines. Over time, libraries of digital cell lines accumulate, curate, and systemize our knowledge.

After describing our new hypothesis grammar, we demonstrate five progressive case studies that express complex multicellular systems behaviors using the behavior hypothesis grammar. The models represent incrementally more complex systems of interacting Eukaryotic cells, and each seeks to highlight different capabilities and features of PhysiCell and the cell hypothesis rules grammar. (**Figures 2-6**, **Tables 4-8**)

**Figure 2.**
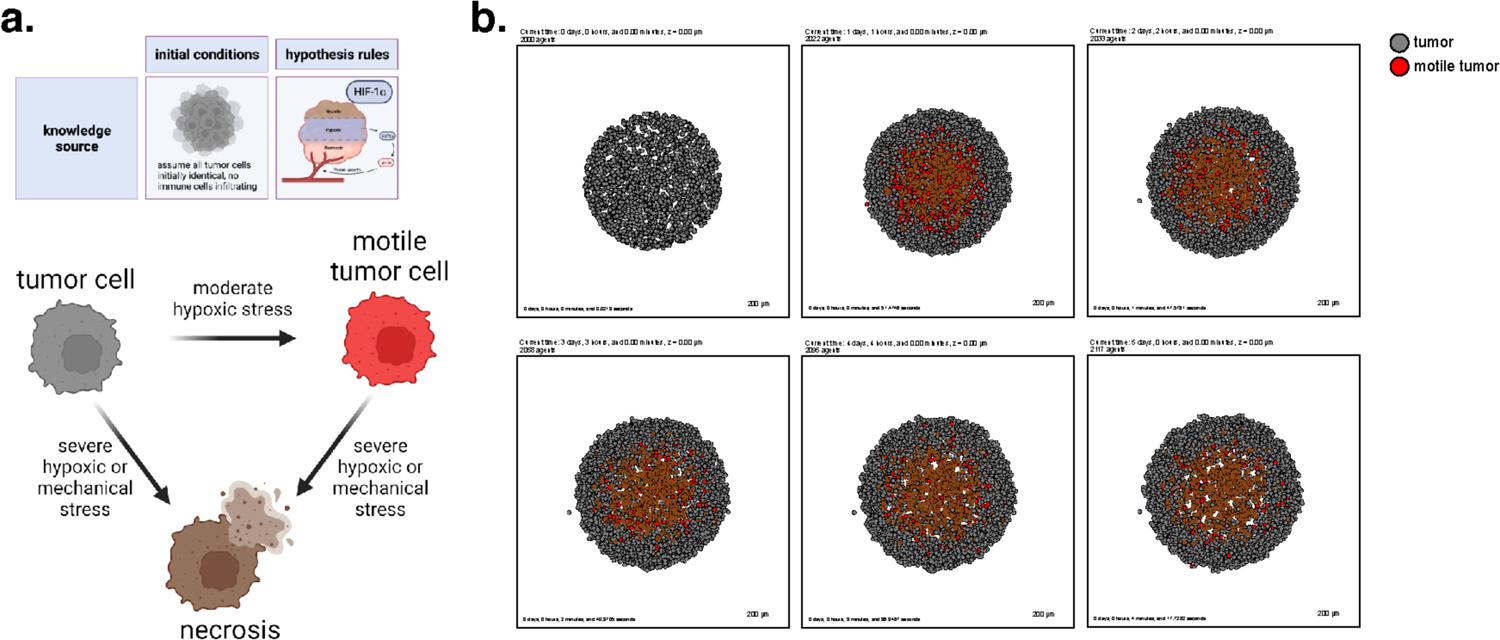
A transient, hypoxia-induced migratory phenotype induced within a homogenous tumor. (a) A cartoon showing the biology in this model and the possible cell type transitions. (b) Simulation snapshots at intervals throughout 5 days. Observe the development of a necrotic core, and the failure of motile tumor cells to reach the tumor boundary before reverting to their prior phenotype.

All five models simulate a tumor microenvironment with increasing cell diversity, including immune cells, culminating in the fifth reference model which performs a virtual clinical trial to predict the effect of combination immune-targeted therapies on tumor clearance using genomics-derived rules (**Fig. 6**). This approach is generalizable to diverse biological systems.

### A grammar encoding cell behavioral responses to extracellular signals

#### Behavioral statements

For any cell type ***T***, we construct simple statements that relate changes in a single behavior ***B*** to an exogenous signal ***S***: “In cell type ***T***, ***S*** increases/decreases ***B*** [with optional arguments].” Here ***B*** is a welldefined biophysical parameter in our dictionary of behaviors (see **Methods** and **Supplementary Information**), ***S*** is a well-defined biophysical variable in our dictionary of signals, and optional arguments further specific the mathematical behavior of the responses. For example:

In malignant epithelial cells, oxygen increases cycle entry.

In MCF-7 breast cancer cells, doxorubicin increases apoptosis.

A full description of the grammar, optional arguments, and examples can be found in the **Supplementary Information**.

#### Mathematical representation: individual rules

With clearly defined behaviors and signals and the grammar to connect them, we can uniquely map human-interpretable cell hypothesis statements onto mathematical expressions that make the grammar both human interpretable and computable. Each individual rule modulates a single behavioral parameter *b* as a function of a signal *S*. Given a response function *R*, we then mathematically represent the individual rule as a function *b*(*S*):

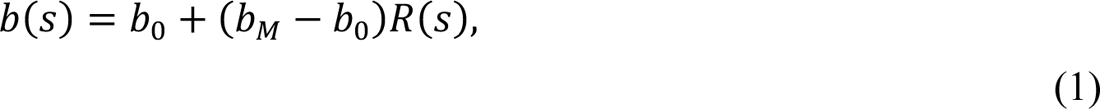

where *b*_0_is the base value of the parameter in the absence of signal, and *b*_M_ is the maximally changed value of the parameter with large signals. By default, we use sigmoidal (Hill) response functions *R*, due to their familiar use in signaling network models and pharmacodynamics, as well as their smooth variation between 0 (at no response) and 1 (at maximum response). However, capped linear response functions (varying between 0 and 1) are also possible. See **Fig. 1C** for a typical rule. Full mathematical details and additional detailed examples are available in the **Supplementary Information**.

#### Generalized mathematical representation: multiple rules

Our full mathematical formulation allows new hypotheses to be directly added to models without modifying prior hypotheses, making our framing extensible and scalable as new knowledge is acquired. Suppose that a behavior ***B*** (with corresponding behavioral parameter *b*) is controlled by multiple rules subject to promoting (up-regulating) signals *u* and inhibiting (down-regulating) signals *d*:

- *u*_1_ increases B (with half-max *u*_1_^∗^ and Hill power *p*_1_)
- *u*_2_ increases B (with half-max *u*_2_^∗^ and Hill power *p*_2_)
- …
- *u*_m_ increases B (with half-max *u*_m_^∗^ and Hill power *p*_m_)
- *d*_1_ decreases B (with half-max *d*_1_^∗^ and Hill power *q*_1_)
- *d*_2_ decreases B (with half-max *d*_2_^∗^ and Hill power *q*_2_)
- …
- *d*_n_ decreases B (with half-max *d*_m_^∗^ and Hill power *q*_n_)

Here, let *b*_M_ be the maximum value of the behavior parameter *b* (under the combined influence of the upregulating signals *u*), let *b*_!_ be its base value in the absence of signals, and let *b*_m_ be its minimum value (under the combined influence of the down-regulating signals *d*).

We define the total up response as:

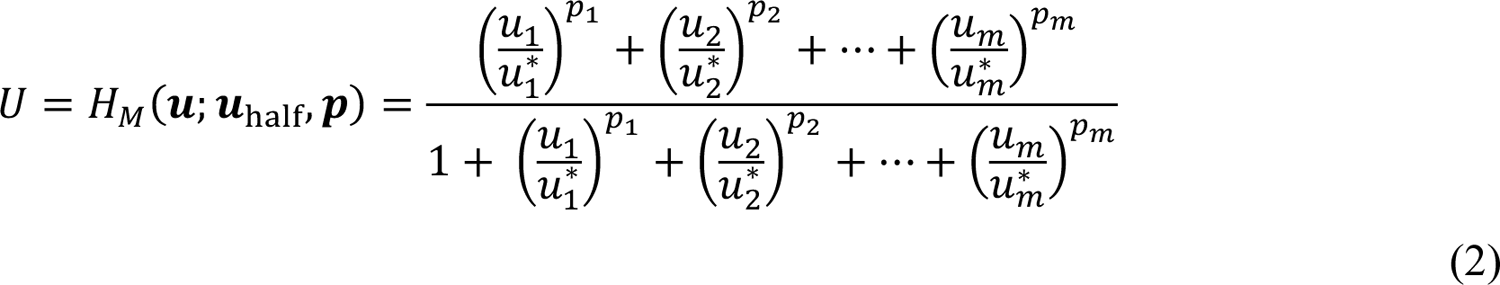

and the total down response as:

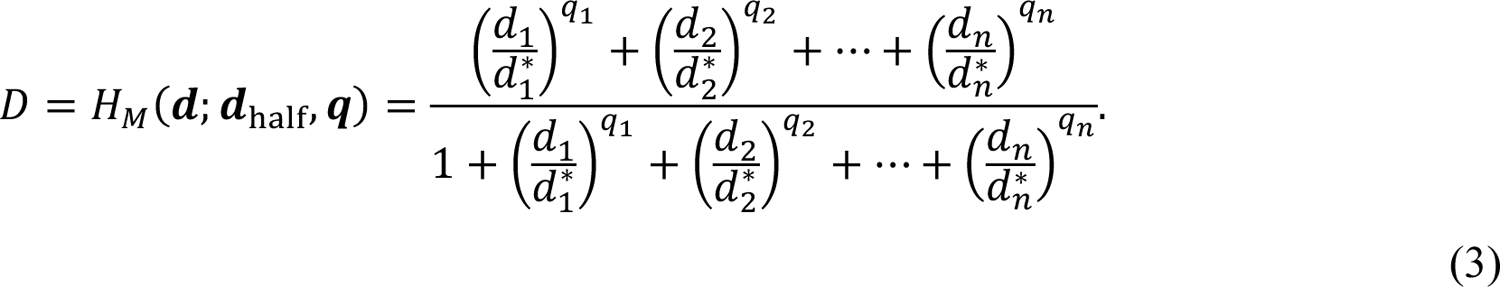

We combine the overall response of the behavioral parameter as bilinear interpolation in the nonlinear up- and down-responses *U* and *D*:

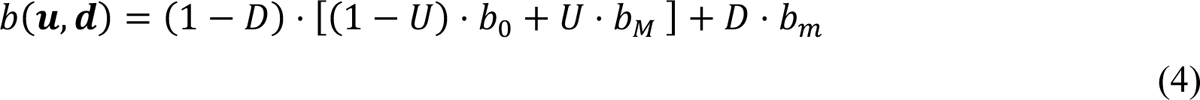

Notice that:

- In the presence of up-regulating signals only, *U* reduces to a multivariate Hill response function *H*_M_(*u*; *u*^∗^, *p*). In the presence of a single up-regulating signal *u* only, *b*(***u****,**d***) reduces to a Hill response curve *b(u)* familiar to systems biology and pharmacodynamics.
- In the presence of down-regulating signals only, *D* reduces to a multivariate Hill response function *H*_M_(*d*; *d*^∗^, *q*). In the presence of a single down-regulating signal *d* only, *b*(***u****,**d***) reduces to a Hill response curve *b(d)* familiar to systems biology and pharmacodynamics.
- Generally, the combined up-regulating signals sets a “target” value of the parameter, which can then be inhibited by the combined down-regulating signals.

Note also that adding and removing individual rules to the form does not require alteration to the remaining rules. In this release, we use multivariate Hill response functions for clarity, but mixed linear and Hill responses could be used in the future. The PhysiCell implementation of this generalized response, additional mathematical details, and expanded examples can be found in the **Supplementary Information**.

#### Automated model annotation

To help drive reproducibility, we generate and save a full description of all rules in HTML and text formats after initial parsing. Future revisions will generate these annotations in alternative formats, such as Microsoft Word tables in DOCX format and LaTeX tables.

### Example 1: Model the progression of hypoxia in a metastatic tumor

In cancer, cell growth becomes unchecked, which exhausts oxygen and nutrients in non-vascularized tumors. Modeling this resource consumption problem has provided a foundation for mathematical modeling of tumors, and this serves as the base example of tumor cell behavior in the absence of an immune response, from which our other modeling examples are built.

As a first example, we show that the grammar can express common mathematical models for oxygen-dependent tumor growth^44,59–65^, with inclusion of additional mechanobiologic feedback on cycle entry as in recent tumor growth models^59,60,66^. Following prior work^59^, we model hypoxia-induced migration, where low oxygen conditions can “reprogram” tumor cells to a transient, post-hypoxic phenotype of increased chemotactic migration, and subsequent prolonged exposure to high oxygen conditions can “revert” those cells back to a less motile phenotype. Consistent with prior modeling predictions and experimental validation^59^, these motile cells fail to exit the tumor and invade the surrounding tissue when their hypoxic adaptations do not persist in higher oxygen conditions (**Fig. 2b**, later simulation times). Note that rather than stating “low oxygen increases necrosis” and “hypoxia increases transformation into motile tumor cells”, the language uses “oxygen decreases necrosis” and “oxygen decreases transformation to motile tumor cells”, as there are currently no symbols for “no”, “lack of”, or “low.”

We used these rules to simulate 5 days of growth of a 2-D tumor in an environment of 38 mmHg oxygenation (physioxic conditions^67^), starting from 2000 viable cells seeded randomly in a virtual disk with a 400 μ*m* radius. Results are shown in **Fig. 2**.

This leads to an oxygen-poor necrotic tumor core, while hypoxic cells are disseminated throughout the tumor. Here we show a model of a transient post-hypoxic phenotype of increased chemotactic migration, where cells eventually return to their baseline phenotype upon reoxygenation. The full model is stored as **example1** in the user_projects directory of the PhysiCell git repository, and an automated model annotation is in the **Supplementary Materials**.

##### Cell Hypothesis Rules (detailed)

###### In tumor cells

oxygen increases cycle entry from 1.7e-05 towards 0.0007 with a Hill response, with half-max 21.5 and Hill power 4.

pressure decreases cycle entry from 1.7e-05 towards 0 with a Hill response, with half-max 0.25 and Hill power 3.

oxygen decreases necrosis from 0.0028 towards 0 with a Hill response, with half-max 3.75 and Hill power 8.

oxygen decreases transform to motile tumor from 0.001 towards 0 with a Hill response, with half-max 6.75 and Hill power 8.

###### In motile tumor cells

oxygen increases cycle entry from 1.7e-05 towards 0.0007 with a Hill response, with half-max 21.5 and Hill power 4.

pressure decreases cycle entry from 1.7e-05 towards 0 with a Hill response, with half-max 0.25 and Hill power 3.

oxygen decreases necrosis from 0.0028 towards 0 with a Hill response, with half-max 3.75 and Hill power 8.

oxygen increases transform to tumor from 0 towards 0.005 with a Hill response, with half-max 6.75 and Hill power 8.

### Example 2: Forecast tumor progression in pancreatic cancer from an initial state defined by spatial transcriptomics

We sought to extend our rules framework from our toy model of tumor cell growth in Example 1 to a personalized, data-driven model of tumor progression. For these models, we leveraged the rules-based framework to encode rules of cell-cell interactions and initial conditions of cells from spatial transcriptomics and single-cell RNA-seq datasets. Due to the dense stroma that is characteristic of PDAC, we sought to use our computational model to investigate the influence of cancer-associated fibroblast (CAF) content on tumor progression (**Fig. 3a**). Our previous studies of cell-cell interactions in single-cell RNA-seq analysis and organoid co-cultures demonstrated two distinct transcriptional phenotypes in neoplastic cells^68^. Fibroblast density and co-culture were associated with co-occurrence of inflammatory signaling and EMT in neoplastic cells, and attributed to cell-cell communication from integrins in fibroblasts (ITGB1) to epithelial cells. This neoplastic cell phenotype was mutually exclusive with proliferative signaling in epithelial cells. These observations led us to hypothesize that fibroblasts promote the epithelial-to-mesenchymal transition (EMT), facilitating and enabling cell invasion in PDAC. We leverage the rules-based framework to test this hypothesis in computational simulations. Our model boils down key features of interest from this transcriptional analysis into a simplified model of a pancreatic tumor microenvironment, where the major actors are epithelial and mesenchymal cells, fibroblasts, the collagen-rich ECM, and other pancreatic cells, here primarily modeled in their role as structure and scaffolding within which the other cell types interact. (Future work will extend the focus to additional processes and roles played by these cells.) The model has a simple pro-inflammatory factor which is pro-tumorigenic and which inhibits the normal mesenchymal-to-epithelial transition, while contact with fibroblasts increases the rate of EMT. In this way the fibroblasts in this model can shift the microenvironment in favor of tumorigenesis consistent with the hypothesis generated from our transcriptional signatures.

**Figure 3.**
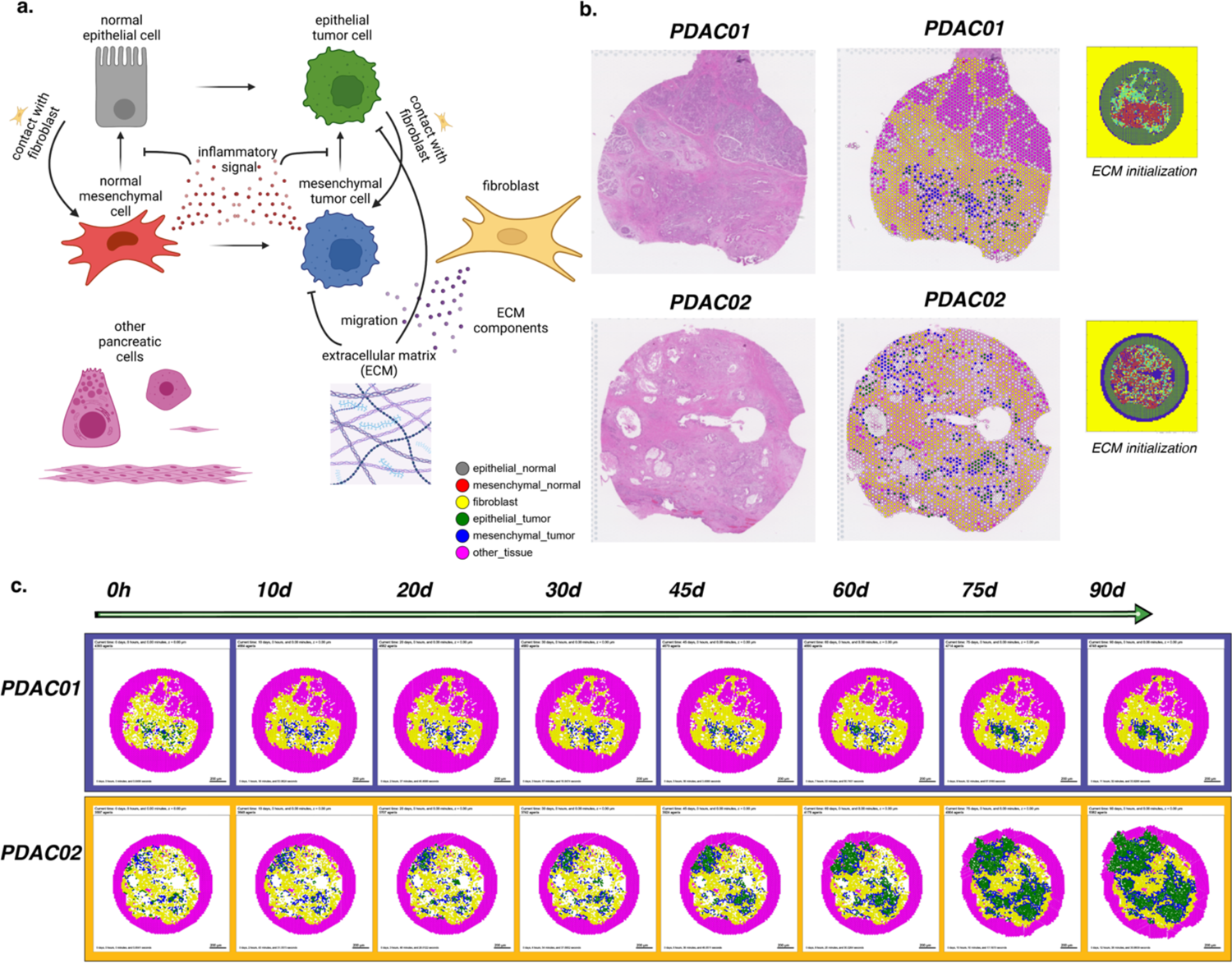
Forecasting tissue outcomes at spatial resolution in the pancreatic tumor microenvironment. (a) Diagram representing the agent types, substrates, and interactions in the model. (b) Visium spatial transcriptomics data from PDAC tissues selected for modeling and the assigned categorical spot annotations. (c) Snapshots from 90 days of simulated tumor progression in PDAC01 and PDAC02.

For this model, we initialized cell positions in a virtual tissue based on Visium spatial transcriptomic data from two pancreatic ductal adenocarcinoma (PDAC) lesions with varying fibroblast density (**Fig. 3b**)^69^. Our bioinformatics methods for three-way integration between H&E imaging data, spatial transcriptomics, and transcriptional signatures of cellular phenotypes^69^ along with pathology curation (see **Methods**) were used to categorize the epithelial cell phenotypes and fibroblasts in our model. ECM density was initialized using a heuristic from image-derived collagen and cancer-associated fibroblast annotations from the H&E imaging, using a machine learning method for tissue annotation called CODA^70^, and the bounding cells were assumed to contain a similarly dense collagen matrix, forming a niche around the known sample and abstractly reflecting the character of the solid pancreatic tissue. Other pancreatic cells in the spatial transcriptomics data were approximated as steady state (no net proliferation, death, motility, or secretion) and are assumed to be essentially inert with regards to carcinogenesis. Agents were initialized from these cellular annotations of the Visium data matrix as described in Methods. Each tumor’s development was forecasted for 90 days. Included in this count is a ring of bounding pancreatic cells. Simulation results are shown in (**Fig. 3c**). We observe a transitory state in which the tumor cells transition from mixed epithelial and mesenchymal states to become nearly uniformly mesenchymal due to interactions with the fibroblasts. Subsequently, masses of epithelial tumor cells arise in both models, consistent with extensive tumor growth and the observed transition between PDAC transcriptional subtypes during tumorigenesis and invasion^71^. An interface of mesenchymal tumor cells is maintained between the epithelial tumor cell and fibroblast cell masses. In PDAC02, rapidly dividing epithelial tumor cell clusters arise from lesions not surrounded by fibroblasts and invade the bounding pancreatic cells. In contrast, the dense, uniform fibroblast surrounding all of the lesions in PDAC01 slows invasion. The reduced rate of invasion results in smaller invasive lesions at 90 days in the PDAC01 sample compared to PDAC02. Whereas all the lesions in PDAC02 are invading bounding pancreatic cells, PDAC01 develops an epithelial tumor cell mass that is constrained from further motility by the surrounding CAFs and the dense ECM they have constructed. These computational predictions show that the hypothesized epithelial-fibroblast interactions inducing the epithelial transition between classical (epithelial-like) and basal (mesenchymal-like) pancreatic transcriptional subtypes observed in primary human pancreatic tumor progression, that return to the more epithelial-like classical subtype in metastatic sites^71^, highlighting the power of using the grammar, genomics-driven rules, and integrating spatial molecular data into dynamical modeling to contextualize and explain complex cellular interactions observed in single-cell data and organoid models from spatial.

A full copy of the automated model annotation can be found in the **Supplementary Materials**, while the code and configuration are included in the GitHub repository as **example2_pdac01** and **example2_pdac02**.

##### Cell Hypothesis Rules (detailed)

###### In epithelial_normal cells

contact with fibroblast increases transform to mesenchymal_normal from 0 towards 0.01 with a Hill response, with half-max 0.01 and Hill power 4.

###### In mesenchymal_normal cells

ecm decreases migration speed from 0 towards 0 with a Hill response, with half-max 0.5 and Hill power 4.

inflammatory_signal decreases transform to epithelial_normal from 0.01 towards 0 with a Hill response, with half-max 0.2 and Hill power 4.

###### In epithelial_tumor cells

pressure decreases cycle entry from 0.001 towards 0 with a Hill response, with half-max 1 and Hill power 4.

contact with fibroblast increases transform to mesenchymal_tumor from 0 towards 0.01 with a Hill response, with half-max 0.01 and Hill power 4.

###### In mesenchymal_tumor cells

ecm decreases migration speed from 0 towards 0 with a Hill response, with half-max 0.5 and Hill power 4.

inflammatory_signal decreases transform to epithelial_tumor from 0.01 towards 0 with a Hill response, with half-max 0.2 and Hill power 4.

### Example 3: Tumor attackers and tumor defenders

The models shown thus far allow for tumor cell growth to continue unchecked by the immune system. In these models, the only constraints on the tumor are mechanical barriers and its own size preventing the core from maintaining sufficient oxygenation. However, in reality, tumors have a microenvironment composed of immune cells that modulate tumor growth and are important components to model and key contributors to therapeutic response.

To introduce virtual immune cells, we extend our model by including CD8+ T cell agents capable of contact-mediated killing, and proand antiinflammatory factors that modulate the probability that killing will occur after a given cell contact. CD8+ T cells perform the cytotoxic killing, while macrophages switch between promoting and suppressing tumor killing (secreting pro- or anti-inflammatory factor) depending on the oxygenation in their immediate surroundings, and consistent with the literature^72–74^. Macrophages are also responsible for phagocytosing dead cells and can increase secretion of pro-inflammatory factors, which in turn attracts CD8+ T cells who home to the tumor by following this chemokine. CD8+ T cells can attack and damage malignant epithelial cells, and accumulated damage can cause tumor cell death.

We used these rules to simulate 5 days of growth of a 2-D tumor in a virtual environment of 38 mmHg oxygenation (physioxic conditions^67^), starting from 1000 viable tumor cells seeded randomly, surrounded by a ring of virtual immune cells where 50 of each non-tumor cell type are seeded. See **Fig. 4**. Through these simulations, we observe that CD8+ T cells cluster together and migrate throughout the tumor along with macrophages to accomplish tumor clearance. As such, this model shows the cooperation of the innate and adaptive immune system in the task of tumor sensing and clearance and demonstrates a simple, continuous, plastic M1-M2 axis of macrophage behavior. However, it does not account for unidirectional programs of macrophage differentiation or for the dynamics of T cell activation and clonal expansion. Since macrophages in this simulation all begin external to the tumor, they initially have the M1 phenotype which predisposes them toward an inflammatory response. Later examples further promoting immune response with immunotherapy treatment will show behavior of a tumor thoroughly infiltrated by antiinflammatory macrophages.

**Figure 4.**
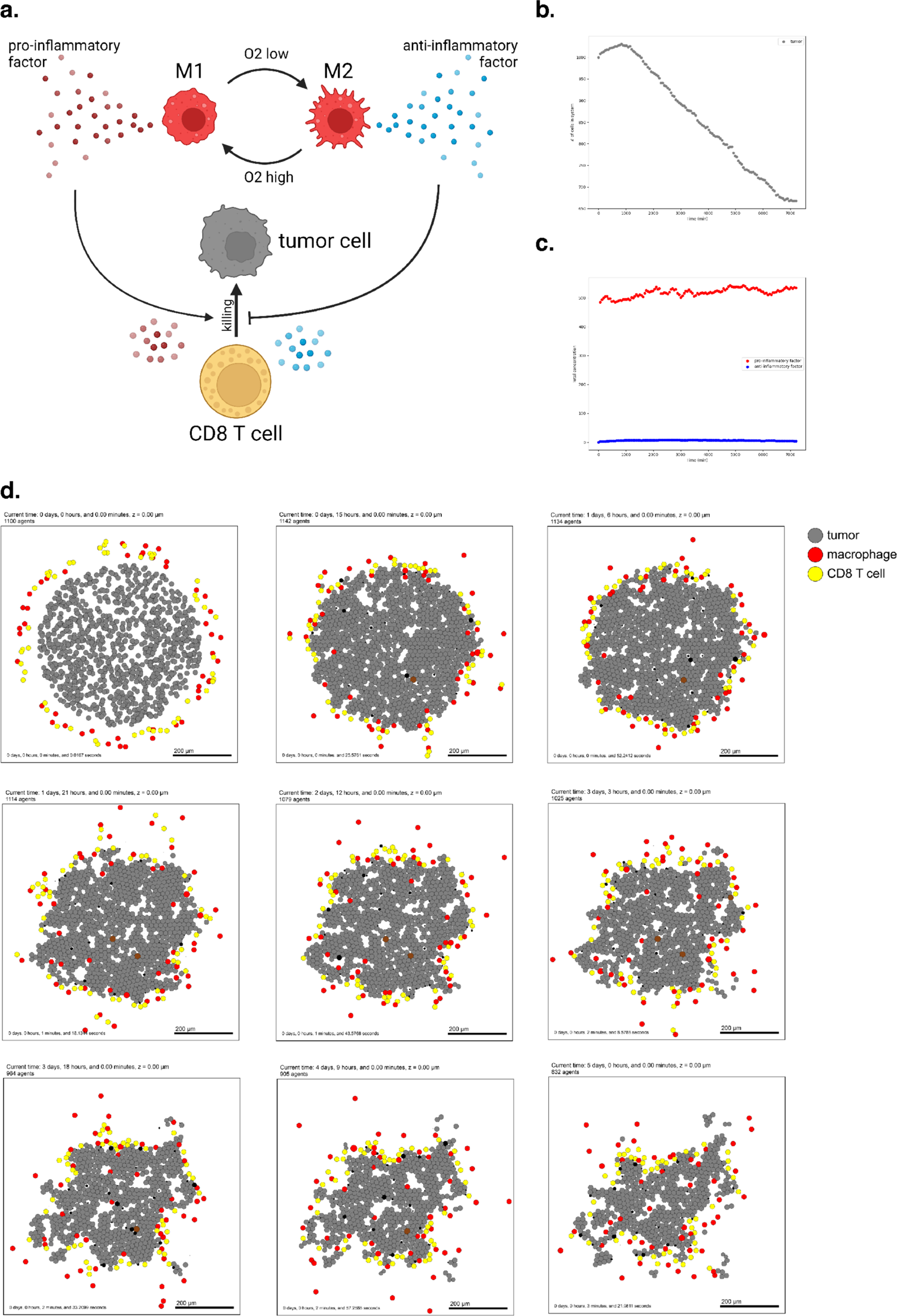
Simulating a simple antitumor immune response. (a) A schematic of cell types and substrates within the simulated tumor microenvironment (b) Tumor cell count dwindles over time as the immune response progresses. (c) Concentration of proand anti-inflammatory factor throughout the simulation. (d) Snapshots from 5 days of simulated immune response.

A full copy the automated model annotation can be found in the **Supplementary Materials**, while the code is included in the GitHub repository as **example3**.

##### Cell Hypothesis Rules (detailed)

###### In tumor cells

oxygen increases cycle entry from 0 towards 0.00072 with a Hill response, with half-max 21.5 and Hill power 4.

pressure decreases cycle entry from 0 towards 0 with a Hill response, with half-max 1 and Hill power 4.

oxygen decreases necrosis from 0.0028 towards 0 with a Hill response, with half-max 3.75 and Hill power 8.

damage increases apoptosis from 7.2e-05 towards 0.072 with a Hill response, with half-max 180 and Hill power 2.

dead increases debris secretion from 0 towards 0.017 with a Hill response, with half-max 0.1 and Hill power 10. Rule applies to dead cells.

###### In macrophage cells

oxygen increases pro-inflammatory factor secretion from 0 towards 10 with a Hill response, with half-max 5 and Hill power 4.

oxygen decreases anti-inflammatory factor secretion from 10 towards 0 with a Hill response, with half-max 5 and Hill power 4.

###### In CD8 T cell cells

anti-inflammatory factor decreases attack tumor from 0.1 towards 0 with a Hill response, with half-max 0.5 and Hill power 8.

pro-inflammatory factor increases attack tumor from 0.1 towards 1 with a Hill response, with half-max 0.5 and Hill power 8.

anti-inflammatory factor decreases migration speed from 1 towards 0 with a Hill response, with half-max 0.5 and Hill power 8.

contact with tumor decreases migration speed from 1 towards 0 with a Hill response, with half-max 0.5 and Hill power 2.

### Example 4: T cell activation, expansion, and exhaustion in a diverse tumor microenvironment

New therapies are developing to harness the immune system to attack tumor cells and are leading to unprecedented benefit in subsets of patients with immune-rich tumors^75^. In spite of this promise, many other tumor subtypes have immunosuppressive microenvironments that limit immunotherapy response. Together these observations have motivated the development of therapeutic strategies designed to promote effective tumor response by favorably modulating the tumor immune microenvironment^76^.

Literature derived from preclinical and correlatives in clinical studies indicate the impact of therapeutics on specific cell types^77–80^. However, given the complexity of the tumor immune microenvironment, it is difficult to extend this estimated effect on individual cell types to the emergent, holistic response in complex tumor tissues. Modeling provides an important tool to estimate the influence of therapy on the overall tumor ecosystem and facilitates identification of mechanisms of therapeutic response and resistance that emerge through the interactions of different cell types^75^. To explore these dynamics, we performed a PubMed search to curate rules of cell-cell interactions implicated in modulating response to diverse therapies and input them to PhysiCell using our defined grammar^81–84^. This simulation shows T cell activation in response to tumor contact.

The previous example included a model of virtual macrophages with a continuous M1-M2 axis of behavior^85^. We now show an alternative model with discrete M0, M1, and M2 states, as well as more developed maturation of naïve CD8^+^ T cells into activated CD8^+^ T cells, in the context of an early tumor-immune microenvironment^86^. The rules also include observations about the functional interactions between these cell types via IL-10 and IFNG. This model of immune invasion shows the establishment of permissive, resident macrophages simultaneously with the initiation of an adaptive response.

We used these rules to simulate 5 days of growth of a 2-D tumor in a virtual environment of 38 mmHg oxygenation (physioxic conditions^65^), starting with 1000 viable tumor cells, surrounded by a ring comprised of 200 Naive CD8^+^ T cells and 200 M0 Macrophages (**Fig. 5**). These results show a highly successful anti-tumor response in terms of T cell activation and expansion, but complete clearance of the tumor core is not achieved by the simulation endpoint. Moreover, as the simulation progresses, a population of M2-like, tumor-protective macrophages emerge, and the number of naïve cells wanes as they activate and expand into effector T cells. The M2 macrophages secrete an anti-inflammatory factor, deterring T cells and protecting the remaining tumor from complete destruction through the duration of the simulation.

**Figure 5.**
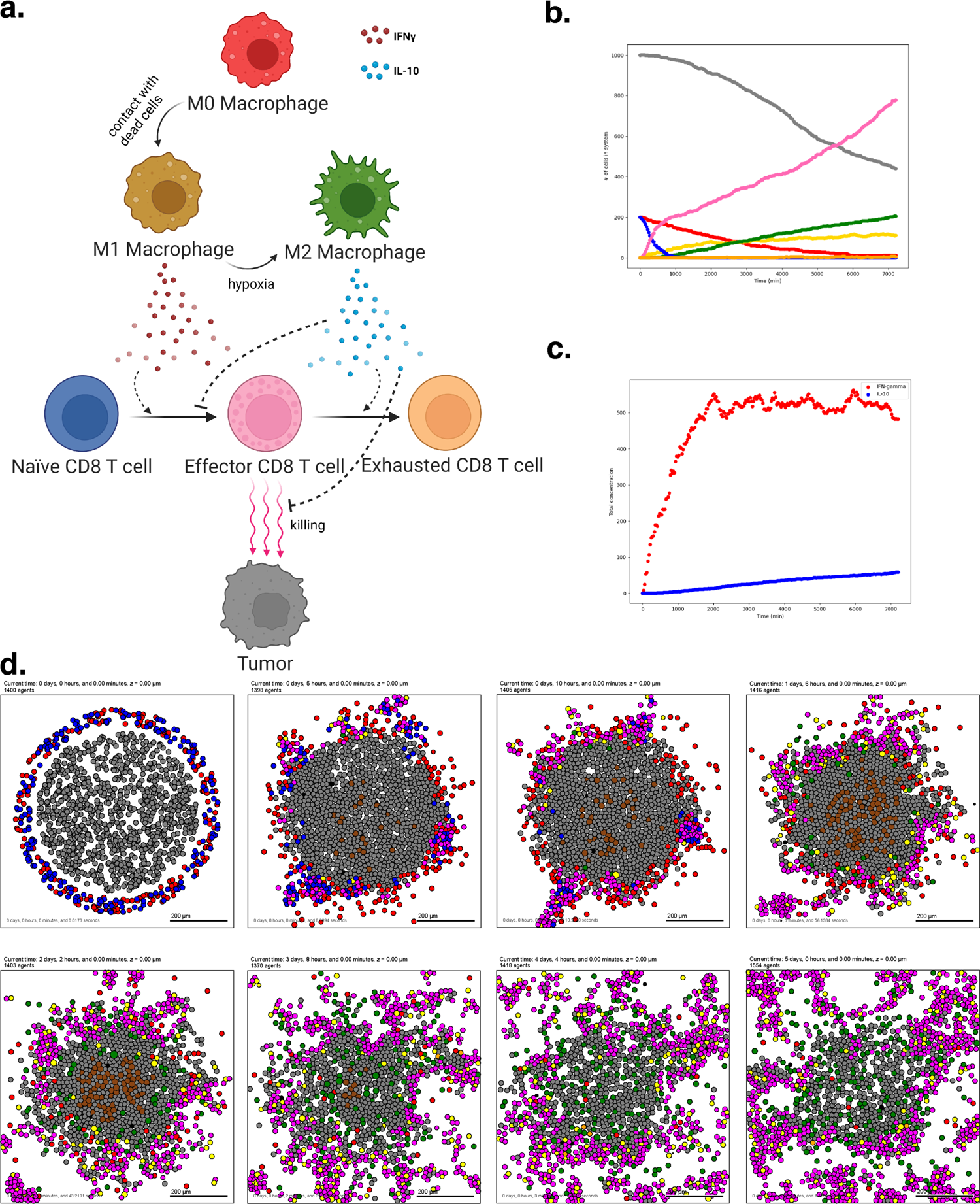
T cell activation and expansion. (a) A schematic of the cell types and states in this model. Note that macrophages can now occupy three distinct states, and transitions are unidirectional. (b) Cell counts for each cell type over time. Note the marked expansion of the CD8 T cell population and the corresponding decline in tumor burden. (c) Pro- and anti-inflammatory factor concentration throughout the simulation (d) Interval snapshots of the simulated immune response.

A full copy the automated model annotation can be found in the **Supplementary Materials**, while the code is included in the GitHub repository as **example4**.

##### Cell Hypothesis Rules (detailed)

###### In tumor cells

oxygen increases cycle entry from 0 towards 0.00072 with a Hill response, with half-max 21.5 and Hill power 4. pressure decreases cycle entry from 0 towards 0 with a Hill response, with half-max 1 and Hill power 4.

oxygen decreases necrosis from 0.0028 towards 0 with a Hill response, with half-max 3.75 and Hill power 8. damage increases apoptosis from 7.2e-05 towards 0.072 with a Hill response, with half-max 180 and Hill power 2.

dead increases debris secretion from 0 towards 0.017 with a Hill response, with half-max 0.1 and Hill power 10. Rule applies to dead cells. IFN-gamma decreases migration speed from 0.5 towards 0 with a Hill response, with half-max 0.25 and Hill power 2.

###### In M0 macrophage cells

contact with dead cell increases transform to M1 macrophage from 0 towards 0.05 with a Hill response, with half-max 0.1 and Hill power 10. contact with dead cell decreases migration speed from 1 towards 0.1 with a Hill response, with half-max 0.1 and Hill power 4.

dead increases debris secretion from 0 towards 0.017 with a Hill response, with half-max 0.1 and Hill power 10. Rule applies to dead cells.

###### In M1 macrophage cells

contact with dead cell decreases migration speed from 1 towards 0.1 with a Hill response, with half-max 0.1 and Hill power 4. oxygen decreases transform to M2 macrophage from 0.01 towards 0 with a Hill response, with half-max 5 and Hill power 4. IFN-gamma increases cycle entry from 7.2e-05 towards 0.00036 with a Hill response, with half-max 0.25 and Hill power 2. IFN-gamma increases phagocytose dead cell from 0.01 towards 0.05 with a Hill response, with half-max 0.25 and Hill power 2.

dead increases debris secretion from 0 towards 0.017 with a Hill response, with half-max 0.1 and Hill power 10. Rule applies to dead cells.

###### In M2 macrophage cells

contact with dead cell decreases migration speed from 1 towards 0.1 with a Hill response, with half-max 0.1 and Hill power 4. IFN-gamma decreases cycle entry from 7.2e-05 towards 0 with a Hill response, with half-max 0.25 and Hill power 2.

IFN-gamma increases phagocytose dead cell from 0.01 towards 0.05 with a Hill response, with half-max 0.25 and Hill power 2.

dead increases debris secretion from 0 towards 0.017 with a Hill response, with half-max 0.1 and Hill power 10. Rule applies to dead cells.

###### In naive T cell cells

IL-10 decreases transform to CD8 T cell from 0.001 towards 0 with a Hill response, with half-max 0.25 and Hill power 2.

IFN-gamma increases transform to CD8 T cell from 0.001 towards 0.01 with a Hill response, with half-max 0.25 and Hill power 2.

dead increases debris secretion from 0 towards 0.017 with a Hill response, with half-max 0.1 and Hill power 10. Rule applies to dead cells.

###### In CD8 T cell cells

IFN-gamma increases cycle entry from 7.2e-05 towards 0.00041 with a Hill response, with half-max 0.25 and Hill power 2. IL-10 decreases attack tumor from 0.01 towards 0 with a Hill response, with half-max 0.25 and Hill power 2.

IL-10 decreases migration speed from 1 towards 0.1 with a Hill response, with half-max 0.25 and Hill power 2.

contact with tumor decreases migration speed from 1 towards 0.1 with a Hill response, with half-max 0.1 and Hill power 2.

IL-10 increases transform to exhausted T cell from 0 towards 0.005 with a Hill response, with half-max 0.25 and Hill power 4.

dead increases debris secretion from 0 towards 0.017 with a Hill response, with half-max 0.1 and Hill power 10. Rule applies to dead cells.

###### In exhausted T cell cells

dead increases debris secretion from 0 towards 0.017 with a Hill response, with half-max 0.1 and Hill power 10. Rule applies to dead cells.

### Example 5: Using experimental insight to model combination immune-targeted therapies in the pancreatic cancer microenvironment

While the literature contains a wealth of prior knowledge of cellular roles in cancer, distilling cell behaviors into quantifiable networks for modeling previously required extensive manual curation. New singlecell datasets are uncovering novel cellular mechanisms of tumors in the microenvironment and their response to therapy. As we showed in example 2, data-driven cell-cell interaction rules derived from these datasets provide the opportunity to refine knowledge-driven or literature-based rules in the context of individual tumors. Integrating genomics datasets into ABMs can further personalize virtual prediction of therapeutic outcomes.

An ongoing platform neoadjuvant clinical trial^87^ is evaluating combination therapies in human pancreatic cancer to rewire the immune system through systematic, rational combination therapeutic strategies. Building on the essential tumor-immune paradigm demonstrated in preceding examples, we sought to model and understand behaviors observed from human pancreatic tumor biospecimens.

Multi-omics data from our previous neoadjuvant clinical trial of GVAX and Nivolimab demonstrated that immunotherapy activates chemokine signaling in CD4 T cells signaling to CD8 T cells to promote changes in lymphocyte chemotaxis^88^. In this example, we (computationally) simulate therapy for a cohort of patients using our behavior rules and initial cell numbers derived from the cellular proportions in our reference atlas of single-cell RNA-seq data from untreated pancreatic tumors^89^. These simulations demonstrate inter-patient heterogeneity of treatment effects, which we hypothesize resulted from differences in initial immune cell ratios in the simulated tumor microenvironment (**Fig. 6b**). Indeed, when comparing predicted tumor cell killing between baseline tissues, tumor growth correlates negatively with baseline CD8+ T cell content, and is attenuated when simulated therapy is applied (**Fig. 6c**). These results demonstrate a correlation between baseline CD8+ T cell numbers and tumor volume, consistent with our previous analysis of overall survival in the PDAC immunotherapy clinical trial^88^.

**Figure 6.**
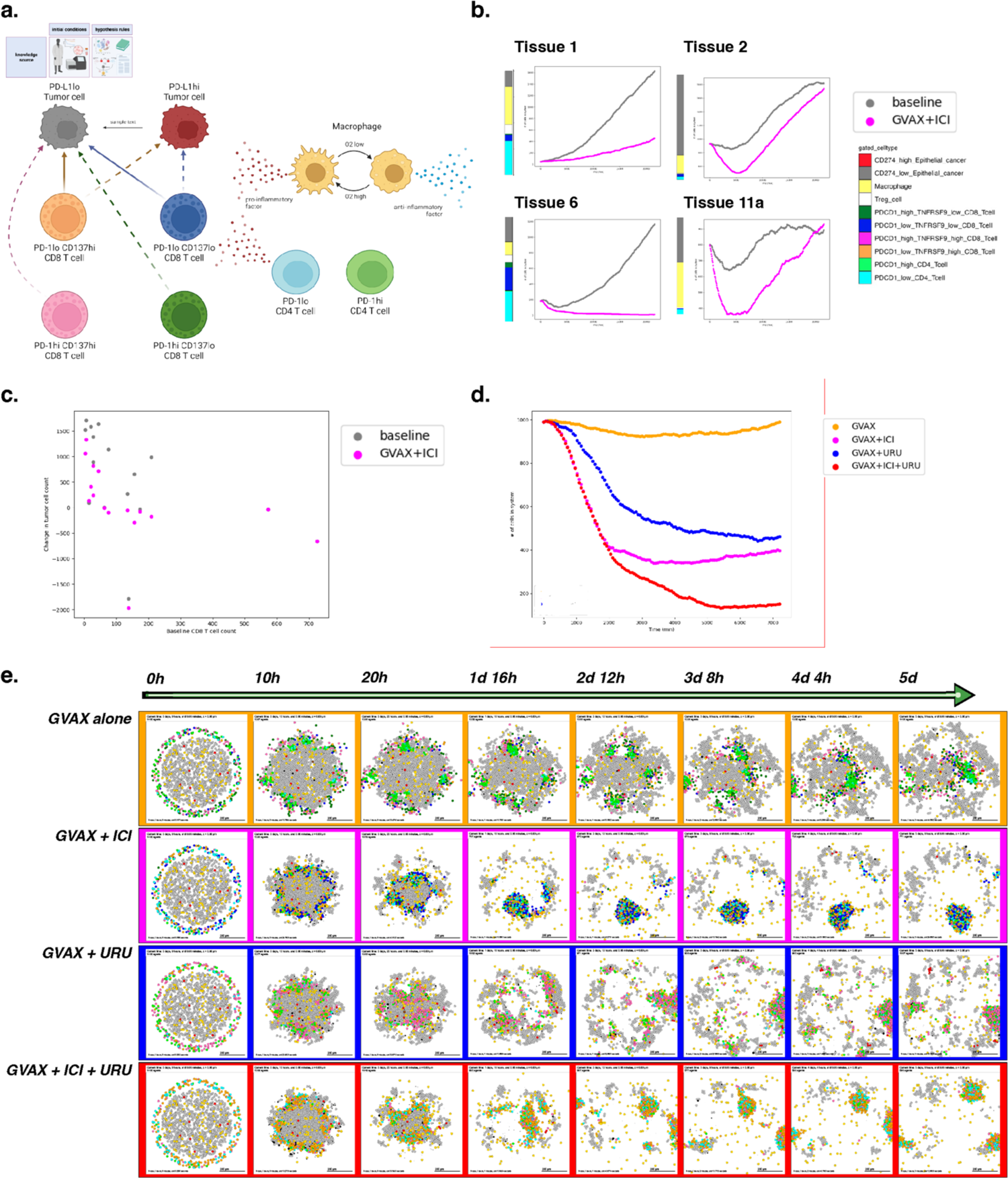
A simulation of combination immune-targeted anti-cancer therapies. (a) Schematic of cell types and states present in the model. Note differential killing ability and factor secretion between CD8 T cell subtypes. Therapy is modeled by shifting cell phenotypes according to the agonist or blocking antibodies they have received. (b) Tumor cell count over time simulated at baseline and with therapy. Four tissues were chosen to represent observed patterns of immune response across 16 total tissues. (c) Relationship between simulated tumor volume change over 15 days and baseline CD8 counts for each pancreas tissue identified from the PDAC atlas. (d) Tumor cell counts across simulated time for each therapeutic condition. Note tumor clearance following simulated triple combination therapy. (e) Snapshots over five days of simulation for four therapy conditions.

The limited responsiveness of PDAC to immunotherapy has led to new combination therapeutic strategies to further enhance T cell function. To test the mechanisms whereby different therapies can alter T cells, we next simulate monotherapy and combination therapy on a single patient’s hypothetical pancreatic microenvironment.

Our group has recently observed tumor response associated with prolonged disease-free survival through the addition of CD137 agonist therapy to a combination of an irradiated, granulocyte-macrophage colonystimulating factor (GM-CSF)–secreting, allogeneic PDAC vaccine (GVAX)^90^ and immune checkpoint inhibitor therapy^87^, consistent with the promise shown by this approach in preclinical studies. Previously, our ligand-receptor analysis of the PDAC atlas single-cell data further predicted increased interferon expression in CD137 high CD8 T cells^88^. We hypothesize that this difference in the cellular regulatory network will increase CD8+ T cells ability to infiltrate the tumor and maintain an anti-tumor state, impacting immunotherapy outcome when the proportion of CD137 high CD8 T cells is increased through the addition of an agonist therapy. We can represent these genomics-driven rules in PhysiCell to inform a model to predict and explain the impact of therapeutic combination between GVAX, Nivolumab (anti PD-1), and Urelumab (CD137 agonist) therapies.

We used our genomics observations of cell-cell interactions to motivate the rules of cellular agents in PhysiCell and built four parallel models of tumor-immune interactions. To reflect the dense immunosuppressive nature of pancreatic tumors, the models are initialized as having recruited resident suppressors with dense internal macrophages and external T cells, excluded from the tumor. As inferred in the bulk RNA-seq data, PD-1lo CD4+ T cells secrete chemokines that can attract CD8+ T cells to cause a bias in migration. Based on the single-cell RNA-seq-derived rules, CD137hi CD8+ T cells further secrete proinflammatory factor, allowing for further motility bias (i.e., more directed migration) and tumor sensing. We model treatment by including CD4+ T cells following the vaccine treatment, and further altering the proportion of PD-1lo cells to simulate Nivolumab therapy and CD137hi cells to simulate the impact of Urelumab. In contrast to example 2, fibroblasts were excluded from our model due to their underrepresentation in the reference bulk and single-cell datasets from which these rules of cell-cell interactions were derived.

In the first simulation, we model tumor-immune interactions occurring after immune priming via GVAX cancer vaccine alone; this is the first row in **Fig. 6e**. In the second, we simulate a combination of GVAX and anti-PD-1 (Nivolumab), showing increased killing but poor and uneven infiltration, with some tumor foci completely escaping immune detection; see the second row in **Fig. 6e**. In the third simulation (third row in **Fig. 6e**), we model the combination of GVAX and CD137 agonist, and shows superior tumor infiltration by T cells but inefficient killing. And in the fourth simulation (fourth row in **Fig. 6e**), we modeled the combination of GVAX, Checkpoint, and CD137 agonist, which in our model resulted in effective tumor clearance through effective infiltration and killing (**Fig. 6c**).

When we apply this approach, we observe that immune cells aggregate or “clump” together after vaccine therapy, which limits the effective infiltration of T cells throughout the tumor area and subsequent success of tumor killing. This observation is consistent with clinical observations of the formation of proximal sites of lymphoid activation (tertiary lymphoid structures or TLS) observed after neoadjuvant vaccination in PDAC^91^. Though there is clear T cell priming and expansion occurring at the TLS, presumably comprising anti-tumor neoantigen or vaccine neoantigen-specific T cells, immune infiltration into the tumor mass is limited in both our mathematical model and clinical studies. The addition of anti-PD-1 blocking antibody and CD137 agonist antibody allows T cells to effectively traffic into the tumor, leading to enhanced tumor cell killing in the triple combination in our mathematical model. Our early “toy” models of tumor-T cell interactions—which did not incorporate further immune cell communication—also predicted T cell aggregation, inconsistent infiltration, and poor response^92,93^; this further highlights the key importance of immune cell coordination across time and space for effective immunotherapeutic responses, and shows the utility of the grammar and models in elucidating these key communications.

A full copy the automated model annotation can be found in the **Supplementary Materials**, while the code is included in the GitHub repository as **example5_gvax** (case 1), **example5_gvax_ipi** (case 2), **example5_gvax_uru** (case 3), and **example5_gvax_ipi_uru** (case 4).

##### Cell Hypothesis Rules (detailed)

###### In PD-L1lo_tumor cells

oxygen increases cycle entry from 0 towards 0.00072 with a Hill response, with half-max 21.5 and Hill power 4. pressure decreases cycle entry from 0 towards 0 with a Hill response, with half-max 1 and Hill power 4.

oxygen decreases necrosis from 0.0028 towards 0 with a Hill response, with half-max 3.75 and Hill power 8. damage increases apoptosis from 7.2e-05 towards 0.072 with a Hill response, with half-max 180 and Hill power 2.

dead increases debris secretion from 0 towards 0.017 with a Hill response, with half-max 0.1 and Hill power 10. Rule applies to dead cells.

###### In PD-L1hi_tumor cells

oxygen increases cycle entry from 0 towards 0.00072 with a Hill response, with half-max 21.5 and Hill power 4. pressure decreases cycle entry from 0 towards 0 with a Hill response, with half-max 1 and Hill power 4.

oxygen decreases necrosis from 0.0028 towards 0 with a Hill response, with half-max 3.75 and Hill power 8. damage increases apoptosis from 7.2e-05 towards 0.072 with a Hill response, with half-max 180 and Hill power 2.

dead increases debris secretion from 0 towards 0.017 with a Hill response, with half-max 0.1 and Hill power 10. Rule applies to dead cells.

###### In macrophage cells

oxygen increases pro-inflammatory factor secretion from 0 towards 1 with a Hill response, with half-max 5 and Hill power 4. oxygen decreases anti-inflammatory factor secretion from 10 towards 0 with a Hill response, with half-max 5 and Hill power 4.

###### In PD-1hi_CD137lo_CD8_Tcell cells

contact with PD-L1hi_tumor decreases migration speed from 1 towards 0 with a Hill response, with half-max 0.5 and Hill power 2.

###### In PD-1lo_CD137lo_CD8_Tcell cells

anti-inflammatory factor decreases attack PD-L1hi_tumor from 1e-06 towards 0 with a Hill response, with half-max 0.5 and Hill power 8. pro-inflammatory factor increases attack PD-L1hi_tumor from 1e-06 towards 1 with a Hill response, with half-max 0.5 and Hill power 8. anti-inflammatory factor decreases attack PD-L1lo_tumor from 1e-05 towards 0 with a Hill response, with half-max 0.5 and Hill power 8. pro-inflammatory factor increases attack PD-L1lo_tumor from 1e-05 towards 1 with a Hill response, with half-max 0.5 and Hill power 8. anti-inflammatory factor decreases migration speed from 1 towards 0 with a Hill response, with half-max 0.5 and Hill power 8.

contact with PD-L1hi_tumor decreases migration speed from 1 towards 0 with a Hill response, with half-max 0.5 and Hill power 2.

###### In PD-1hi_CD137hi_CD8_Tcell cells

contact with PD-L1hi_tumor decreases migration speed from 1 towards 0 with a Hill response, with half-max 0.5 and Hill power 2.

###### In PD-1lo_CD137hi_CD8_Tcell cells

anti-inflammatory factor decreases attack PD-L1hi_tumor from 1e-06 towards 0 with a Hill response, with half-max 0.5 and Hill power 8. pro-inflammatory factor increases attack PD-L1hi_tumor from 1e-06 towards 1 with a Hill response, with half-max 0.5 and Hill power 8. anti-inflammatory factor decreases attack PD-L1lo_tumor from 1e-05 towards 0 with a Hill response, with half-max 0.5 and Hill power 8. pro-inflammatory factor increases attack PD-L1lo_tumor from 1e-05 towards 1 with a Hill response, with half-max 0.5 and Hill power 8. anti-inflammatory factor decreases migration speed from 1 towards 0 with a Hill response, with half-max 0.5 and Hill power 8.

contact with PD-L1hi_tumor decreases migration speed from 1 towards 0 with a Hill response, with half-max 0.5 and Hill power 2.

###### In PD-1hi_CD4_Tcell cells

anti-inflammatory factor decreases migration speed from 1 towards 0 with a Hill response, with half-max 0.5 and Hill power 8.

###### In PD-1lo_CD4_Tcell cells

anti-inflammatory factor decreases migration speed from 1 towards 0 with a Hill response, with half-max 0.5 and Hill power 8.

## DISCUSSION

The real-world limitations inherent to human-focused research, especially for clinical pathology samples and population- or trial-level data, do not exist *in silico*. Thus, scientists are turning to computational models to guide and supplement lab experiments. For example, the NCI digital twins initiative aims to develop models of patient tumors to predict which therapies will most benefit each individual^34,36^. Mathematical models allow investigators to simulate many replicates of their system’s behavior under different sets of conditions. The ability to perform large numbers of replicates and numerous iterations cheaply and easily maximizes the chance of capturing extremely rare critical events, something key to those who study cancer etiology. An agent-based model is an abstraction that can be run thousands or millions of times, and whose parameters and in-built hypotheses are all readily modifiable by the user. Here, we introduced a new conceptual framing (a grammar) for specifying cell behavior hypotheses, which can systemize and facilitate our thinking of how cells interact to drive tissue ecosystems. The grammar made it possible to introduce new capabilities in the PhysiCell Agent Based Modeling (ABM) framework that simplify the workflow for investigators to build a simulation of their experimental system.

The hypothesis grammar gives the ability to encode complex behaviors and responses to signals into model agents via a single line of human readable text. Previously, custom hand-written code and a high level of technical knowledge were required to implement even basic models. Now, a scientist can easily create, specify, and modify cell behaviors without writing code or hand-edited markup languages, particularly when used in combination of graphical and cloud-based modeling frameworks^94^. In this implementation, it is simple to modulate and apply behaviors to different agents in the system and to build hypotheses into the model. Any multi-cell model system can be translated into an ABM. Further, PhysiCell is open-source, built and maintained by a large community of mathematicians, statisticians, and scientists. From their efforts and from the user community, a vast amount of knowledge is encoded in the system at baseline; however, everything is completely customizable, extensible, and modifiable. Even when hypotheses are not fully executed in simulations, the cell behavior grammar affords an opportunity to systematically collect, annotate, curate, and grow our knowledge of behaviors of many cell types.

Additionally, we have added the ability to directly translate parameters from spatial transcriptomics data— namely spatial location and annotated identity—to an ABM, and therefore use spatial data to initialize an ABM directly. Thus, models can now exactly match the tissue structure and transcriptional profile of samples directly. Spatial relationships between cells matter immensely and spatial relationships can have a great impact on simulated (and real) outcomes. This strong dependence of many cancer systems and ABM trajectories on initial conditions can complicate model inquiry and impact critical system behaviors and model parameters obtained through inference; by leveraging robust single-cell spatial transcriptomics tissue profiles as initial conditions in the digital modeling stage, the hypothesis-driven rules modeling paradigm is grounded in precise referential data but also offers a path to both stronger model inquiry and more confident mathematical inference. These models nonetheless still require annotation of a finite number of agents identified in spatial molecular data, often annotating cells into broad phenotypes and abstracting cellular subtypes. Future work must evaluate the sensitivity of models to the granularity of cellular phenotypes in these high-throughput datasets and accurate inference of parameters in the resulting higher-dimensional models. Moreover, future work must facilitate simple and robust identification of cell types and model parameterization, starting by identifying which parameters are most critical, and curating best parameter estimates for community reuse.

We demonstrate a variety of models built using the hypothesis grammar, and two also informed by genomics-derived rules, which we hope will serve as helpful references for users. These examples cover multicellular behavior, demonstrated through the case studies of carcinogenesis and immune response to tumor growth. In some cases, all agents follow the same rules and their fate is decided by the actions of those around them. In other cases, cell agents are acting at cross purposes and actively seeking to outcompete, evade, or hunt and kill each other. We model immune processes such as macrophage plasticity, T cell activation and expansion, antigen recognition, and inflammation. All of these examples are available and can be re-run on any user machine within minutes. While these provide biologically and clinically relevant models to test hypotheses, we note that future work must determine the sensitivity of models to cellular resolution, initial conditions, and model parameters. Metrics to benchmark mathematical models both qualitatively and quantitatively are essential to fully leverage these models to predict personalized tumor conditions and to empower virtual clinical trials.

In future work, we hope to further refine the hypothesis grammar to expand its usability and flexibility. We are considering incorporating the ability to specify “wild card” rules (*) and other special cases using regular expression-type syntax, as well as negation symbols (e.g., “low oxygen” or “hypoxia”) that can simplify the examples presented in this paper. However, we first wish to consult with the wider user community to receive input and feedback on such extensions. Moreover, emerging large language models (LLMs) such as Chat-GPT may facilitate “translation” of familiar language (e.g., “fibrosis”) into the smaller set of symbols in the current grammar. The language currently treats all statements as independent (inclusive OR), but we may need additional language operators to signify relationships between rules such

as AND or REQUIRES. Other generalizations and improvements to the forms of response curves, consensus process models, and default parameter values are likely to emerge from widespread community use, feedback, and discussion. We envision formal community governance from key stakeholders including experimentalists, clinicians, bioinformaticians, mathematicians, and software developers. We also envision that the hypothesis grammar could become a common language to unite current non-compatible agent-based and multicellular simulation platforms, allowing greater reproducibility.

Additionally, we aspire to establish a repository to collect and curate biological hypothesis statements grouped as digital cell lines^52^, enabling users to contribute and share cell behavior statements for future reuse in other models of the same system. This repository can serve as a valuable resource for scientists to draw upon when digitizing their system. Through collaboration and collective contributions, we can foster the growth, expansion, and maintenance of this repository. Community discussion, curated validation data, better leverage of high-throughput computing resources, and uncertainty quantification analyses techniques should lead to better understanding of the reliability and sensitivity of model parameters. We also hope to achieve fuller harmonization with other omics data analysis software and high-throughput data types as we move towards integrating data directly into agent-based models. By achieving this integration, we can enhance the model’s capability to incorporate relevant biological information and improve its alignment with the broader computational biology field.

This language framework will be useful to those seeking to build models of multicellular systems, and we are excited to continue to move toward fuller biological completeness and more complete integration with omics data, to increasingly define agent behavior in an automated and data-driven fashion. These advancements expand the functionality, usability, and compatibility of our approach, empowering interdisciplinary researchers in their computational or systems biology endeavors. These advancements expand the functionality, usability, and compatibility of our approach, empowering researchers across disciplines to unlock the full potential of their single-cell data. Armed with this conceptual framing and tools, they can extrapolate beyond single-cell characterizations to predict tissue dynamics, and ultimately perform virtual experiments to steer tissues from disease to health.

## ACKNOWLEDGEMENTS

Thanks to Laura Wood and Jennifer Elisseeff for feedback on this study.

## METHODS

### PhysiCell agent-based modeling framework

PhysiCell^44^ is an open source, agent-based modeling framework written in C++ that can run on a broad variety of desktop platforms, in the cloud^95^, and on high performance computing resources^92,93,96^. PhysiCell simulates each cell as an agent with lattice-free position and volume, individual birth and death rates, and motion driven by the balance of mechanical forces and biased random migration. In more recent versions of PhysiCell, agents can also interact with built-in models of phagocytosis, effector attack, fusion, and elastic cell-cell adhesion. PhysiCell is coupled to a reaction-diffusion solver (BioFVM^97^) that models secretion and uptake (consumption) of diffusible factors by individual cell agents at their individual positions, as well as diffusion and decay of these substrates through extracellular spaces. PhysiCell bundles its key cell behavioral parameters as a *phenotype object* for simpler representation. Modelers simulate biological hypotheses by writing custom C++ functions that dynamically vary the cell agent’s phenotype parameters based on conditions at the cell’s position, such as contact with other cells, mechanical pressure, and concentrations and gradients of signaling factors.

### Cell behaviors

To build this grammar, we require clear abstractions of key cell behaviors that frequently occur in multicellular observations and corresponding reference models. In this context, a ***cell behavior*** is a cell-scale process, such as cycling, death, or phagocytosis. Generally, each behavior can be represented by a small number of continuous phenotypic parameters, describing the rate, magnitude, or frequency of the behavior. In earlier work, Sluka et al. developed the Cell Behavior Ontology (CBO)^51^ as a controlled vocabulary of individual cell behaviors. More recently, we worked with a multidisciplinary coalition to extend and structure behaviors from the CBO and other sources into MultiCellDS^52^ (multicellular data standard). In particular, this work defined a *behavioral cell phenotype* that collects of biophysical characterizations of a cell’s behavior, organized hierarchically by function: cycling, death, volume, mechanics, secretion (including uptake), and motility. Since releasing MultiCellDS as a preprint, we have tested this approach to cell behavior through a variety of agent-based simulation and modeling projects^44,59,60,92,93,98–103^. Based upon recent immunologic modeling work^99–102^, we extended phenotype to include cell-cell interactions (phagocytosis, effector attack, and fusion), as well as transformations between cell types (e.g., differentiation, transdifferentiation, and other state changes that persist even when exogenous signals are removed).

The grammar’s supported cell behaviors and key biophysical parameters are summarized in **Table 1**. See the **Supplementary Information** for a full description of these cell behaviors, including reference model implementation details in the PhysiCell framework.

**TABLE 1.** Key cell behaviors included in the grammar, with a summary of controllable behavior parameters.

| Behavior | Main controllable behavior parameters |
| --- | --- |
| Cycling | Exit rates from each cycle phase |
| Death | Apoptotic (controlled / non-inflammatory) death rate<br>Necrotic (uncontrolled / inflammatory) death rate |
| Secretion and Uptake | Secretion rates and targets (for each diffusible extracellular substrate)<br>Uptake (consumption) rates (for each diffusible extracellular substrate)<br>Generalized net export rates (for each diffusible extracellular substrate) |
| Migration and Chemotaxis | Migration speed, bias, and persistence time<br>Chemotactic sensitivities (for each diffusible extracellular substrate) |
| Cell-cell adhesion | Interaction distances, strength<br>Attachment and detachment rates<br>Maximum number of cellular adhesions<br>Adhesion affinities (to each cell type) |
| Resistance to deformation | strength of cell repulsion |
| Transformation (changing cell type) | Rate of changing to each cell type |
| Fusion | Rate of fusing with each cell type |
| Phagocytosis<br>(predation or ingestion) | Rate of phagocytosing dead cells<br>Rate of phagocytosing each live cell type |
| (Effector) attack | Attack rates (for each cell type)<br>Rate of damaging attacked cells<br>Immunogenicity (to each potential attacking cell type) |
| Other | User-defined custom cell behavior parameters |

### Signals

Signals are (typically exogeneous but sometimes internal) stimuli or information that can be interpreted by a cell to drive behavioral or state changes. In the context of mathematical modeling, signals are inputs to constitutive laws or agent rules. We broadly surveyed mathematical and biological models from cancer biology^41,63,104–113^, tissue morphogenesis^104,114–118^, immunology^41,99–101,119,120^, and microbial ecosystems^121,122^, to generalize classes of inputs to cell behavioral rules, generally including chemical factors, mechanical cues, cell volume (e.g., for volume-based cycle checkpoints), physical contact with cells, live/dead status, current simulation time (for use in triggering events), and accumulated damage (e.g., from effector attack^123–125^). The signals are summarized in **Table 2**. See the **Supplementary Information** for a full description.

**Table 2.** A summary of the signals that can be used to drive behavioral or state changes in cells.

| Signal types symbol | accessible variables |
| --- | --- |
| Diffusible chemical substrates | extracellular concentration and gradient (for each diffusible substrate)<br>total internalized substrate (for each diffusible substrate) |
| Cell mechanics/physics | total cell volume, mechanical pressure |
| Contact | Total numbers of live and dead cells in contact<br>Total number of each live cell type in contact, contact with basement membrane |
| Effector attack | Total cell damage and accumulated attack time by effector cells |
| Death | Status as dead. Finer-grained status as apoptotic or necrotic |
| Other | Current elapsed simulation time, access to user-defined parameters |

### Coordinate transforms and spatial transcriptomics integration

To simulate cellular profiles based on initial conditions defined from VISIUM single-cell spatial transcriptomics data in PhysiCell, we interpret a given tissue profile *A*—a collection of the form (*x*_i,_*y*_i_) ∈ ℝ^+^ arranged in a triangular lattice with uniform spacing d = 71.000 μ*m*—as an (*n* × 2) (*n* × 2) matrix.

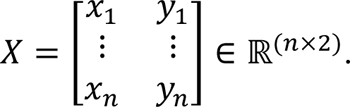

Choosing a prospective center location *X*_0_= (*x*_0_, *y*_0_) ∈ ℝ^2^ and using a hypothesized average cell diameter *d*_f_ in the tissue, we apply a linear transformation *T*_(x0,df)_ on *X* to produce a scaled cell coordinate matrix *X*_f_ = *T*_(x0,df)_ (*X*) describing a digital tissue with center (0,0) = *T*_(x0,df)_ (*X*_!_) and ideal lattice spacing on the order of *d*_3_ to encode cell signals and behaviors in the PhysiCell stage at scale.

#### Example

To generate the initial cell coordinate matrix *X*_3_used in Example 2, we start with a collection *A* and the corresponding matrix *X* ∈ ℝ(–×+)(as above) of coordinates of the form B*x*_.,_*y*.E ∈ ℝ^+^ from the spatial transcriptomics data, which contain a subset ℋ ⊂ *A* of *n*_6_ coordinates of agents of type other_cells in the main tumor mass. We compute a shift

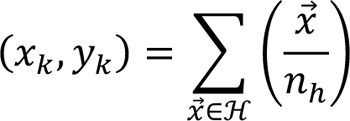

representing the coordinate-wise center of the main tissue region and the corresponding matrix translation

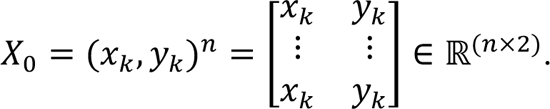

The transformed coordinate matrix *X*_f_ then takes the form

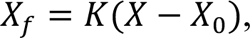

where *K* is the (*n* × *n*) scaling matrix of the form

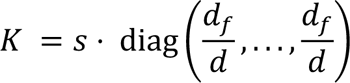

determined by a global spacing bias *S* ≈ 0.97, hypothesized average cell diameter *d*_3_ = 16.825 μ*m*, and VISIUM spacing *d* = 71.000 μ*m* from above.

### Spatial transcriptomics data for example 2

Two resected pancreatic lesions were subjected to the commercial Visium spatial transcriptomics sequencing FFPE protocol. Slides were stained with H&E and imaged prior to RNA extraction, and image analysis was performed in parallel with transcriptomic analysis. An artificial intelligence method for annotation of pancreatic tumor tissue regions called CODA^70^ was used to annotate acinar cells, islet cells, smooth muscle cells, and the distribution of collagen. This method was also used to distinguish normal ductal, neoplastic, and tumor cells from the H&E imaging, which were further visually confirmed by a pathologist (E.D.T.). Spots with greater than 70% purity of ductal cells were further annotated to assign agent types for the associated tumor and normal cells in each spot. For this annotation, we used our transfer learning method ProjectR^126^ version 1.8.0 to distinguish proliferative signaling (modeled as an epithelial phenotype) from co-occurrence of EMT and inflammatory signaling (modeled as the mesenchymal phenotype) as defined in CoGAPS non-negative matrix factorization analysis of our reference scRNA-seq atlas of PDAC tumors using methods described previously^69,89,127^. To locate fibroblasts, Seurat version 4.1.0 was used to compute module scores from a pan-CAF gene signature as described previously^69^. Two python scripts were used to translate from annotated ST data into a PhysiCell-readable csv file containing coordinates and categorical cell types available in the github repository as extract_visium_coordinates.py and pull_physicell_init.py. Submission of spatial transcriptomics data to dbGAP is in progress^128,129^.

### Single-cell RNA-seq PDAC atlas data as a reference dataset for example 5

This example uses our PDAC single-cell RNA-seq atlas data to further define immune cell subtypes in reference tumors ^130,131^. Cell-cell communication analysis was performed using the Domino package^132–134^. To determine the by-tissue cell counts, the single-cell RNA sequencing data was preprocessed, clustered, and annotated using the Seurat R package^135^. Cell identity clusters of interest (“Activated_CD4”, “B cell”, “CD4”, “CD8”, “Effector_CD8”, “Epithelial_cancer”, “Macrophage”, “Mast”, “Neutrophil”, “NK/CTL”, “T cell”, “Treg cell”) were then thresholded based on median normalized expression of genes of interest (here CD274/PD-L1, PDCD1/PD-1, TNFRSF9/CD137) and the number of cells falling into lo/hi categories were reported as described previously^88^. These cell numbers were then used to initialize the PhysiCell model in example 5, while Domino analysis informed the model rules. Whereas the cell phenotypes from the single-cell PDAC atlas data for example 2 are based on data from GSE155698^131^ and CRA001160, example 5 uses only the former dataset due the immune enrichment of this sample cohort^130^.

## Code availability

PhysiCell Version 1.12.0^136^ and later includes a full reference implementation of the grammar and grammar-based simulation modeling. The specific modeling examples presented in the results section presented below are available on GitHub at https://github.com/physicell-models/grammar_samples. To get a list of all the example models:

`make list-user-projects`

To load and compile an example named **myproject**, use:

`make load PROJ=myproject && make`

Similar to our prior work to create cloud-based training materials^98^ and cloud-based model dissemination^95^, and inspired by other recent advances on “zero-install” models^137^, we have created a cloud-based version^94,138^ of PhysiCell based on the nanoHUB platform^139^. This cloud implementation allows scientists to create, execute, visualize, and explore grammar-based models interactively in a web browser, without need for programming expertise or software setup. (See documentation and training materials in the supplementary information). The cloud-hosted model is available at https://nanohub.org/tools/pcstudio. Alternatively, scientists can download the latest release of the PhysiCell Studio^94^ desktop application at https://github.com/PhysiCell-Tools/PhysiCell-Studio/releases. Assuming an executable model has been compiled, the Studio allows interactive creation and editing of rules (**Fig. 7**), running a simulation, and visualizing results. Refer to the Studio user guide at https://github.com/PhysiCell-Tools/Studio-Guide/blob/main/README.md for more information.

**Figure 7.**
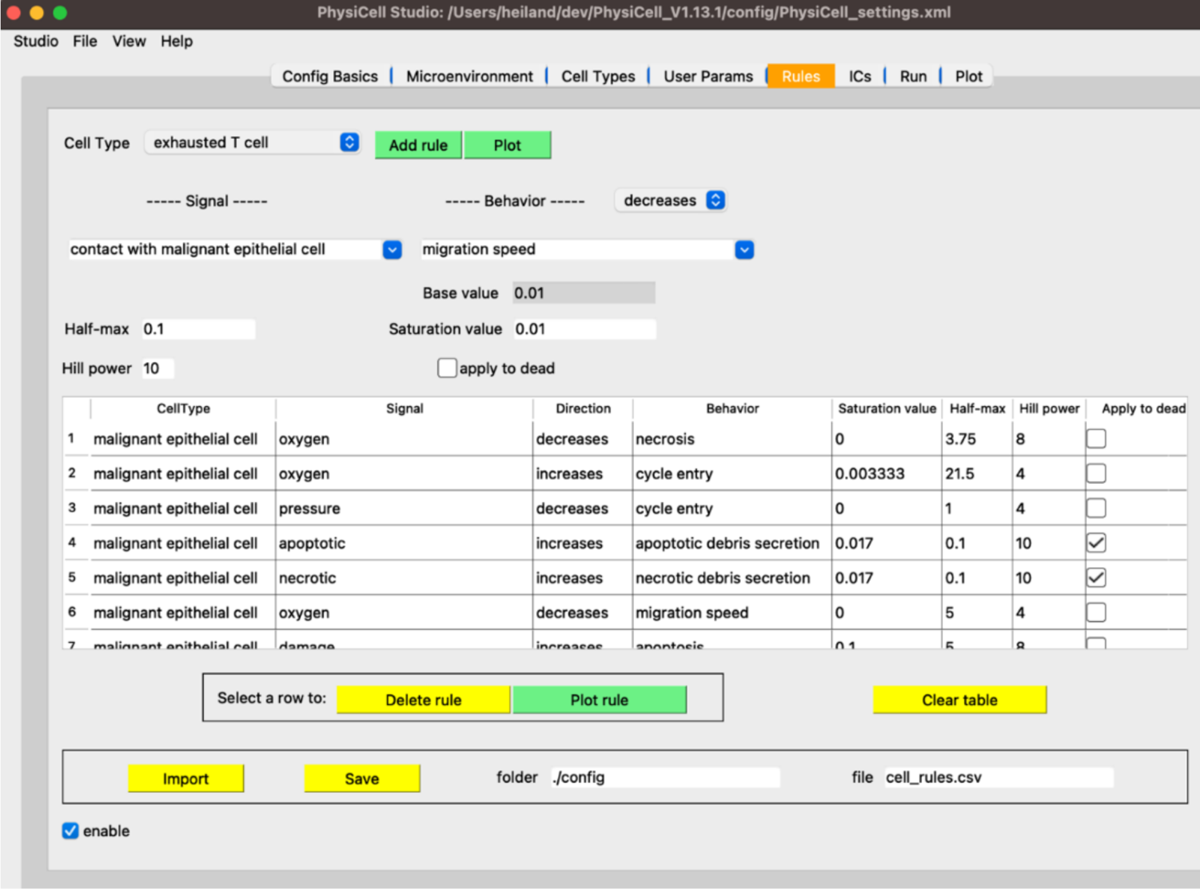
The Rules tab in PhysiCell Studio. Rules can be created and edited interactively. The choices available in Signals and Behaviors are dynamically updated based on the microenvironment and cell types.

## Inclusion and diversity

We support inclusive, diverse, and equitable conduct of research.

## References

1. Regev, A., Teichmann, S.A., Lander, E.S., Amit, I., Benoist, C., Birney, E., Bodenmiller, B., Campbell, P., Carninci, P., Clatworthy, M., et al. (2017). The Human Cell Atlas. Elife 6. 10.7554/eLife.27041.

2. HuBMAP Consortium (2019). The human body at cellular resolution: the NIH Human Biomolecular Atlas Program. Nature 574, 187–192. 10.1038/s41586-019-1629-x.

3. Rozenblatt-Rosen, O., Regev, A., Oberdoerffer, P., Nawy, T., Hupalowska, A., Rood, J.E., Ashenberg, O., Cerami, E., Coffey, R.J., Demir, E., et al. (2020). The Human Tumor Atlas Network: Charting Tumor Transitions across Space and Time at Single-Cell Resolution. Cell 181, 236–249. 10.1016/j.cell.2020.03.053.

4. Uhlen, M., Fagerberg, L., Hallstrom, B.M., Lindskog, C., Oksvold, P., Mardinoglu, A., Sivertsson, A., Kampf, C., Sjostedt, E., Asplund, A., et al. (2015). Proteomics. Tissue-based map of the human proteome. Science 347, 1260419. 10.1126/science.1260419.

5. Stein-O’Brien, G.L., Le, D.T., Jaffee, E.M., Fertig, E.J., and Zaidi, N. (2023). Converging on a Cure: The Roads to Predictive Immunotherapy. Cancer Discov 13, 1053–1057. 10.1158/2159-8290.CD-23-0277.

6. Saelens, W., Cannoodt, R., Todorov, H., and Saeys, Y. (2019). A comparison of single-cell trajectory inference methods. Nat Biotechnol 37, 547–554. 10.1038/s41587-019-0071-9.

7. Fertig, E.J., Jaffee, E.M., Macklin, P., Stearns, V., and Wang, C. (2021). Forecasting cancer: from precision to predictive medicine. Med (N Y) 2, 1004–1010. 10.1016/j.medj.2021.08.007.

8. Ding, J., Sharon, N., and Bar-Joseph, Z. (2022). Temporal modelling using single-cell transcriptomics. Nat Rev Genet 23, 355–368. 10.1038/s41576-021-00444-7.

9. Qiu, X., Zhang, Y., Martin-Rufino, J.D., Weng, C., Hosseinzadeh, S., Yang, D., Pogson, A.N., Hein, M.Y., Hoi Joseph Min, K., Wang, L., et al. (2022). Mapping transcriptomic vector fields of single cells. Cell 185, 690–711 e645. 10.1016/j.cell.2021.12.045.

10. Erbe, R., Stein-O’Brien, G., and Fertig, E.J. (2023). Transcriptomic forecasting with neural ordinary differential equations. Patterns (N Y) 4, 100793. 10.1016/j.patter.2023.100793.

11. Macklin, P. (2017). When Seeing Isn’t Believing: How Math Can Guide Our Interpretation of Measurements and Experiments. Cell Syst 5, 92–94. 10.1016/j.cels.2017.08.005.

12. Materi, W., and Wishart, D.S. (2007). Computational systems biology in cancer: modeling methods and applications. Gene Regul Syst Bio 1, 91–110.

13. Metzcar, J., Wang, Y.F., Heiland, R., and Macklin, P. (2019). A Review of Cell-Based Computational Modeling in Cancer Biology. Jco Clin Cancer Info 3. 10.1200/Cci.18.00069.

14. Vodovotz, Y., and An, G. (2014). Translational Systems Biology (Academic Press). 10.1016/c2012-0-00091-9.

15. Vodovotz, Y., and An, G. (2019). Agent-based models of inflammation in translational systems biology: A decade later. Wiley Interdiscip Rev Syst Biol Med 11, e1460. 10.1002/wsbm.1460.

16. An, G., Mi, Q., Dutta-Moscato, J., and Vodovotz, Y. (2009). Agent-based models in translational systems biology. Wiley Interdiscip Rev Syst Biol Med 1, 159–171. 10.1002/wsbm.45.

17. An, G. (2008). Introduction of an agent-based multi-scale modular architecture for dynamic knowledge representation of acute inflammation. Theoretical Biology and Medical Modelling 5. 10.1186/1742-4682-5-11.

18. Scott, J.G., Fletcher, A.G., Anderson, A.R., and Maini, P.K. (2016). Spatial Metrics of Tumour Vascular Organisation Predict Radiation Efficacy in a Computational Model. PLoS Comput Biol 12, e1004712. 10.1371/journal.pcbi.1004712.

19. Bozic, I., Reiter, J.G., Allen, B., Antal, T., Chatterjee, K., Shah, P., Moon, Y.S., Yaqubie, A., Kelly, N., Le, D.T., et al. (2013). Evolutionary dynamics of cancer in response to targeted combination therapy. Elife 2, e00747. 10.7554/eLife.00747.

20. Cavenee, W.K., Dryja, T.P., Phillips, R.A., Benedict, W.F., Godbout, R., Gallie, B.L., Murphree, A.L., Strong, L.C., and White, R.L. (1983). Expression of recessive alleles by chromosomal mechanisms in retinoblastoma. Nature 305, 779–784. 10.1038/305779a0.

21. Diaz, L.A., Jr., Williams, R.T., Wu, J., Kinde, I., Hecht, J.R., Berlin, J., Allen, B., Bozic, I., Reiter, J.G., Nowak, M.A., et al. (2012). The molecular evolution of acquired resistance to targeted EGFR blockade in colorectal cancers. Nature 486, 537–540. 10.1038/nature11219.

22. Frank, S.A., Iwasa, Y., and Nowak, M.A. (2003). Patterns of cell division and the risk of cancer. Genetics 163, 1527–1532. 10.1093/genetics/163.4.1527.

23. Jones, S., Chen, W.D., Parmigiani, G., Diehl, F., Beerenwinkel, N., Antal, T., Traulsen, A., Nowak, M.A., Siegel, C., Velculescu, V.E., et al. (2008). Comparative lesion sequencing provides insights into tumor evolution. Proc Natl Acad Sci U S A 105, 4283–4288. 10.1073/pnas.0712345105.

24. Knudson, A.G., Jr. (1971). Mutation and cancer: statistical study of retinoblastoma. Proc Natl Acad Sci U S A 68, 820–823. 10.1073/pnas.68.4.820.

25. Komarova, N.L., and Wodarz, D. (2005). Drug resistance in cancer: principles of emergence and prevention. Proc Natl Acad Sci U S A 102, 9714–9719. 10.1073/pnas.0501870102.

26. Leder, K., Pitter, K., LaPlant, Q., Hambardzumyan, D., Ross, B.D., Chan, T.A., Holland, E.C., and Michor, F. (2014). Mathematical modeling of PDGF-driven glioblastoma reveals optimized radiation dosing schedules. Cell 156, 603–616. 10.1016/j.cell.2013.12.029.

27. Lenaerts, T., Pacheco, J.M., Traulsen, A., and Dingli, D. (2010). Tyrosine kinase inhibitor therapy can cure chronic myeloid leukemia without hitting leukemic stem cells. Haematologica 95, 900-907. 10.3324/haematol.2009.015271.

28. Sanga, S., Sinek, J.P., Frieboes, H.B., Ferrari, M., Fruehauf, J.P., and Cristini, V. (2006). Mathematical modeling of cancer progression and response to chemotherapy. Expert Rev Anticancer Ther 6, 1361–1376. 10.1586/14737140.6.10.1361.

29. Sherratt, J.A., and Nowak, M.A. (1992). Oncogenes, anti-oncogenes and the immune response to cancer: a mathematical model. Proc Biol Sci 248, 261–271. 10.1098/rspb.1992.0071.

30. Swanson, K.R., Alvord, E.C., Jr., and Murray, J.D. (2002). Virtual brain tumours (gliomas) enhance the reality of medical imaging and highlight inadequacies of current therapy. Br J Cancer 86, 14–18. 10.1038/sj.bjc.6600021.

31. Lu, Y., Ng, A.H.C., Chow, F.E., Everson, R.G., Helmink, B.A., Tetzlaff, M.T., Thakur, R., Wargo, J.A., Cloughesy, T.F., Prins, R.M., and Heath, J.R. (2021). Resolution of tissue signatures of therapy response in patients with recurrent GBM treated with neoadjuvant anti-PD1. Nat Commun 12, 4031. 10.1038/s41467-021-24293-4.

32. Pereira, E.J., Burns, J.S., Lee, C.Y., Marohl, T., Calderon, D., Wang, L., Atkins, K.A., Wang, C.C., and Janes, K.A. (2020). Sporadic activation of an oxidative stress-dependent NRF2-p53 signaling network in breast epithelial spheroids and premalignancies. Sci Signal 13. 10.1126/scisignal.aba4200.

33. Kim, M., Park, J., Bouhaddou, M., Kim, K., Rojc, A., Modak, M., Soucheray, M., McGregor, M.J., O’Leary, P., Wolf, D., et al. (2021). A protein interaction landscape of breast cancer. Science 374, eabf3066. 10.1126/science.abf3066.

34. Stahlberg, E.A., Abdel-Rahman, M., Aguilar, B., Asadpoure, A., Beckman, R.A., Borkon, L.L., Bryan, J.N., Cebulla, C.M., Chang, Y.H., Chatterjee, A., et al. (2022). Exploring approaches for predictive cancer patient digital twins: Opportunities for collaboration and innovation. Front Digit Health 4, 1007784. 10.3389/fdgth.2022.1007784.

35. Laubenbacher, R., Niarakis, A., Helikar, T., An, G., Shapiro, B., Malik-Sheriff, R.S., Sego, T.J., Knapp, A., Macklin, P., and Glazier, J.A. (2022). Building digital twins of the human immune system: toward a roadmap. NPJ Digit Med 5, 64. 10.1038/s41746-022-00610-z.

36. Hernandez-Boussard, T., Macklin, P., Greenspan, E.J., Gryshuk, A.L., Stahlberg, E., Syeda-Mahmood, T., and Shmulevich, I. (2021). Digital twins for predictive oncology will be a paradigm shift for precision cancer care. Nat Med 27, 2065–2066. 10.1038/s41591-021-01558-5.

37. Madhavan, S., Beckman, R.A., McCoy, M.D., Pishvaian, M.J., Brody, J.R., and Macklin, P. (2021). Envisioning the future of precision oncology trials. Nat Cancer 2, 9–11. 10.1038/s43018-020-00163-8.

38. An, G. (2004). In silico experiments of existing and hypothetical cytokine-directed clinical trials using agent-based modeling. Crit Care Med 32, 2050–2060. 10.1097/01.ccm.0000139707.13729.7d.

39. Saint-Pierre, P., and Savy, N. (2023). Agent-based modeling in medical research, virtual baseline generator and change in patients’ profile issue. Int J Biostat. 10.1515/ijb-2022-0112.

40. Cosgrove, J., Butler, J., Alden, K., Read, M., Kumar, V., Cucurull-Sanchez, L., Timmis, J., and Coles, M. (2015). Agent-Based Modeling in Systems Pharmacology. CPT Pharmacometrics Syst Pharmacol 4, 615–629. 10.1002/psp4.12018.

41. Norton, K.A., Gong, C., Jamalian, S., and Popel, A.S. (2019). Multiscale Agent-Based and Hybrid Modeling of the Tumor Immune Microenvironment. Processes (Basel) 7. 10.3390/pr7010037.

42. Rockne, R.C., Hawkins-Daarud, A., Swanson, K.R., Sluka, J.P., Glazier, J.A., Macklin, P., Hormuth, D.A., Jarrett, A.M., Lima, E., Tinsley Oden, J., et al. (2019). The 2019 mathematical oncology roadmap. Phys Biol 16, 041005. 10.1088/1478-3975/ab1a09.

43. Rocha, H.L., Aguilar, B., Getz, M., Shmulevich, I., and Macklin, P. (2023). A multiscale model of pulmonary micrometastasis and immune surveillance: towards cancer patient digital twins. bioRxiv [**preprint**] 2023.10.17.562733. 10.1101/2023.10.17.562733.

44. Ghaffarizadeh, A., Heiland, R., Friedman, S.H., Mumenthaler, S.M., and Macklin, P. (2018). PhysiCell: An open source physics-based cell simulator for 3-D multicellular systems. PLOS Computational Biology 14, e1005991. 10.1371/journal.pcbi.1005991.

45. Mirams, G.R., Arthurs, C.J., Bernabeu, M.O., Bordas, R., Cooper, J., Corrias, A., Davit, Y., Dunn, S.J., Fletcher, A.G., Harvey, D.G., et al. (2013). Chaste: an open source C++ library for computational physiology and biology. PLoS Comput Biol 9, e1002970. 10.1371/journal.pcbi.1002970.

46. Kang, S., Kahan, S., McDermott, J., Flann, N., and Shmulevich, I. (2014). Biocellion: accelerating computer simulation of multicellular biological system models. Bioinformatics 30, 3101–3108. 10.1093/bioinformatics/btu498.

47. Swat, M.H., Thomas, G.L., Belmonte, J.M., Shirinifard, A., Hmeljak, D., and Glazier, J.A. (2012). Multi-scale modeling of tissues using CompuCell3D. Methods Cell Biol 110, 325–366. 10.1016/B978-0-12-388403-9.00013-8.

48. Breitwieser, L., Hesam, A., de Montigny, J., Vavourakis, V., Iosif, A., Jennings, J., Kaiser, M., Manca, M., Di Meglio, A., Al-Ars, Z., et al. (2022). BioDynaMo: a modular platform for highperformance agent-based simulation. Bioinformatics 38, 453–460. 10.1093/bioinformatics/btab649.

49. Starruss, J., de Back, W., Brusch, L., and Deutsch, A. (2014). Morpheus: a user-friendly modeling environment for multiscale and multicellular systems biology. Bioinformatics 30, 1331–1332. 10.1093/bioinformatics/btt772.

50. Daub, J.T., and Merks, R.M. (2015). Cell-based computational modeling of vascular morphogenesis using Tissue Simulation Toolkit. Methods Mol Biol 1214, 67–127. 10.1007/978-1-4939-1462-3_6.

51. Sluka, J.P., Shirinifard, A., Swat, M., Cosmanescu, A., Heiland, R.W., and Glazier, J.A. (2014). The cell behavior ontology: describing the intrinsic biological behaviors of real and model cells seen as active agents. Bioinformatics 30, 2367–2374. 10.1093/bioinformatics/btu210.

52. Friedman, S.H., Anderson, A.R.A., Bortz, D.M., Fletcher, A.G., Frieboes, H.B., Ghaffarizadeh, A., Grimes, D.R., Hawkins-Daarud, A., Hoehme, S., Juarez, E.F., et al. (2016). MultiCellDS: a community-developed standard for curating microenvironment-dependent multicellular data. bioRxiv **[preprint**] 090456 10.1101/090456.

53. Ashburner, M., Ball, C.A., Blake, J.A., Botstein, D., Butler, H., Cherry, J.M., Davis, A.P., Dolinski, K., Dwight, S.S., Eppig, J.T., et al. (2000). Gene Ontology: tool for the unification of biology. Nature Genetics 25, 25–29. 10.1038/75556.

54. Aleksander, S.A., Balhoff, J., Carbon, S., Cherry, J.M., Drabkin, H.J., Ebert, D., Feuermann, M., Gaudet, P., Harris, N.L., Hill, D.P., et al. (2023). The Gene Ontology knowledgebase in 2023. Genetics 224 10.1093/genetics/iyad031.

55. Börner, K., Teichmann, S.A., Quardokus, E.M., Gee, J.C., Browne, K., Osumi-Sutherland, D., Herr, B.W., Bueckle, A., Paul, H., Haniffa, M., et al. (2021). Anatomical structures, cell types and biomarkers of the Human Reference Atlas. Nature Cell Biology 23, 1117–1128. 10.1038/s41556-021-00788-6.

56. Germann, P., Marin-Riera, M., and Sharpe, J. (2019). ya||a: GPU-Powered Spheroid Models for Mesenchyme and Epithelium. Cell Systems 8, 261–266.e263. 10.1016/j.cels.2019.02.007.

57. Bravo, R.R., Baratchart, E., West, J., Schenck, R.O., Miller, A.K., Gallaher, J., Gatenbee, C.D., Basanta, D., Robertson-Tessi, M., and Anderson, A.R.A. (2020). Hybrid Automata Library: A flexible platform for hybrid modeling with real-time visualization. PLoS Comput Biol 16, e1007635. 10.1371/journal.pcbi.1007635.

58. Hoehme, S., and Drasdo, D. (2010). A cell-based simulation software for multi-cellular systems. Bioinformatics 26, 2641–2642. 10.1093/bioinformatics/btq437.

59. Rocha, H.L., Godet, I., Kurtoglu, F., Metzcar, J., Konstantinopoulos, K., Bhoyar, S., Gilkes, D.M., and Macklin, P. (2021). A persistent invasive phenotype in post-hypoxic tumor cells is revealed by fate mapping and computational modeling. iScience 24, 102935. 10.1016/j.isci.2021.102935.

60. Wang, Y., Brodin, E., Nishii, K., Frieboes, H.B., Mumenthaler, S.M., Sparks, J.L., and Macklin, P. (2021). Impact of tumor-parenchyma biomechanics on liver metastatic progression: a multi-model approach. Sci Rep 11, 1710. 10.1038/s41598-020-78780-7.

61. Jagiella, N., Muller, B., Muller, M., Vignon-Clementel, I.E., and Drasdo, D. (2016). Inferring Growth Control Mechanisms in Growing Multi-cellular Spheroids of NSCLC Cells from Spatial-Temporal Image Data. PLoS Comput Biol 12, e1004412. 10.1371/journal.pcbi.1004412.

62. Hamis, S., Kohandel, M., Dubois, L.J., Yaromina, A., Lambin, P., and Powathil, G.G. (2020). Combining hypoxia-activated prodrugs and radiotherapy in silico: Impact of treatment scheduling and the intra-tumoural oxygen landscape. PLoS Comput Biol 16, e1008041. 10.1371/journal.pcbi.1008041.

63. Anderson, A.R.A. (2007). A Hybrid Multiscale Model of Solid Tumour Growth and Invasion: Evolution and the Microenvironment. In Single-Cell-Based Models in Biology and Medicine, pp. 3–28. 10.1007/978-3-7643-8123-3_1.

64. Kingsley, J.L., Costello, J.R., Raghunand, N., and Rejniak, K.A. (2021). Bridging cell-scale simulations and radiologic images to explain short-time intratumoral oxygen fluctuations. PLOS Computational Biology 17. 10.1371/journal.pcbi.1009206.

65. Szabó, A., and Merks, R.M.H. (2013). Cellular Potts Modeling of Tumor Growth, Tumor Invasion, and Tumor Evolution. Frontiers in Oncology 3. 10.3389/fonc.2013.00087.

66. Van Liedekerke, P., Neitsch, J., Johann, T., Alessandri, K., Nassoy, P., and Drasdo, D. (2019). Quantitative cell-based model predicts mechanical stress response of growing tumor spheroids over various growth conditions and cell lines. PLOS Computational Biology 15. 10.1371/journal.pcbi.1006273.

67. McKeown, S.R. (2014). Defining normoxia, physoxia and hypoxia in tumours-implications for treatment response. Br J Radiol 87, 20130676. 10.1259/bjr.20130676.

68. Kinny-Köster, B., Guinn, S., Tandurella, J.A., Mitchell, J.T., Sidiropoulos, D.N., Loth, M., Lyman, M.R., Pucsek, A.B., Seppälä, T.T., Cherry, C., et al. (2022). Inflammatory Signaling in Pancreatic Cancer Transfers Between a Single-cell RNA Sequencing Atlas and Co-Culture. bioRxiv [preprint], 2022.2007.2014.500096. 10.1101/2022.07.14.500096.

69. Bell, A.T.F., Mitchell, J.T., Kiemen, A.L., Fujikura, K., Fedor, H., Gambichler, B., Deshpande, A., Wu, P.-H., Sidiropoulos, D.N., Erbe, R., et al. (2022). PanIN and CAF Transitions in Pancreatic Carcinogenesis Revealed with Spatial Data Integration. bioRxiv [preprint] 2022.07.16.500312. 10.1101/2022.07.16.500312.

70. Kiemen, A.L., Braxton, A.M., Grahn, M.P., Han, K.S., Babu, J.M., Reichel, R., Jiang, A.C., Kim, B., Hsu, J., Amoa, F., et al. (2022). CODA: quantitative 3D reconstruction of large tissues at cellular resolution. Nat Methods 19, 1490–1499. 10.1038/s41592-022-01650-9.

71. Chan-Seng-Yue, M., Kim, J.C., Wilson, G.W., Ng, K., Figueroa, E.F., O’Kane, G.M., Connor, A.A., Denroche, R.E., Grant, R.C., McLeod, J., et al. (2020). Transcription phenotypes of pancreatic cancer are driven by genomic events during tumor evolution. Nat Genet 52, 231–240. 10.1038/s41588-019-0566-9.

72. El-Kenawi, A., Gatenbee, C., Robertson-Tessi, M., Bravo, R., Dhillon, J., Balagurunathan, Y., Berglund, A., Vishvakarma, N., Ibrahim-Hashim, A., Choi, J., et al. (2019). Acidity promotes tumour progression by altering macrophage phenotype in prostate cancer. Br J Cancer 121, 556–566. 10.1038/s41416-019-0542-2.

73. Henze, A.T., and Mazzone, M. (2016). The impact of hypoxia on tumor-associated macrophages. J Clin Invest 126, 3672–3679. 10.1172/JCI84427.

74. Leblond, M.M., Gerault, A.N., Corroyer-Dulmont, A., MacKenzie, E.T., Petit, E., Bernaudin, M., and Valable, S. (2016). Hypoxia induces macrophage polarization and re-education toward an M2 phenotype in U87 and U251 glioblastoma models. Oncoimmunology 5, e1056442. 10.1080/2162402X.2015.1056442.

75. Waldman, A.D., Fritz, J.M., and Lenardo, M.J. (2020). A guide to cancer immunotherapy: from T cell basic science to clinical practice. Nat Rev Immunol 20, 651–668. 10.1038/s41577-020-0306-5.

76. Schoenfeld, A.J., and Hellmann, M.D. (2020). Acquired Resistance to Immune Checkpoint Inhibitors. Cancer Cell 37, 443–455. 10.1016/j.ccell.2020.03.017.

77. Barretina, J., Caponigro, G., Stransky, N., Venkatesan, K., Margolin, A.A., Kim, S., Wilson, C.J., Lehar, J., Kryukov, G.V., Sonkin, D., et al. (2012). The Cancer Cell Line Encyclopedia enables predictive modelling of anticancer drug sensitivity. Nature 483, 603–607. 10.1038/nature11003.

78. Garnett, M.J., Edelman, E.J., Heidorn, S.J., Greenman, C.D., Dastur, A., Lau, K.W., Greninger, P., Thompson, I.R., Luo, X., Soares, J., et al. (2012). Systematic identification of genomic markers of drug sensitivity in cancer cells. Nature 483, 570–575. 10.1038/nature11005.

79. Johnson, B.E., Creason, A.L., Stommel, J.M., Keck, J.M., Parmar, S., Betts, C.B., Blucher, A., Boniface, C., Bucher, E., Burlingame, E., et al. (2022). An omic and multidimensional spatial atlas from serial biopsies of an evolving metastatic breast cancer. Cell Rep Med 3, 100525. 10.1016/j.xcrm.2022.100525.

80. Tatarova, Z., Blumberg, D.C., Korkola, J.E., Heiser, L.M., Muschler, J.L., Schedin, P.J., Ahn, S.W., Mills, G.B., Coussens, L.M., Jonas, O., and Gray, J.W. (2022). A multiplex implantable microdevice assay identifies synergistic combinations of cancer immunotherapies and conventional drugs. Nat Biotechnol 40, 1823–1833. 10.1038/s41587-022-01379-y.

81. Muller, E., Christopoulos, P.F., Halder, S., Lunde, A., Beraki, K., Speth, M., Oynebraten, I., and Corthay, A. (2017). Toll-Like Receptor Ligands and Interferon-gamma Synergize for Induction of Antitumor M1 Macrophages. Front Immunol 8, 1383. 10.3389/fimmu.2017.01383.

82. Coller, S.P., and Paulnock, D.M. (2001). Signaling pathways initiated in macrophages after engagement of type A scavenger receptors. J Leukoc Biol 70, 142–148.

83. Acosta-Iborra, B., Elorza, A., Olazabal, I.M., Martin-Cofreces, N.B., Martin-Puig, S., Miro, M., Calzada, M.J., Aragones, J., Sanchez-Madrid, F., and Landazuri, M.O. (2009). Macrophage oxygen sensing modulates antigen presentation and phagocytic functions involving IFN-gamma production through the HIF-1 alpha transcription factor. J Immunol 182, 3155–3164. 10.4049/jimmunol.0801710.

84. Ruffell, B., Chang-Strachan, D., Chan, V., Rosenbusch, A., Ho, C.M., Pryer, N., Daniel, D., Hwang, E.S., Rugo, H.S., and Coussens, L.M. (2014). Macrophage IL-10 blocks CD8+ T cell-dependent responses to chemotherapy by suppressing IL-12 expression in intratumoral dendritic cells. Cancer Cell 26, 623–637. 10.1016/j.ccell.2014.09.006.

85. Mantovani, A., Allavena, P., Marchesi, F., and Garlanda, C. (2022). Macrophages as tools and targets in cancer therapy. Nat Rev Drug Discov 21, 799–820. 10.1038/s41573-022-00520-5.

86. van der Leun, A.M., Thommen, D.S., and Schumacher, T.N. (2020). CD8(+) T cell states in human cancer: insights from single-cell analysis. Nat Rev Cancer 20, 218–232. 10.1038/s41568-019-0235-4.

87. Heumann, T., Judkins, C., Li, K., Lim, S.J., Hoare, J., Parkinson, R., Cao, H., Zhang, T., Gai, J., Celiker, B., et al. (2023). A platform trial of neoadjuvant and adjuvant antitumor vaccination alone or in combination with PD-1 antagonist and CD137 agonist antibodies in patients with resectable pancreatic adenocarcinoma. Nat Commun 14, 3650. 10.1038/s41467-023-39196-9.

88. Li, K., Tandurella, J.A., Gai, J., Zhu, Q., Lim, S.J., Thomas, D.L, 2nd., Xia, T., Mo, G., Mitchell, J.T., Montagne, J., et al. (2022). Multi-omic analyses of changes in the tumor microenvironment of pancreatic adenocarcinoma following neoadjuvant treatment with anti-PD-1 therapy. Cancer Cell. 10.1016/j.ccell.2022.10.001.

89. Kinny-Köster, B., Guinn, S., Tandurella, J.A., Mitchell, J.T., Sidiropoulos, D.N., Loth, M., Lyman, M.R., Pucsek, A.B., Seppälä, T.T., Cherry, C., et al. (2022). Inflammatory Signaling in Pancreatic Cancer Transfers Between a Single-cell RNA Sequencing Atlas and Co-Culture. bioRxiv **[preprint**] 2022.07.14.500096. 10.1101/2022.07.14.500096.

90. Jaffee, E.M., Hruban, R.H., Biedrzycki, B., Laheru, D., Schepers, K., Sauter, P.R., Goemann, M., Coleman, J., Grochow, L., Donehower, R.C., et al. (2001). Novel allogeneic granulocyte-macrophage colony-stimulating factor-secreting tumor vaccine for pancreatic cancer: a phase I trial of safety and immune activation. J Clin Oncol 19, 145–156. 10.1200/JCO.2001.19.1.145.

91. Lutz, E.R., Wu, A.A., Bigelow, E., Sharma, R., Mo, G., Soares, K., Solt, S., Dorman, A., Wamwea, A., Yager, A., et al. (2014). Immunotherapy converts nonimmunogenic pancreatic tumors into immunogenic foci of immune regulation. Cancer Immunol Res 2, 616–631. 10.1158/2326-6066.CIR-14-0027.

92. Ozik, J., Collier, N., Heiland, R., An, G., and Macklin, P. (2019). Learning-accelerated discovery of immune-tumour interactions. Mol Syst Des Eng 4, 747–760. 10.1039/c9me00036d.

93. Ozik, J., Collier, N., Wozniak, J.M., Macal, C., Cockrell, C., Friedman, S.H., Ghaffarizadeh, A., Heiland, R., An, G., and Macklin, P. (2018). High-throughput cancer hypothesis testing with an integrated PhysiCell-EMEWS workflow. BMC Bioinformatics 19, 483. 10.1186/s12859-018-2510x.

94. Heiland, R., Bergman, D., Lyons, B., Cass, J., Rocha, H.L., Ruscone, M., Noël, V., and Macklin, P. (2023). PhysiCell Studio: a graphical tool to make agent-based modeling more accessible. bioRxiv [preprint] 2023.10.24.563727. 10.1101/2023.10.24.563727.

95. Heiland, R., Mishler, D., Zhang, T., Bower, E., and Macklin, P. (2019). xml2jupyter: Mapping parameters between XML and Jupyter widgets. J Open Source Softw 4. 10.21105/joss.01408.

96. Saxena, G., Ponce-de-Leon, M., Montagud, A., Vicente Dorca, D., and Valencia, A. (2021). BioFVM-X: An MPI+OpenMP 3-D Simulator for Biological Systems. In Computational Methods in Systems Biology, pp. 266–279. 10.1007/978-3-030-85633-5_18.

97. Ghaffarizadeh, A., Friedman, S.H., and Macklin, P. (2016). BioFVM: an efficient, parallelized diffusive transport solver for 3-D biological simulations. Bioinformatics 32, 1256–1258. 10.1093/bioinformatics/btv730.

98. Sundus, A., Kurtoglu, F., Konstantinopoulos, K., Chen, M., Willis, D., Heiland, R., and Macklin, P. (2022). PhysiCell training apps: Cloud hosted open-source apps to learn cell-based simulation software. bioRxiv [preprint] 10.1101/2022.06.24.497566.

99. Jenner, A.L., Smalley, M., Goldman, D., Goins, W.F., Cobbs, C.S., Puchalski, R.B., Chiocca, E.A., Lawler, S., Macklin, P., Goldman, A., and Craig, M. (2022). Agent-based computational modeling of glioblastoma predicts that stromal density is central to oncolytic virus efficacy. iScience 25, 104395. 10.1016/j.isci.2022.104395.

100. Islam, M.A., Getz, M., Macklin, P., and Versypt, A.N.F. (2022). An agent-based modeling approach for lung fibrosis in response to COVID-19. bioRxiv **[preprint**]. 10.1101/2022.10.03.510677.

101. Getz, M., Wang, Y., An, G., Asthana, M., Becker, A., Cockrell, C., Collier, N., Craig, M., Davis, C.L., Faeder, J.R., et al. (2021). Iterative community-driven development of a SARS-CoV-2 tissue simulator. bioRxiv **[preprint]**. 10.1101/2020.04.02.019075.

102. Risner, K.H., Tieu, K.V., Wang, Y., Bakovic, A., Alem, F., Bhalla, N., Nathan, S., Conway, D.E., Macklin, P., and Narayanan, A. (2020). Maraviroc inhibits SARS-CoV-2 multiplication and sprotein mediated cell fusion in cell culture. bioRxiv **[preprint]**. 10.1101/2020.08.12.246389.

103. Letort, G., Montagud, A., Stoll, G., Heiland, R., Barillot, E., Macklin, P., Zinovyev, A., and Calzone, L. (2019). PhysiBoSS: a multi-scale agent-based modelling framework integrating physical dimension and cell signalling. Bioinformatics 35, 1188–1196. 10.1093/bioinformatics/bty766.

104. Hoehme, S., Friebel, A., Hammad, S., Drasdo, D., and Hengstler, J.G. (2017). Creation of ThreeDimensional Liver Tissue Models from Experimental Images for Systems Medicine. Methods Mol Biol 1506, 319–362. 10.1007/978-1-4939-6506-9_22.

105. Finley, S.D., and Popel, A.S. (2013). Effect of tumor microenvironment on tumor VEGF during anti-VEGF treatment: systems biology predictions. J Natl Cancer Inst 105, 802–811. 10.1093/jnci/djt093.

106. Swan, A., Hillen, T., Bowman, J.C., and Murtha, A.D. (2018). A Patient-Specific Anisotropic Diffusion Model for Brain Tumour Spread. Bull Math Biol 80, 1259–1291. 10.1007/s11538-017-0271-8.

107. Chaplain, M.A., Graziano, L., and Preziosi, L. (2006). Mathematical modelling of the loss of tissue compression responsiveness and its role in solid tumour development. Math Med Biol 23, 197–229. 10.1093/imammb/dql009.

108. Alarcon, T., Byrne, H.M., and Maini, P.K. (2003). A cellular automaton model for tumour growth in inhomogeneous environment. J Theor Biol 225, 257–274. 10.1016/s0022-5193(03)00244-3.

109. Scott, J.G., Basanta, D., Anderson, A.R., and Gerlee, P. (2013). A mathematical model of tumour self-seeding reveals secondary metastatic deposits as drivers of primary tumour growth. J R Soc Interface 10, 20130011. 10.1098/rsif.2013.0011.

110. Kaznatcheev, A., Vander Velde, R., Scott, J.G., and Basanta, D. (2017). Cancer treatment scheduling and dynamic heterogeneity in social dilemmas of tumour acidity and vasculature. Br J Cancer 116, 785–792. 10.1038/bjc.2017.5.

111. Poleszczuk, J., Hahnfeldt, P., and Enderling, H. (2014). Biphasic modulation of cancer stem cell-driven solid tumour dynamics in response to reactivated replicative senescence. Cell Prolif 47, 267–276. 10.1111/cpr.12101.

112. Powathil, G.G., Adamson, D.J., and Chaplain, M.A. (2013). Towards predicting the response of a solid tumour to chemotherapy and radiotherapy treatments: clinical insights from a computational model. PLoS Comput Biol 9, e1003120. 10.1371/journal.pcbi.1003120.

113. Hamis, S., Nithiarasu, P., and Powathil, G.G. (2018). What does not kill a tumour may make it stronger: In silico insights into chemotherapeutic drug resistance. J Theor Biol 454, 253–267. 10.1016/j.jtbi.2018.06.014.

114. Fortuna, I., Perrone, G.C., Krug, M.S., Susin, E., Belmonte, J.M., Thomas, G.L., Glazier, J.A., and de Almeida, R.M.C. (2020). CompuCell3D Simulations Reproduce Mesenchymal Cell Migration on Flat Substrates. Biophys J 118, 2801–2815. 10.1016/j.bpj.2020.04.024.

115. Dunn, S.J., Appleton, P.L., Nelson, S.A., Nathke, I.S., Gavaghan, D.J., and Osborne, J.M. (2012). A two-dimensional model of the colonic crypt accounting for the role of the basement membrane and pericryptal fibroblast sheath. PLoS Comput Biol 8, e1002515. 10.1371/journal.pcbi.1002515.

116. Glen, C.M., Kemp, M.L., and Voit, E.O. (2019). Agent-based modeling of morphogenetic systems: Advantages and challenges. PLoS Comput Biol 15, e1006577. 10.1371/journal.pcbi.1006577.

117. Schubert, M., Dokmegang, J., Yap, M.H., Han, L., Cavaliere, M., and Doursat, R. (2021). Computational modelling unveils how epiblast remodelling and positioning rely on trophectoderm morphogenesis during mouse implantation. Plos One 16. 10.1371/journal.pone.0254763.

118. Camacho-Gómez, D., García-Aznar, J.M., and Gómez-Benito, M.J. (2022). A 3D multi-agent-based model for lumen morphogenesis: the role of the biophysical properties of the extracellular matrix. Engineering with Computers 38, 4135–4149. 10.1007/s00366-022-01654-1.

119. Cess, C.G., and Finley, S.D. (2020). Multi-scale modeling of macrophage—T cell interactions within the tumor microenvironment. PLOS Computational Biology 16. 10.1371/journal.pcbi.1008519.

120. Ruiz-Martinez, A., Gong, C., Wang, H., Sove, R.J., Mi, H., Kimko, H., and Popel, A.S. (2022). Simulations of tumor growth and response to immunotherapy by coupling a spatial agent-based model with a whole-patient quantitative systems pharmacology model. PLoS Comput Biol 18, e1010254. 10.1371/journal.pcbi.1010254.

121. Ni, C., and Lu, T. (2022). Individual-Based Modeling of Spatial Dynamics of Chemotactic Microbial Populations. ACS Synth Biol 11, 3714–3723. 10.1021/acssynbio.2c00322.

122. Hellweger, F.L., and Bucci, V. (2009). A bunch of tiny individuals—Individual-based modeling for microbes. Ecological Modelling 220, 8–22. 10.1016/j.ecolmodel.2008.09.004.

123. Osinska, I., Popko, K., and Demkow, U. (2014). Perforin: an important player in immune response. Cent Eur J Immunol 39, 109–115. 10.5114/ceji.2014.42135.

124. Farhood, B., Najafi, M., and Mortezaee, K. (2019). CD8(+) cytotoxic T lymphocytes in cancer immunotherapy: A review. J Cell Physiol 234, 8509–8521. 10.1002/jcp.27782.

125. Raskov, H., Orhan, A., Christensen, J.P., and Gogenur, I. (2021). Cytotoxic CD8(+) T cells in cancer and cancer immunotherapy. Br J Cancer 124, 359–367. 10.1038/s41416-020-01048-4.

126. Sharma, G., Colantuoni, C., Goff, L.A., Fertig, E.J., and Stein-O’Brien, G. (2020). projectR: an R/Bioconductor package for transfer learning via PCA, NMF, correlation and clustering. Bioinformatics 36, 3592–3593. 10.1093/bioinformatics/btaa183.

127. Johnson, J., Tsang, A., Mitchell, J.T., Davis-Marcisak, E., Sherman, T., Liefeld, T., Loth, M., Goff, L.A., Zimmerman, J., Kinny-Köster, B., et al. (2022). Inferring cellular and molecular processes in single-cell data with non-negative matrix factorization using Python, R, and GenePattern Notebook implementations of CoGAPS. bioRxiv [preprint] 2022.07.09.499398. 10.1101/2022.07.09.499398.

128. Barrett, T., Wilhite, S.E., Ledoux, P., Evangelista, C., Kim, I.F., Tomashevsky, M., Marshall, K.A., Phillippy, K.H., Sherman, P.M., Holko, M., et al. (2013). NCBI GEO: archive for functional genomics data sets--update. Nucleic Acids Res 41, D991–995. 10.1093/nar/gks1193.

129. Edgar, R., Domrachev, M., and Lash, A.E. (2002). Gene Expression Omnibus: NCBI gene expression and hybridization array data repository. Nucleic Acids Res 30, 207–210. 10.1093/nar/30.1.207.

130. Peng, J., Sun, B.F., Chen, C.Y., Zhou, J.Y., Chen, Y.S., Chen, H., Liu, L., Huang, D., Jiang, J., Cui, G.S., et al. (2019). Single-cell RNA-seq highlights intra-tumoral heterogeneity and malignant progression in pancreatic ductal adenocarcinoma. Cell Res 29, 725–738. 10.1038/s41422-0190195-y.

131. Steele, N.G., Carpenter, E.S., Kemp, S.B., Sirihorachai, V.R., The, S., Delrosario, L., Lazarus, J., Amir, E.D., Gunchick, V., Espinoza, C., et al. (2020). Multimodal Mapping of the Tumor and Peripheral Blood Immune Landscape in Human Pancreatic Cancer. Nat Cancer 1, 1097–1112. 10.1038/s43018-020-00121-4.

132. Cherry, C., Maestas, D.R., Han, J., Andorko, J.I., Cahan, P., Fertig, E.J., Garmire, L.X., and Elisseeff, J.H. (2021). Computational reconstruction of the signalling networks surrounding implanted biomaterials from single-cell transcriptomics. Nat Biomed Eng 5, 1228–1238. 10.1038/s41551-021-00770-5.

133. Sherman, T.D., Gao, T., and Fertig, E.J. (2020). CoGAPS 3: Bayesian non-negative matrix factorization for single-cell analysis with asynchronous updates and sparse data structures. BMC Bioinformatics 21, 453. 10.1186/s12859-020-03796-9.

134. Fertig, E.J., Ding, J., Favorov, A.V., Parmigiani, G., and Ochs, M.F. (2010). CoGAPS: an R/C++ package to identify patterns and biological process activity in transcriptomic data. Bioinformatics 26, 2792–2793. 10.1093/bioinformatics/btq503.

135. Hao, Y., Hao, S., Andersen-Nissen, E., Mauck, W.M3rd., Zheng, S., Butler, A., Lee, M.J., Wilk, A.J., Darby, C., Zager, M., et al. (2021). Integrated analysis of multimodal single-cell data. Cell 184, 3573–3587 e3529. 10.1016/j.cell.2021.04.048.

136. PhysiCell (2023). PhysiCell Version 1.12.0. https://github.com/MathCancer/PhysiCell/releases/tag/1.12.0.

137. Wortel, I.M., and Textor, J. (2021). Artistoo, a library to build, share, and explore simulations of cells and tissues in the web browser. Elife 10. 10.7554/eLife.61288.

138. Heiland, R., and Macklin, P. (2023). PhysiCell Studio Cloud (Version 1.0). https://nanohub.org/tools/pcstudio.

139. Madhavan, K., Zentner, L., Farnsworth, V., Shivarajapura, S., Zentner, M., Denny, N., and Klimeck, G. (2013). nanoHUB.org: cloud-based services for nanoscale modeling, simulation, and education. Nanotechnology Reviews 2, 107–117. 10.1515/ntrev-2012-0043.

