## Supplementary Information for "Digitize your Biology! Modeling multicellular systems through interpretable cell behavior"

### REFERNCE CELL BEHAVIORS AND SIGNALS

#### Cell behaviors and reference models

To build this grammar, we require clear abstractions of key cell behaviors that frequently occur in multicellular observations and corresponding reference models. In this context, a cell *behavior* is a cell-scale process, such as cycling, death, or phagocytosis. Generally, each behavior can be represented by a small number of continuous phenotypic parameters, describing the rate, magnitude, or frequency of the behavior. In earlier work, Sluka et al. developed the Cell Behavior Ontology (CBO)[1] as a controlled vocabulary of individual cell behaviors. More recently, we worked with a multidisciplinary coalition to extend and structure behaviors from the CBO and other sources into MultiCellDS[2] (multicellular data standard). In particular, this work defined a *behavioral cell phenotype* that collects of biophysical characterizations of a cell's behavior, organized hierarchically by function: cycling, death, volume, mechanics, secretion (including uptake), and motility. Since releasing MultiCellDS as a preprint, we have tested this approach to cell behavior through a variety of agent-based simulation and modeling projects[3-13]. Based upon recent immunologic modeling work[4,5,8,9], we extended phenotype to include cell-cell interactions (phagocytosis, effector attack, and fusion), as well as transformations between cell types (e.g., differentiation, transdifferentiation, and other persistent state changes).

Here, we fully describe the grammar's supported cell behaviors, as well as their *reference implementation* in PhysiCell and corresponding biophysical parameters.

***Cycling***

As introduced in prior work[2,13,14], a cell cycle model is a sequence of phases (with indices $i = 0,1, \dots, n$) and transition rates (or exit rates) $\{r_i\}$ connecting phase $i$ to the next phase $i + 1$. By convention, exit from the first phase (with index 0) is called *cycle entry*, and cells divide upon exiting last phase (with index $n$) to return to the 0th phase. In the reference implementation, cell transitions between phases are probabilistic, based upon the transition rate; if a cell is in phase $i$ at time $t$, then the probability of advancing to phase $i + 1$ between the current time $t$ and a future time $t + \Delta t$ is given by

$$P(\text{exit phase } i) = r_i \Delta t. \tag{1}$$

Note that the mean time spent in the $i^{\text{th}}$ phase is $\frac{1}{r_i}$.

For example, a five-phase model ("flow cytometry separated") consists of phases $G_0$, $G_1$, S, $G_2$, and M, cycle entry is a transition from $G_0$ (Phase 0) to $G_1$ (Phase 1), and division occurs at the end of M (Phase 4). In the reference implementation[13], at division daughter cells are assigned half the volume of the parent cell and positioned randomly such that (1) they both fit within the parent cell's volume, and (2) the daughter cells have the same center of volume as their parent cell. In the current grammar, the dictionary of cell cycling behaviors and their corresponding parameters are in **Table 1.** Future versions of the grammar may include finer grained control on placement of daughter cells after division.

| **dictionary symbol** | **synonyms** | **controllable phenotype parameter** |
|---|---|---|
| exit from cycle phase 0 | cycle entry | $r_0$ |
| exit from cycle phase 1 | | $r_1$ |
| exit from cycle phase 2 | | $r_2$ |
| exit from cycle phase 3 | | $r_3$ |
| exit from cycle phase 4 | | $r_4$ |
| exit from cycle phase 5 | | $r_5$ |

**TABLE 1.** Cell cycle terms in the behavior dictionary, along with corresponding controllable parameters in the PhysiCell reference implementation.

***Death***

In the initial grammar, we support apoptosis (as a primary form of non-immunogenic cell death) and necrosis (as a key form of immunogenic cell death). We note that at a cellular and multicellular perspective, death is not merely a discrete event, but rather entry into a cascade of processes[15-20]; as noted in prior modeling and analysis[13,15,21], every model of death has a nonzero, finite duration on the order of hours (for apoptosis) to days or weeks (for necrosis). In the reference implementation[13], live cells have an apoptotic death rate $d_A$, such that the probability that a cell becomes apoptotic between the current time $t$ and a future time $t + \Delta t$ is given by

$$P(\text{cell becomes apoptotic}) = d_A \Delta t. \tag{2}$$

Once the cell becomes apoptotic, it shrinks until being removed from the simulation.

Similarly, live cells have a necrotic death rate $d_N$, such that the probability that a cell becomes necrotic between the current time $t$ and a future time $t + \Delta t$ is given by

$$P(\text{cell becomes necrotic}) = d_N \Delta t \tag{3}$$

In the reference necrosis model, cells initially swell until reaching a bursting volume (by default, 200% of the cell's volume at the start of necrosis), and then gradually shrink over the course of days.

Full details on the reference parameter values for apoptosis and necrosis (including rates of cell volume change and durations) are found in prior work[13,21]. In the current grammar, the dictionary of behaviors and their corresponding parameters are in **Table 2.**

| dictionary symbol | synonyms | controllable phenotype parameter |
|---|---|---|
| apoptosis | | $d_A$ |
| necrosis | | $d_N$ |

**TABLE 2.** Death terms in the behavior dictionary, along with corresponding controllable parameters in the PhysiCell reference implementation.

##### *Secretion, uptake, and generalized chemical export*

Cells secrete and uptake (consume) chemical factors that diffuse in the extracellular microenvironment, with a fixed mathematical form first introduced in BioFVM[22] and later incorporated in PhysiCell[13]. For any diffusible substrate $\rho$, BioFVM solves the reaction-diffusion equation in Equation (4):

$$\frac{\partial \rho}{\partial t} = D\nabla^2\rho - \lambda\rho + \sum_{\text{cells } i}\left(\delta(\boldsymbol{x}-\boldsymbol{x}_i)V_i\left[\overbrace{S_i(\rho_i^*-\rho)}^{\text{secretion}} - \overbrace{U_i\rho}^{\text{uptake}}\right] + \delta(\boldsymbol{x}-\boldsymbol{x}_i)\overbrace{E_i}^{\text{export}}\right) \tag{4}$$

Here, $\boldsymbol{x}_i$ is the $i^{\text{th}}$ cell's position, $\delta(\boldsymbol{x})$ is the Dirac delta function that mathematically focuses the cell-based source/sink at its center, $S_i$ is its secretion rate of the substrate (with dimensions 1/time), $\rho_i^*$ is the "target value" of its secretion (i.e., secretion slows as $\rho$ approaches this value), and $U_i$ is its uptake or consumption rate (also with dimensions 1/time). When these fixed secretion and uptake forms are insufficient to match experiments or advanced modeling forms, we provide a generic *net export* term $E_i$ (with dimensions substrate/cell/time). Because the microenvironment can have multiple diffusing substrates, the grammar can automatically expand to reference secretion, uptake, and export parameters for each one, therefore defining a family of symbols as summarized in **Table 3**.

| dictionary symbol | synonyms | controllable phenotype parameter |
|---|---|---|
| *X* secretion | | $S_i$ |
| *X* secretion target | *X* secretion saturation density | $\rho_i^*$ |
| *X* uptake | | $U_i$ |
| *X* export | | $E_i$ |

**TABLE 3.** Secretion, uptake, and export terms in the behavior dictionary, along with corresponding controllable parameters in the PhysiCell reference implementation. *X* denotes the name of any diffusible substrate (e.g., oxygen).

##### *(Biased) migration and chemotaxis*

As previously introduced in PhysiCell[13], cell migration (or motility) is represented as a biased random walk: the cell's migration direction $\boldsymbol{d}_{\text{mot}}$ is a combination of a random (unit vector) component $\boldsymbol{\xi}$ and a non-random directed component (a migration bias direction) $\boldsymbol{d}_{\text{bias}}$, which is normalized to be a unit vector as in Equation (5):

$$\boldsymbol{d}_{\text{mot}} = \frac{b\,\boldsymbol{d}_{\text{bias}} + (1-b)\boldsymbol{\xi}}{\|b\,\boldsymbol{d}_{\text{bias}} + (1-b)\boldsymbol{\xi}\|} \tag{5}$$

Here, $0 \leq b \leq 1$ is the cell's *migration bias*: if $b = 1$, then migration occurs completely along the bias direction $\boldsymbol{d}_{\text{bias}}$, while $b = 0$ corresponds to purely Brownian motion. We then obtain the *migration velocity* $\boldsymbol{v}_{\text{mot}}$ by multiplying the direction by the *migration speed* ($s$):

$$\boldsymbol{v}_{\text{mot}} = s\,\boldsymbol{d}_{\text{mot}}. \tag{6}$$

In our formulation, the cell's migration speed (and hence migration velocity) can change dynamically based upon any further modeling rules, even if the migration direction does not. Cell migration has a *persistence time* $T_{\text{persist}}$: between the current time $t$ and a future time $t + \Delta t$, the probably of choosing a new migration direction (by re-evaluating Equation (6)) is given by

$$P(\text{change migration direction}) = \frac{\Delta t}{T_{\text{persist}}}. \tag{7}$$

Note that this gives a mean time of $T_{\text{persist}}$ between direction changes.

Chemotaxis is modeled by setting the migration bias direction along the gradient of one or more chemical substrates in the microenvironment. If available chemical substrates are $c_0, c_1, \ldots, c_m$, then we set

$$\boldsymbol{d}_{\text{bias}} = \omega_0 \nabla c_0 + \cdots + \omega_m \nabla c_m, \tag{8}$$

where $\omega_i$ is a weighting (*chemotactic response*) for each chemical gradient: $\omega_i < 0$ for migration against the gradient (along $-\nabla c_i$), $\omega_i > 0$ for migration along the gradient, and $\omega_i = 0$ if the gradient makes no contribution. Because the microenvironment can have multiple diffusing substrates, the grammar can automatically expand to a family of chemotactic response terms. See **Table 4.**

| dictionary symbol | synonyms | controllable phenotype parameter |
|---|---|---|
| migration speed | | $s$ |
| migration bias | | $b$ |
| migration persistence time | | $T_{\text{persist}}$ |
| chemotactic response to *X* | chemotactic sensitivity to *X* | $\omega_i$ |

**TABLE 4.** Migration terms in the behavior dictionary, along with corresponding controllable parameters in the PhysiCell reference implementation. *X* denotes the name of any diffusible substrate (e.g., oxygen).

##### *Cell-cell adhesion*

Two models of cell-cell adhesion are supported: a looser cell-cell adhesion using potential functions (as in prior mathematical models[23-28]), and more persistent elastic springs[6,29,30]. In the reference PhysiCell implementation for potential-based adhesions, if cells $i$ and $j$ are adhered and within interaction distance, then the contribution to the velocity of cell $i$ is given by

$$\sqrt{A_{ij}\,\alpha_i \cdot A_{ji}\alpha_j}\left(1 - \frac{\|\boldsymbol{x}_j - \boldsymbol{x}_i\|}{R_{A,i}R_i + R_{A,j}R_j}\right)^{n+1} \frac{(\boldsymbol{x}_j - \boldsymbol{x}_i)}{\|\boldsymbol{x}_j - \boldsymbol{x}_i\|}, \tag{9}$$

where $\alpha_i$ is cell $i$'s adhesive strength, $A_{ij}$ is the (relative) adhesive affinity of cell $i$ to cell $j$ (which for example could be modeled based upon adhesive receptor expressions of cells $i$ and $j$), $\boldsymbol{x}_i$ is the position of cell $i$, and $R_{A,i}$ is cell $i$'s relative maximum adhesion distance (the largest distance it can extend for adhesion, as a multiple of its radius), and $R_i$ is the cell's radius[13]. Notice that this adhesion only depends upon the relative distance between cells, and not their prior history. If the cells are identical, the coefficient reduces to $A_{ii}\alpha_i$. See **Table 5**.

If more sophisticated cell-cell adhesion is required (e.g., to model that cell adhesions may form more readily than they break), we also support an elastic cell-cell adhesion model. Cell $i$ forms an elastic adhesion to cell $j$ between the current time $t$ and a future time $t + \Delta t$ at an attachment rate $r_{A,ij}$ (and thus probability $r_{A,ij}\Delta t$), provided that:

- The distance $\|\boldsymbol{x}_j - \boldsymbol{x}_i\|$ between the cells does not exceed the maximum adhesion interaction distance, and
- Neither cell $i$ nor cell $j$ has exceeded their individual maximum number of cell adhesions ($n_{M,i}$ and $n_{M,j}$).

We calculate this cell-cell attachment rate $r_{A,ij}$ as

$$r_{A,ij} = r_{A,i}\, A_{ij}, \tag{10}$$

where $r_{A,i}$ is cell $i$'s rate of forming elastic attachments and $A_{ij}$ is the adhesive affinity of cell $i$ to cell $j$. (Notice that cell $i$ can form attachments to cell $j$, and cell $j$ can form attachments to cell $i$ independently with independent rates.) Analogously, cell $i$ can detach from cell $j$ at rate $u_{D,i}$, so that during any time interval $t$ to $t + \Delta t$, the cell $i$ can detach from cell $j$ with probability $u_{D,i}\Delta t$. As previously modeled[ref], when the cells are elastically adhered, the adhesion contributes to cell $i$ velocity via:

$$\sqrt{A_{ij}\,\epsilon_i \cdot A_{ji}\epsilon_j} \cdot (\boldsymbol{x}_j - \boldsymbol{x}_i), \tag{11}$$

where $\epsilon_i$ is cell $i$'s elastic constant. Notice that if cells $i$ and $j$ are identical, then the coefficient simplifies to $A_{ij}\epsilon_i$. The full parameter list for basic and elastic cell-cell adhesion (and cell repulsion) is found in **Table 5.**

| dictionary symbol | synonyms | controllable phenotype parameter |
|---|---|---|
| cell-cell adhesion | | $\alpha_i$ |
| cell-cell adhesion elastic constant | | $\epsilon_i$ |
| adhesive affinity to X | adhesive affinity to cell type X | $A_{ij}$ |
| relative maximum adhesion distance | | $\frac{R_{A,i}}{R_i}$ |
| cell attachment rate | | $r_{A,i}$ |
| cell detachment rate | | $u_{D,i}$ |
| maximum number of cell attachments | | $n_{M,i}$ |
| cell-cell repulsion | | $\beta_i$ |
| movable | is_movable, is movable | $m$ |

**Table 5.** Cell-cell adhesion and repulsion terms in the behavior dictionary, along with corresponding controllable param- [illegible]

#### Resistance to deformation and movement

PhysiCell[13], as many agent-based simulation frameworks[23-27], uses cell "repulsion" to model resistance to deformation, compression, and cell overlap. In the reference PhysiCell implementation for potential-based repulsion, if cells $i$ and $j$ are in contact, then the contribution to the velocity of cell $i$ is given by

$$-\sqrt{\beta_i \cdot \beta_j}\left(1 - \frac{\|\boldsymbol{x}_j - \boldsymbol{x}_i\|}{R_i + R_j}\right)^{n+1} \frac{(\boldsymbol{x}_j - \boldsymbol{x}_i)}{\|\boldsymbol{x}_j - \boldsymbol{x}_i\|}, \tag{12}$$

where $R_i$ is cell $i$'s radius and $\beta_i$ is its cell-cell repulsion coefficient as defined in prior work [ref]. In cases where a cell should be regarded as a rigid object or obstacle (e.g., if rigidly adhered to an underlying matrix), it can be flagged as *unmovable* using a "movable" parameter $m$. If $m$ is false ($m = 0$), then the cell can impart adhesive and repulsive forces on other cells, but they cannot impose reciprocal forces that contribute to the cell's movement. The full parameter list for cell-cell adhesion and repulsion is found in **Table 5**.

##### *Transformation (changing type)*

Many biological systems require cell type transformations from one type to another, particularly via differentiation, transdifferentiation, and mutation. Mathematically, these all can be represented as a transformation (type change) rate $r_{T,ij}$from type $i$ to type $j$, with the probability of a type change between the current time $t$ and a future time $t + \Delta t$ given by

$$r_{T,ij}\ \Delta t. \tag{13}$$

In the simplest reference implementation, when a cell transforms, it retains its volume and position, and overwrites all other phenotype parameters from the new cell type. These parameters are summarized in **Table 6**.

| dictionary symbol | synonyms | controllable phenotype parameter |
|---|---|---|
| transform to X | transform to cell type X | $r_{T,ij}$ |
| fuse to X | fuse to cell type X | $r_{F,ij}$ |
| phagocytose dead cell | phagocytosis of dead cell, phagocytosis of dead cells | $r_{PD,i}$ |
| phagocytose X | phagocytose cell type X, phagocytosis of X | $r_{PL,ij}$ |
| attack X | attack cell type X | $r_{A,ij}$ |
| immunogenicity to X | immunogenicity to cell type X | $g_{A,ij}$ |
| damage rate | | $r_{\text{damage}}$ |

**Table 6.** Key cell-cell interaction terms (focused on transformation, fusion, phagocytosis, and effector attack) in the behavior dictionary, along with corresponding controllable parameters in the PhysiCell reference implementation. Here, *X* denotes any cell type.

##### *Fusion*

As a simple reference model of cell fusion, if cell $i$ is in contact with cell $j$, the cells can fuse between the current time $t$ and a future time $t + \Delta t$ with probability

$$r_{F,ij}\ \Delta t, \tag{14}$$

where $r_{F,ij}$ is the fusion rate for the cell $i$ to the type of cell $j$. In our reference model, when cells $i$ and $j$ fuse, the newly fused cell:

- is placed at the center of volume of the pre-fused cells $i$ and $j$
- combines the volumes of the cells $i$ and $j$
- combines the number of nuclei in cells $i$ and $j$ (if tracked)
- combines all internalized substrates in cells $i$ and $j$ (if tracked)

At the present time, there is no community consensus on what *phenotype parameters* should be acquired by the newly fused cell; future versions of the reference model may take a (volume-weighted) average of the pre-fused cells' phenotypes. Moreover, community consensus is required to determine if the fused cell should shrink towards a typical single cell size or retain its larger size in the long term. Behavioral parameters that can be modulated by our grammar are summarized in **Table 6**.

***Phagocytosis (or predation or ingestion)***

We include a simple reference model of phagocytosis (or predation or ingestion), based on recent modeling[4,5,8]: if cell $i$ is in contact with (live) cell $j$, then between the current time $t$ and a future time $t + \Delta t$, cell $i$ has a probability of phagocytosing cell $j$ given by:

$$r_{PL,ij}\,\Delta t, \tag{15}$$

where $r_{P,ij}$ is cell $i$'s rate of phagocytosing live cells of type $j$. Similarly, if cell $j$ is dead, then the probability of phagocytosing the cell in that time interval is $r_{PD,i}\,\Delta t$.

In the reference model, when a cell phagocytoses (or ingests) another, it absorbs all its volume, while retaining its original position. (In the event that a more detailed volume is represented as in prior work[13], cell $j$'s solid volume is added to cell $i$'s cytoplasmic solid volume, and cell $j$'s fluid volume is added to cell $i$'s overall fluid volume.) If the simulation framework actively regulates the volume of cell $i$ (e.g., to maintain a target volume[13]), then over time cell $i$ will return to its previous volume, as a simplified model of degradation of the phagocytosed cell materials. These parameters are summarized in **Table 6**.

***Effector attack***

Based on recent models[4,5,8,31,32], we define a reference model of cytotoxic effector attack (e.g., CD8 T inflicting fatal damage on a target cell via perforin and granzymes[33-35]). If cell i is in contact with cell j, then its probability of damaging cell j between time $t$ and a future time $t + \Delta t$ is given by

$$r_{A,ij} \cdot g_{ji}\,\Delta t, \tag{16}$$

where $r_{A,ij}$ is cell $i$'s rate of attacking cells of type $j$, and $g_{ji}$ is cell $j$'s immunogenicity to cell $i$. When cell $i$ attacks cell $j$, the damage $d_j$ of the cell increases by $r_{\text{damage}}\Delta t$ ($r_{damage}$ is the rate at which cell $i$ causes damage to cell $j$, taken to be 1 in the reference implementation), and the total attack time logged by cell $j$ is increased by $\Delta t$. Note that if multiple cells are attacking cell $j$ simultaneously, then both the damage and total attack time increase more rapidly. We note that in this reference model, effector attack increases damage in the target cell $j$, but it does not cause death without additional hypotheses relating damage to a death rate. Behavioral parameters that can be modulated by our grammar are summarized in **Table 6**.

***Other terms***

Because scientists may require custom biology not yet supported in our grammar or reference implementation, we allow a *custom* symbol to that can access any custom cell parameters. We also reserve symbols for cell-basement mechanical interactions that currently lack reference implementations. See **Table 7**.

| dictionary symbol | synonyms | controllable phenotype parameter |
|---|---|---|
| custom:X | custom: X, custom X | X |
| cell-BM adhesion | | None (symbol reserved) |
| cell-BM repulsion | | None (symbol reserved) |

**Table 7.** Other cell terms in the behavior dictionary, along with corresponding controllable parameters in the PhysiCell reference implementation. Here, X is a custom cell variable (parameter).

#### Signals

Signals are (typically exogeneous but sometimes internal) stimuli or information that can be interpreted by a cell to drive behavioral or state changes. In the context of mathematical modeling, signals are inputs to constitutive laws or agent rules. We broadly surveyed mathematical and biological models from cancer biology[36-47], tissue morphogenesis[38,48-52], immunology[4,5,8,36,53,54], and microbial ecosystems[55,56], to generalize classes of inputs to cell behavioral rules, generally including chemical factors, mechanical cues, cell volume (e.g., for volume-based cycle checkpoints), physical contact with cells, live/dead status, current simulation time (for use in triggering events), and accumulated damage (e.g., from effector attack[33-35]). We thus define the following forms of signals in **Table 8.**

| dictionary symbol | synonyms | accessible variable (and notes) |
|---|---|---|
| X | | extracellular diffusible substrate X at cell location |
| intercellular X | internalized X | total internalized substrate X in the cell |
| X gradient | grad(X), gradient of X | norm of the gradient of extracellular diffusible substrate X |
| volume | | total cell volume |
| pressure | | (nondimensionalized) mechanical pressure acting upon the cell |
| contact with Y | contact with cell type Y | number of live cells of type Y in physical contact with the cell |
| contact with live cell | contact with live cells | number of live cells in physical contact with the cell |
| contact with dead cell | contact with dead cells | number of dead cells in physical contact with the cell |
| contact with basement membrane | contact with BM | 1 if in contact with a basement membrane, and 0 otherwise (reserved symbol for future reference models) |
| damage | | total cell damage (see ***effector attack*** above) |
| total attack time | | total accumulated attack time on the cell by effector cells (see ***effector attack*** above) |
| dead | is dead | 1 if the cell is dead, and 0 otherwise |
| apoptotic | is_apoptotic | 1 if the cell is apoptotic, and 0 otherwise (or if the cell is undergoing a non-necrotic cell death) |
| necrotic | is_necrotic | 1 if the cell is necrotic, and 0 otherwise |
| time | current time, global time | current elapsed simulation time |
| custom:Z | custom: Z, custom Z | access to a cell's custom parameter Z |

**Table 8.** A list of symbols in our vocabulary of signals that can be used to drive behavioral or state changes in cells. Here, *X* is any diffusible substrate, *Y* is a cell type, and *Z* is a customized cell parameter (or variable).

Some signals were included at the request of the mathematical modeling community; for instance, the *time* signal can trigger global model events (e.g., exposure to a dose of radiation therapy), and *apoptotic* and *necrotic* allow more fine-grained control of dead cells. The *damage* signal is currently a flexible, generic term that could describe membrane, nuclear, mitochondrial, DNA, or other types of damage as needed by the specific context of the modeling application. The *total attack time* signal similarly allows "area-under-the-curve" (AUC) models that require total exposure time to attacking cells. The *custom* family of signals enables greater extensibility by allowing direct access to model-specific cell variables, without the need to formally extend the language and its vocabulary itself.

Over time, new signals can readily be added to the standard vocabulary based on community feedback, or when new signals emerge as widespread in modeling. For example, more specific types of damage (e.g., DNA damage or mitochondrial damage) may appear widely in some biological domains, thus justifying the addition of multiple, more specific damage signals in the vocabulary.

We note that while some signals can be found in existing ontologies (e.g., many diffusible chemical substrates in ChEBI[57,58], mechanical pressure in the ontology for physics in biology (OPB)[59] or non-specific pressure in PATO[60], or non-specific cell-cell contact in the cell behavior ontology (CBO)[1], no single ontology was found to incorporate our broader family of signals in a form that can be concretely and algorithmically generated for a specific simulation or mathematical model system.

### CELL BEHAVIOR GRAMMAR: FULL DESCRIPTION

Now that we have introduced the key signals and behaviors, we now describe our grammar to modulate cell behaviors in response to signals. Cell behavioral hypotheses are written as human-readable statements of the form:

In [cell type T]:

[signal S] [increases or decreases] [behavior B] [optional parameters 1 & 2]. [optional statements].

Here, [signal S] and [behavior B] are defined symbols defined above. This construction allows us to efficiently "bundle" multiple behavioral response statements for a cell type.

***Optional parameters 1: base and maximum effect***

These parameters are used to specify the maximum effect of the rule. They take the form:

from [base] towards [value].

Here, [base] is the unperturbed value of the behavioral parameter in the absence of signals, and [value] is its saturated response under maximum signal. The values should be stated with units, with a preference for microns for spatial units and minutes for time units.

When [base] and [effect] are not specified, parsers should assume a tenfold (one order-of-magnitude) change in behavior from the base parameter value:

$$p_M = 10\, p_0 \quad \text{for increasing responses}$$

$$p_m = 0.1\, p_0 \quad \text{for decreasing responses}$$

***Optional parameters 2: response form and parameters***

Any rule can specify the form of its response function (currently linear or Hill) by appending:

with a [linear or Hill] response.

If the rule further specifies the parameters of the response, it takes the form:

with a [linear or Hill] response, with [parameters].

For linear responses, [parameters] takes the form:

with minimum threshold [value] and maximum threshold [value].

For Hill responses, [parameters] takes the form:

with half-max [value] and Hill power [value].

***Optional statements: effect on dead cells***

By default, we assume that the hypothesis rules apply to live cells only, to avoid non-physical behaviors such as nonzero motility or proliferation of dead cells. However, rules can be designated to apply to dead cells in addition to live cells with the optional statement:

Rule applies to dead cells.

***Examples:***

Here are examples of behavioral response statements with optional arguments noted in red.

- Oxygen increases cycle entry.
- Oxygen increases cycle entry from 7e-6 1/min towards 7e-4 1/min.
- Oxygen increases cycle entry from 7e-6 1/min towards 0.0007 1/min with a Hill response, with half-max 21.5 mmHg and Hill power 4.

- Pressure decreases cycle entry towards 0 1/min with a linear response, with minimum threshold 0 and maximum threshold 0.5.
- doxorubicin increases apoptosis.
- doxorubicin increases apoptosis towards 0.01 1/min with a Hill response, with half-max 0.1 and Hill power 2.
- Virus increases fusion to tumor cells. Rule applies to dead cells.

As an example of bundling multiple statements for a single cell type:

In malignant epithelial cells:

- Oxygen increases cycle entry towards 0.0007 1/min, with a Hill response, with half-max 21.5 mmHg and Hill power 4.
- Pressure decreases cycle entry towards 0.0 1/min, with a linear response, with minimum threshold 0 and maximum threshold 0.5.
- Dead increases debris secretion towards 1 1/min, with a Hill response, with half-max 0.1 and Hill power 4. Rule applies to dead cells.

#### Mathematical representation

With clearly defined behaviors and signals, and grammar to connect them, we can now uniquely map human-interpretable cell hypothesis statements onto mathematical expressions that make the grammar both human interpretable and computable. Moreover, our mathematical formulation allows new hypotheses to be directly added to models without modifying prior hypotheses, making our framing extensible and scalable as new knowledge is acquired.

##### *Response functions*

We use *response functions* to mathematically represent how a behavior varies with a signal towards its maximal (saturated) response. Our initial set of supported response functions are drawn from recurring forms in mathematical biology. In general, a response function $R$ satisfies these properties:

| | |
|---|---|
| 1. $R(0) = 0$. | In the absence of a signal, there is no response. |
| 2. $R'(s) \geq 0$ | Responses increase monotonically as the signals increases. |
| 3. $R(s) \to 1$ as $s \to \infty$ | The response saturates at 100% for large amounts of signal. |
| 4. $R(s) = 0$ if $s < 0$ | For convenience, negative signals are ignored. |

###### <u>Linear response function</u>

Many mathematical models use linear constitutive relations, so we define:

$$L(s; s_0, s_1) = \begin{cases} 0 & \text{if } s \leq s_0 \\ \dfrac{s - s_0}{s_1 - s_0} & \text{if } s_0 < s < s_1 \\ 1 & \text{if } s \geq s_1 \end{cases}$$

Here, $s_0$ and $s_1$ are (nonnegative) parameters of the response function governing where the response reaches its minimum and maximum values. For simplicity, we will call $s_0$ and $s_1$ the minimum and maximum response thresholds, respectively.

###### <u>Hill (sigmoidal) response function</u>

Hill functions (or sigmoidal functions) are commonly used in pharmacodynamics and in computational models of cell responses to chemical signals. We define:

$$H(s; s_{\text{half}}, \text{h}) = \frac{s^h}{s_{\text{half}}^h + s^h} = \frac{\left(\frac{s}{s_{\text{half}}}\right)^h}{1 + \left(\frac{s}{s_{\text{half}}}\right)^h} \text{ if } s \geq 0, \quad \text{and } H(s) = 0 \text{ if } s < 0.$$

Here, $s_{\text{half}}$ and $h$ are (nonnegative) parameters governing where the response reaches half of its maximum value ($s_{\text{half}}$) and the steepness of the response (the Hill power $h$).

**Multivariate Hill (sigmoidal) response function**

Sometimes, we may need to consider the impact of multiple signals $\boldsymbol{s}$ (with half-maximum parameters $\boldsymbol{s}_{\text{half}}$ and Hill powers $\boldsymbol{h}$) on a behavior. We therefore introduce a multivariate version:

$$H_M(\boldsymbol{s}; \boldsymbol{s}_{\text{half}}, \boldsymbol{h}) = \frac{\sum_i \left(\frac{s_i}{s_{\text{half},i}}\right)^{h_i}}{1 + \sum_i \left(\frac{s_i}{s_{\text{half},i}}\right)^{h_i}},$$

with the additional stipulation that we replace $s_i$ by $(s_i)^+ = \max(s_i, 0)$ as needed. It satisfies:

- $H_M(\boldsymbol{0}) = 0$
- $H_M(0, \dots, 0, s_i, 0, \dots 0) = H(s_i)$ for any $i$
- $0 \leq H_M \leq 1$ for all $\boldsymbol{s}$
- $H_M(\boldsymbol{s}) \to 1$ as $|\boldsymbol{s}| \to \infty$

Thus, the signals can contribute to a response individually and in combination, and when only one signal is supplied, the response behaves as the more common Hill response function. This functional form has the benefit that individual Hill response parameters do not need to be recalibrated as new arguments (i.e., new biological knowledge and rules) are added. See **Fig. S1 (left).**

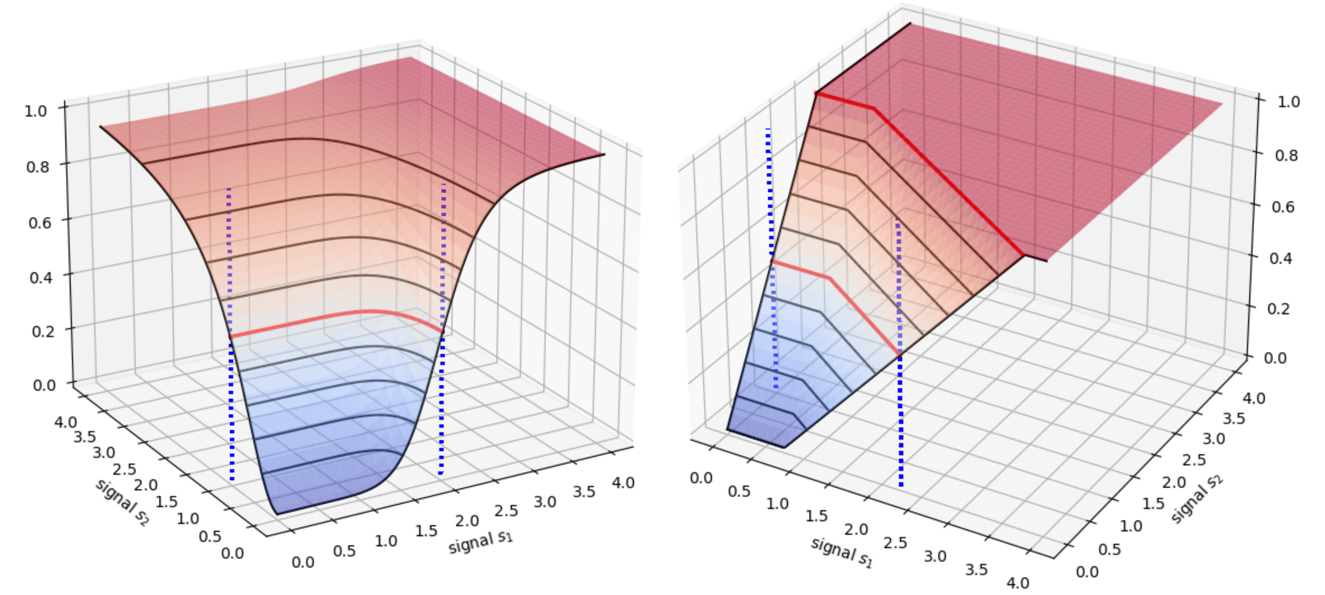


**Fig. S1: Multivariate response functions.** ***Left:*** A multivariate Hill response function, where $s_1$ has half-max 2 and Hill power 8, and $s_2$ has half-max 1 and Hill power 2. Half-maxes are plotted on each variable as blue dashed lines, while the half-max across the multivariate function is plotted as a red contour. $H(s_1)$ and $H(s_2)$ are plotted as black curves along the respective axes. ***Right:*** A multivariate linear response function, where $s_1$ has min and max thresholds 0.75 and 3.75, and $s_2$ has thresholds 0 and 2. Half-maxes are plotted on each variable as blue dashed lines, while the half-max across the multivariate function is plotted as a red contour. $L(s_1)$ and $L(s_2)$ are plotted as black curves along the respective axes.

#### Multivariate linear response function

Similarly, we may need to consider the impact of multiple signals $\boldsymbol{s}$ (with thresholds $\boldsymbol{s}_0$ and $\boldsymbol{s}_1$ ) on a behavior. Motivated by the properties of the multivariate Hill function, we introduce a multivariate linear response function:

$$L_M(\boldsymbol{s}; \boldsymbol{s}_0, \boldsymbol{s}_1) = \min\left(1, \sum_i L\left(s_i; s_{0,i}, s_{1,i}\right)\right),$$

with the additional stipulation that we replace $s_i$ by $(s_i)^+ = \max(s_i, 0)$ as needed. It satisfies:

- $L(\mathbf{0}) = 0$
- $L_M(0, \ldots, 0, s_i, 0, \ldots 0) = L(s_i)$ for any $i$
- $0 \leq L_M \leq 1$ for all $\boldsymbol{s}$
- $L_M(\boldsymbol{s}) \to 1$ as $|\boldsymbol{s}| \to \infty$

Thus, the signals can contribute to a response individually and in combination, and when only one signal is supplied, the response behaves as the simpler linear response function. This functional form has the benefit that individual linear response parameters do not need to be recalibrated as new arguments (i.e., new biological knowledge and rules) are added. See **Fig. S1 (right).**

#### Approximating linear responses with Hill response functions

When curating and aggregating prior knowledge, we may encounter linear relationships and seek to approximate them Hill response functions for a more unified framework. If a linear response function has thresholds $s_0$ and $s\_1$, then we stipulate that:

- The half-max is $s_{\text{half}} = \frac{1}{2}(s_0 + s_1)$
- The Hill response function reaches 90% of its peak value at $s_1$

This can be satisfied when the Hill power is given by

$$h = \frac{\log\left(\frac{1-\epsilon}{\epsilon}\right)}{\log\left(\frac{s_1}{s_{\text{half}}}\right)},$$

where $0 < \epsilon < 1$ is a tolerance. (We set $\epsilon = 0.1$.) An Example is shown **Fig. S2**. We note that computationally, integer-valued Hill powers are at least tenfold faster in execution than non-integer powers, so it can be advantageous to round to the nearest whole number.

#### Approximating Hill responses with linear response functions

Similarly, when curating and aggregating prior knowledge, we may encounter Hill relationships and seek to approximate them as linear responses in a broader, unified framework. Moreover, this offers an opportunity for computational acceleration. If a Hill response function has half-max $s_{\text{half}}$ and Hill power $h$, then we stipulate that:

- The right threshold $s_1$is reached where the Hill response function reaches 90% of its peak value, and
- The half-max and thresholds are related by $s_{\text{half}} = \frac{1}{2}(s_0 + s_1)$

This can be satisfied by setting:

$$s_1 = s_{\text{half}}\left(\frac{1-\epsilon}{\epsilon}\right)^{\frac{1}{h}}$$

and

$$s_0 = 2\, s_{\mathrm{half}} - s_1.$$

As before, $0 < \epsilon < 1$ is a tolerance, which we set to $\epsilon = 0.1$. An Example is show in **S2.**

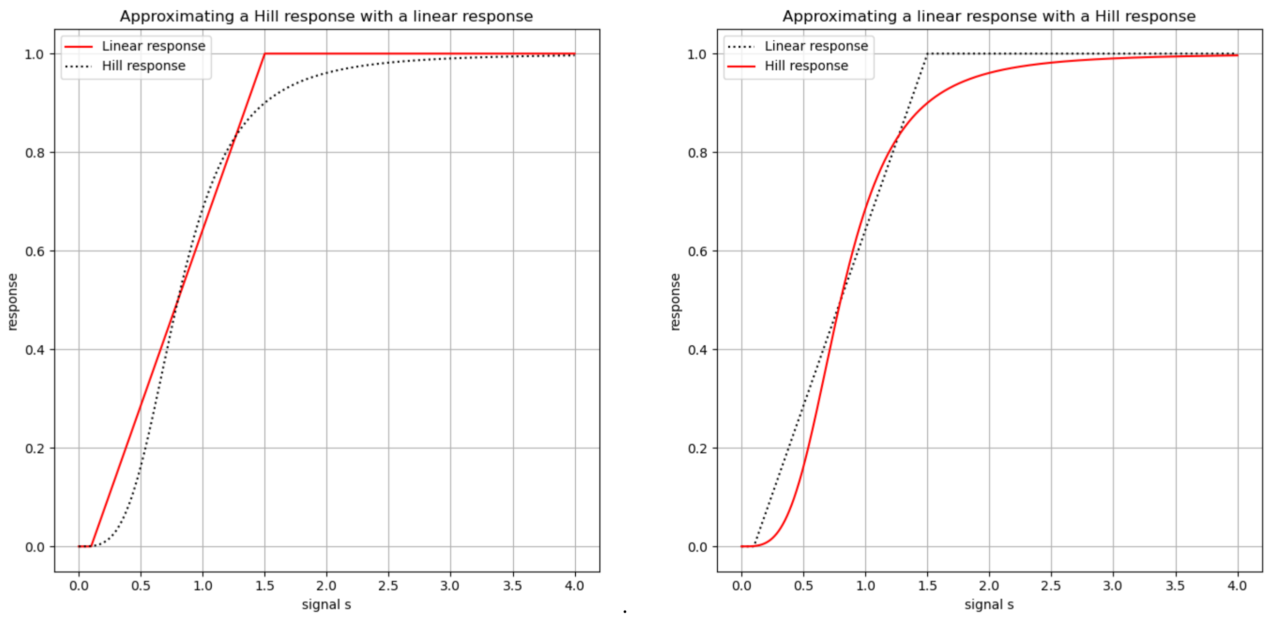


**Fig. S2: Converting between linear and Hill response functions.** ***Left:*** Approximating a linear response function (black dotted curve) with a Hill response function (red). ***Right:*** Approximating a Hill response function (black dotted) with a linear response function (red).

##### *Single hypothesis statement: S increases B or S decreases B*

For any biological hypothesis statement of the form "*S* increases *B*" or "*S* decreases *B*", the behavior *B* can be associated with a phenotypic behavioral parameter $p$. The statement then takes the form of a linear interpolation in the nonlinear response function $R$:

$$p(s) = p_0 + (p_{\mathrm{M}} - \mathrm{p}_0)R(s) = \big(1 - R(s)\big)\cdot p_0 + R(s)\cdot p_M$$

Here, $p_0$ is the base level of the parameter (from the base behavioral phenotype), $p_M$ is the maximal (saturated) response, and $R(s)$ is a user-selected response function.

**Example: oxygen-driven cycling**

The statement "oxygen increases cycle entry" (with a "base" value of $p_0 = 0.001\ \mathrm{hr}^{-1}$ and a maximum rate of $p_{\mathrm{M}} = 0.042\ \mathrm{hr}^{-1}$) takes the form

$$r_{01} = 0.001 + (0.042 - 0.001)R(\mathrm{pO}_2).$$

If we use a linear response function as in prior work[6,7,10,13,21] with minimum and maximum thresholds at 5 mmHg and 38 mmHg, respectively, then the response function takes the form

$$r_{01} = \begin{cases} 0.001 & \text{if } \mathrm{pO}_2 < 5\ \mathrm{mmHg} \\ 0.001 + (0.042 - 0.001)\left(\dfrac{\mathrm{pO}_2 - 5}{38 - 5}\right) & \text{if } 5 \le \mathrm{pO}_2 \le 38\ \mathrm{mmHg} \\ 0.042 & \text{if } \mathrm{pO}_2 > 38\ \mathrm{mmHg} \end{cases}$$

If we use a Hill response function (with a half-max of 21.5 mmHg, and Hill power of 4), the mathematical rule takes the form

$$r_{01} = 0.001 + (0.042 - 0.001)\frac{(\mathrm{pO}_2)^4}{21.5^4 + (\mathrm{pO}_2)^4}$$

**Example: mechanoregulation of cycling**

The statement "pressure decreases cycle entry" (with a "base" value of $p_0 = 0.001\ \mathrm{hr}^{-1}$ and a maximally inhibited rate of $p_{\mathrm{M}} = 0\ \mathrm{hr}^{-1}$) takes the form

$$r_{01} = 0.001 + (0 - 0.001)R(p).$$

If we use a Hill response function (with a half-max of 0.25, and Hill power of 3), the mathematical rule takes the form

$$r_{01} = 0.001 + (0.0 - 0.001)\frac{p^3}{0.25^3 + p^3} = 0.001\left(1 - \frac{p^3}{0.25^3 + p^3}\right)$$

***Multiple hypothesis statements: $S_1$ increases B, $S_2$ increases B***

For any biological hypothesis statement of the form "*S* decreases *B*", the behavior *B* can be associated with a phenotypic behavioral parameter $p$. The statement then takes the form:

$$p(s_1, s_2) = p_0 + (p_{\mathrm{M}} - p_0)R(s_1, s_2)$$

Here, $p_0$ is the base level of the parameter (its value in the absence of any signals), $p_M$ is the maximal (saturated) response, and $R(s)$ is a user-selected response function. We use the multivariate Hill response function $H_M$ for $R(\boldsymbol{s})$.

**Example: oxygen- and hormone-dependent cycling**

The statements "oxygen increases cycle entry" and "estrogen increases cycle entry" (with a "base" value of $p_0 = 0.001\ \mathrm{hr}^{-1}$and a maximum rate of $p_{\mathrm{M}} = 0.042\ \mathrm{hr}^{-1}$) takes the form

$$r_{01} = 0.001 + (0.042 - 0.001)\ H_M(\mathrm{pO_2}, e).$$

If we use an extended Hill response function (where oxygen has a half-max of 21.5 mmHg and Hill power of 4, and nondimensionalized estrogen has a half-max of 0.5 and Hill power of 3), the mathematical rule takes the form

$$r_{01} = 0.001 + (0.042 - 0.001)\frac{\left(\frac{\mathrm{pO_2}}{21.5}\right)^4 + \left(\frac{e}{0.5}\right)^3}{1 + \left(\frac{\mathrm{pO_2}}{21.5}\right)^4 + \left(\frac{e}{0.5}\right)^3}.$$

***Multiple competing hypothesis statements: S1 increases B, S2 decreases B***

For competing biological hypothesis statements of the form "$S_1$ increases *B*" and "$S_2$ decreases *B*", if the behavior has the associated phenotypic parameter $p$, then we use the form:

$$p(s_1, s_2) = \left(p_0 + (p_M - p_0)\frac{s_1^p}{(s_1^*)^p + s_1^p}\right) + \left(p_m - \left(p_0 + (p_M - p_0)\frac{s_1^p}{(s_1^*)^p + s_1^p}\right)\right)\frac{s_2^q}{(s_2^*)^q + s_2^q}$$

Here, $p_0$ is the base level of the parameter (from the base behavioral phenotype in the absence of signals), $p_M$ is the maximal response to the promoting signal $s_1$ (with half-max $s_1^*$ and Hill power $p$), and $p_m$ is the maximal response to the inhibiting signal $s_2$ (with half-max $s_2^*$ and Hill power $q$).

We can write this more simply with:

$$U = \frac{s_1^p}{(s_1^*)^p + s_1^p} \quad , \quad D = \frac{s_2^q}{(s_2^*)^q + s_2^q}$$

giving the overall response as a bilinear interpolation:

$$p(s_1, s_2) = (1 - D) \cdot [\,(1 - U) \cdot p_0 + U \cdot p_M] + D \cdot p_m$$

In this formulation, the promoting signal $s_1$ pushes the behavior towards a "target" value that is then subject to inhibition by $s_2$. Notice that:

- If $s_2 = 0$, then $D = 0$ and $p(s_1, s_2)$ behaves as the regular Hill response function $H(s_1)$ for the promoting signal.
- If $s_1 = 0$, then $U = 0$ and $p(s_1, s_2)$ behaves as the regular Hill response function $H(s_2)$ for the inhibiting signal.
- In the absence of either signal, $p(0,0) = p_0$.
- As $s_1 \to \infty$ and $s_2 \to \infty$, then $p(s_1, s_2) \to p_m$.

**Example: oxygen-driven cycling with negative mechanofeedback**

Suppose "oxygen increases cycle entry" and "pressure decreases cycle entry" with parameters as in the prior examples. Then our functional form is:

$$U = \frac{(\mathrm{pO}_2)^4}{21.5^4 + (\mathrm{pO}_2)^4}, \qquad D = \frac{p^3}{0.25^3 + p^3}$$

$$p(\mathrm{pO}_2, p) = (1 - D) \cdot [(1 - U) \cdot 0.001 + U \cdot 0.042] + D \cdot 0$$

See **Fig. S3 (left)**.

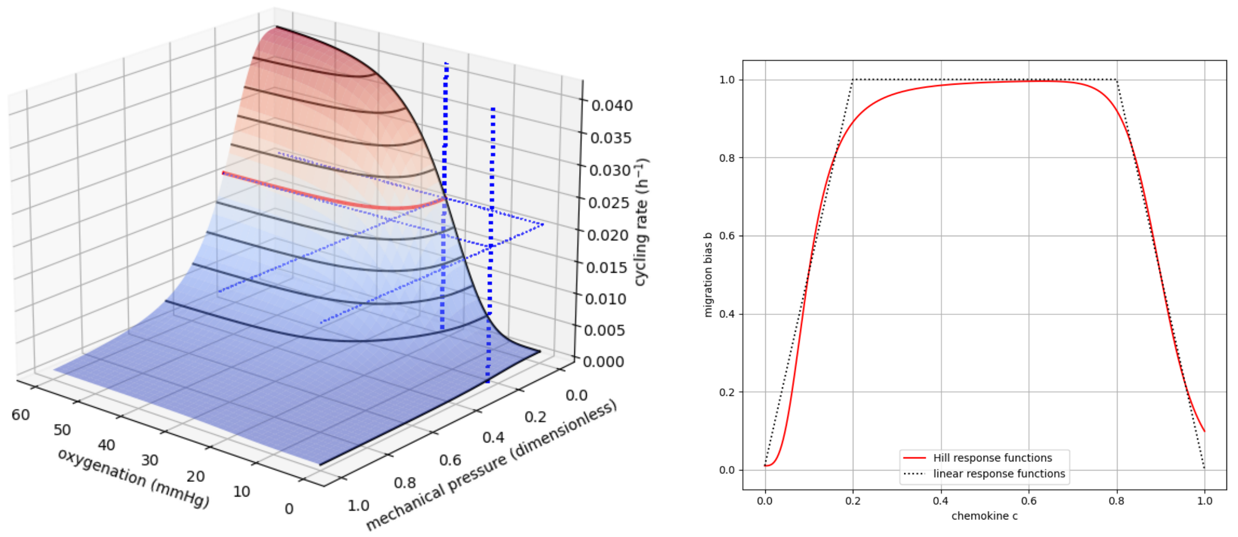


**Fig. S3: Sample response functions.** ***Left:*** The behavioral response function for the statements "oxygen increases cycle entry" and "pressure decreases cycle entry", using the multivariate Hill response function. ***Right:*** A non-monotonic response function from the statements "*c* increases migration bias" (for lower values of *c*) and "*c* decreases migration bias" (for higher values of *c*), showing both linear and Hill response constructions.

**Example: "contradictory" observations on cell migration**

Suppose a cell is migrating by chemotaxis towards a (scaled to be nondimensional) chemokine $c$, where we observe:

- Migration becomes less random and more directed as the chemokine concentration increases (it is more difficult to sense a chemical gradient in low concentrations)
- Migration becomes more random and less directed for extremely high chemokine concentrations (high concentrations can saturate the cell's chemical receptors, making it difficult to sense the chemical gradient)

In the cell behavior grammar, these statements become:

- $c$ increases migration bias
  - We will use a base migration bias of 0.01 and maximum bias of 1.
  - If the effect reaches its maximum at $c = 0.2$, we set the linear response minimum and maximum thresholds to $s_0 = 0$ and $s_1 = 0.2$. For the Hill response approximation, we set $s_{\text{half}} = 0.1$ and $h = 3$
- $c$ decreases migration bias
  - We will use a minimum migration bias of 0.
  - If the effect is first notable at $c = 0.8$ and increases towards its maximum at $c = 1$, we use minimum and maximum response thresholds of $s_0 = 0.8$ and $s_1 = 1$ for a linear response. For the Hill response approximation, we set $s_{\text{half}} = 0.9$ and $h = 21$

These seemingly contradictory statements jointly create non-monotonic behavior:

$$b(c) = \big(1 - H(c; 0.9,21)\big) \cdot \big[\big(1 - H(c; 0.1,3)\big) \cdot 0.01 + H(c; 0.1,3) \cdot 1\big] + H(c; 0.9,21) \cdot 0$$

We plot the overall response using both Hill responses (red) and linear response (dotted black) in **Fig. S3 (right).**

***General form: multiple promoting and inhibiting rules***

Suppose we have the statements:

- $u_1$ increases B (with half-max $u_1^*$ and Hill power $p_1$)
- $u_2$ increases B (with half-max $u_2^*$ and Hill power $p_2$)
- ...
- $u_m$ increases B (with half-max $u_m^*$ and Hill power $p_m$)
- $d_1$ decreases B (with half-max $d_1^*$ and Hill power $q_1$)
- $d_2$ decreases B (with half-max $d_2^*$ and Hill power $q_2$)
- …
- $d_n$ decreases B (with half-max $d_m^*$ and Hill power $q_n$)

Here, let $p_M$be the maximum value of the parameter $p$ (under the combined influence of the up-regulating signals $\boldsymbol{u}$ ), let $p_0$ be its base value in the absence of signals, and let $p_m$ be its minimum value (under the combined influence of the down-regulating signals $\boldsymbol{d}$ ).

We define the total up response as:

$$U = H_M(\boldsymbol{u}; \boldsymbol{u}_{\text{half}}, \boldsymbol{p}) = \frac{\left(\frac{u_1}{u_1^*}\right)^{p_1} + \left(\frac{u_2}{u_2^*}\right)^{p_2} + \cdots + \left(\frac{u_m}{u_m^*}\right)^{p_m}}{1 + \left(\frac{u_1}{u_1^*}\right)^{p_1} + \left(\frac{u_2}{u_2^*}\right)^{p_2} + \cdots + \left(\frac{u_m}{u_m^*}\right)^{p_m}}$$

and the total down response as:

$$D = H_M(\boldsymbol{d}; \boldsymbol{d}_{\text{half}}, \boldsymbol{q}) = \frac{\left(\frac{d_1}{d_1^*}\right)^{q_1} + \left(\frac{d_2}{d_2^*}\right)^{q_2} + \cdots + \left(\frac{d_n}{d_n^*}\right)^{q_n}}{1 + \left(\frac{d_1}{d_1^*}\right)^{q_1} + \left(\frac{d_2}{d_2^*}\right)^{q_2} + \cdots + \left(\frac{d_n}{d_n^*}\right)^{q_n}}.$$

Using these, we combine the overall response of the behavioral parameter as bilinear interpolation int eh nonlinear up- and down-responses $U$ and $D$:

$$p(\boldsymbol{u}, \boldsymbol{d}) = (1 - D) \cdot [(1 - U) \cdot p_0 + U \cdot p_M\,] + D \cdot p_m$$

Notice that:

- In the presence of up-regulating signals only, this reduces to the multivariate Hill response function $H_M(\boldsymbol{u};\boldsymbol{u}^*,\boldsymbol{p})$.
- In the presence of down-regulating signals only, this reduces to the multivariate Hill response function $H_M(\boldsymbol{d};\boldsymbol{d}^*,\boldsymbol{q})$
- Generally, the combined up-regulating signals sets a "target" value of the parameter, which can then be inhibited by the combined down-regulating signals.

Note also that adding and removing individual rules to the form does not require alteration to the remaining rules. In this release, we use multivariate Hill response functions for clarity, but mixed linear and Hill responses could be used in the future.

**Example:**

We combine three hypothesis statements from prior examples that modulate cycle entry:

- Oxygen increases cycle entry
- Estrogen increases cycle entry
- Pressure decreases cycle entry

The combined mathematical form for the cycle entry rate $r_{01}$ is:

$$U = \frac{\left(\frac{\mathrm{pO_2}}{21.5}\right)^4 + \left(\frac{\mathrm{e}}{0.5}\right)^3}{1 + \left(\frac{\mathrm{pO_2}}{21.5}\right)^4 + \left(\frac{\mathrm{e}}{0.5}\right)^3} \quad , \quad D = \frac{\left(\frac{p}{0.25}\right)^3}{1 + \left(\frac{p}{0.25}\right)^3}$$

$$r_{01} = (1 - D) \cdot [p_0 + (p_M - p_0)] + D \cdot p_m$$

$$p_0 = 0.001\ \mathrm{hr}^{-1} \quad , \quad p_M = 0.042\ \mathrm{hr}^{-1} \quad , \quad p_m = 0\ \mathrm{hr}^{-1}.$$

#### Reference PhysiCell implementation

We implemented support for the grammar in PhysiCell version 1.12.0[61]. Rules are imported at the start of a simulation using a compact CSV format (see supplementary information), parsed, and stored for each cell type. Rules are stored in a ruleset data structure, with a separate ruleset for each cell definition:

1. For each modulated behavior (with corresponding parameter $p$), store a set of rules:
    a. For each rule, we store:
        i. The "direction" of the response (increases/promotes or decreases/inhibits the behavior)
        ii. The signal used in the rule
        iii. The half-max, Hill parameter, and maximal value of the behavior of the parameter
    b. Based upon all the up-regulating rules, store the largest maximum parameter value $p_M$
    c. Based upon all the down-regulating rules, store the lowest minimum parameter value $p_m$

To execute a set of rules (for a single cell):

1. For each modulated behavior (with corresponding parameter $p$ ):
    a. For each rule:
        i. Sample the corresponding signal $s$
        ii. Query its Hill parameter $h$ and half-max $s_{\mathrm{half}}$
        iii. Compute $\left(\frac{s}{s_{\mathrm{half}}}\right)^h$
        iv. If the rule increases (promotes) the behavior, add the contribution from iii to $U$ in the generalized multivariate response function. Otherwise, add the contribution to $D$.

b. Query the base parameter value $p_0$, the maximum parameter value $p_M$, and the minimum parameter value $p_m$.
c. Compute the modulated parameter value via $p = (1 - D) \cdot [(1 - U) \cdot p_0 + U \cdot p_M] + D \cdot p_m$
d. Update the phenotypic parameter $p$ for the cell.

At the start of each simulation step, each cell evaluates its individual rules (based on its current cell type) as noted above to set its current phenotype parameters, and then runs its standard phenotype processes.

We envision that other agent-based simulation frameworks (e.g., Chaste, CompuCell3D, Biocellion, Morpheus, and Tissue Simulation Toolkit) can independently implement this framework so long as they (1) implement the reference cell process models, (2) define cell types, (3) can generate compatible dictionaries of signals and behaviors, (4) can parse the rules, and (5) use these to modulate the cell behavioral parameters using the reference multivariate response functions as noted above. For an up-front development cost, simulation frameworks could advance reproducibility and support cross-model validation.

### FULL MODEL ANNOTATIONS

#### Example 1: Model the progression of hypoxia in a metastatic tumor

| AGENT TYPE | BASE BEHAVIORS | HYPOTHESIS RULES |
|---|---|---|
| Tumor cell<br>(malignant epithelial cell) | Consume oxygen<br><br>Adhere to one another and exert mechanical forces<br><br>Proliferate as much as physical conditions will allow | Oxygen increases cycle entry<br><br>Pressure decreases cycle entry<br><br>Oxygen decreases necrosis<br><br>Oxygen decreases transformation to motile malignant epithelial cells |
| Motile tumor cell<br>(motile malignant epithelial cell) | Consume oxygen<br><br>Adhere to one another and exert mechanical forces<br><br>Proliferate as much as physical conditions will allow | Oxygen increases cycle entry<br><br>Pressure decreases cycle entry<br><br>Oxygen decreases necrosis<br><br>Oxygen increases transformation to non-motile malignant epithelial cells |

#### Example 2: Forecast locations of tumor cell invasion from an initial state defining tumor cell phenotypes with spatial transcriptomics

| AGENT TYPE | BASE BEHAVIORS | HYPOTHESIS RULES |
|---|---|---|
| Non-hypoxic (pre-hypoxic) cells | Consume oxygen<br><br>Adhere to one another and exert mechanical forces<br><br>Proliferate as much as physical conditions will allow | Oxygen increases cycle entry<br><br>Pressure decreases cycle entry<br><br>Oxygen decreases necrosis |
| Hypoxic cells | Consume oxygen<br><br>Adhere to one another and exert mechanical forces<br><br>Proliferate as much as physical conditions will allow | Oxygen increases cycle entry<br><br>Pressure decreases cycle entry<br><br>Oxygen decreases necrosis<br><br>Oxygen increases transformation to post-hypoxic cell |
| Post-hypoxic cells | Consume oxygen<br><br>Adhere to one another and exert mechanical forces<br><br>Proliferate as much as physical conditions will allow | Oxygen increases cycle entry<br><br>Pressure decreases cycle entry<br><br>Oxygen decreases necrosis |

### Example 3: Tumor attackers and tumor defenders

| AGENT TYPE | BASE BEHAVIORS | HYPOTHESIS RULES |
|---|---|---|
| Malignant epithelial cells | Consume oxygen<br>Proliferation<br>Apoptosis<br>Adhesion | Oxygen increases cycle entry<br>Pressure decreases cycle entry<br>Oxygen decreases necrosis<br>Damage increases apoptosis<br>Death increases debris secretion |
| Macrophages | Consume oxygen<br>Chemotaxis towards debris<br>Consume debris<br>Phagocytose dead cells<br>Exertion of mechanical forces | Contact with dead cell increases pro-inflammatory secretion<br>Oxygen increases pro-inflammatory factor secretion<br>Oxygen decreases anti-inflammatory factor secretion |
| CD8 T cells | Consume oxygen<br>Chemotaxis towards pro-inflammatory factor<br>Consume pro-inflammatory factor<br>Consume anti-inflammatory factor<br>Attack malignant epithelial cells<br>Exertion of mechanical forces | Pro-inflammatory factor increases attack of malignant epithelial cells<br>Anti-inflammatory factor decreases attack of malignant epithelial cells.<br>Contact with tumor cells decreases migration speed |

#### Example 4: T cell activation, expansion, and exhaustion in a diverse tumor microenvironment

| AGENT TYPE | BASE BEHAVIORS | HYPOTHESIS RULES |
|---|---|---|
| Malignant epithelial cells | Consume oxygen<br>Proliferation<br>Apoptosis<br>Adhesion<br>Exertion of mechanical forces<br>Randomly motile | Oxygen increases cycle entry<br>Pressure decreases cycle entry<br>Oxygen decreases necrosis<br>Damage increases apoptosis<br>Dead increases debris secretion<br>IFN-gamma decreases migration speed |
| M0 Macrophages | Consume oxygen<br>Chemotaxis towards debris<br>Consume debris<br>Phagocytose dead cells<br>Exertion of mechanical forces | Contact with dead cell increases transform to M1 macrophage<br>Contact with dead cell decreases migration speed |
| M1 Macrophages | Consume oxygen<br>Chemotaxis towards debris<br>Consume debris<br>Phagocytose dead cells<br>Exertion of mechanical forces<br>Secrete IFN-gamma (pro-inflammatory)<br>Consume IFN-gamma ß due to receptor-ligand binding<br>Transform to M2 macrophage | Contact with dead cell decreases migration speed<br>Oxygen decreases transformation to M2 (hypoxia increases transformation)<br>IFN-gamma increases cycle entry<br>IFN-gamma increases phagocytosis of dead cells |
| M2 Macrophages | Consume oxygen<br>Chemotaxis towards debris<br>Consume debris<br>Phagocytose dead cells<br>Exertion of mechanical forces<br>Secrete IL-10 (anti-inflammatory)<br>Consume IFN-gamma ß due to receptor-ligand binding | Contact with dead cell decreases migration speed<br>IFN-gamma decreases proliferation<br>IFN-gamma increases phagocytosis of dead cells |
| Naive CD8 T cells | Consume oxygen<br>Chemotaxis towards IFN-gamma<br>Consume IFN-gamma<br>Consume IL-10<br>Exertion of mechanical forces<br>Transform into CD8 T cell | IL-10 decreases transform to CD8 T cell<br>IFN-gamma increases transform to CD8 T cell |
| Effector CD8 T cells | Consume oxygen<br>Chemotaxis towards IFN-gamma<br>Consume IFN-gamma<br>Consume IL-10<br>Exertion of mechanical forces<br>Transform into CD8 T cell<br>Attack malignant epithelial cells | IFN-gamma Pro-inflamatory factor increases proliferation<br>IL-10 decreases attack malignant epithelial cells<br>IL-10 decreases migration speed<br>IL-10 increases transform to exhausted T cell<br>Contact with tumor cell decreases migration speed |
| Exhausted CD8 T cells | Consume oxygen<br>Very slow Chemotaxis towards IFN-gamma<br>Consume IFN-gamma<br>Consume IL-10 | |

#### Example 5: Using experimental insight to model combination immune-targeted therapies in the pancreatic cancer microenvironment

| AGENT TYPE | BASE BEHAVIORS | HYPOTHESIS RULES |
|---|---|---|
| PD-L1lo tumor cell | Slightly randomly motile but does not actively migrate<br>Consumes oxygen<br>Does not secrete anything | Oxygen increases cycle entry<br>Pressure decreases cycle entry<br>Oxygen decreases cell death<br>Cell damage leads to apoptosis<br>Dead cells emit debris |
| PD-L1hi tumor cell | Slightly randomly motile but does not actively migrate<br>Consumes oxygen<br>Does not secrete anything | Oxygen increases cycle entry<br>Pressure decreases cycle entry<br>Oxygen decreases cell death<br>Cell damage leads to apoptosis<br>Dead cells emit debris |
| Macrophage | Chemotaxes towards debris secreted by dead tumor cells<br>Makes pro- and anti-inflammatory factor in different ratios depending on oxygen availability<br>(more hypoxic = more tumor-favorable behavior) | Secretes pro-inflammatory factor in the presence of oxygen<br>Resistant case: max. PIF secretion is 1 unit<br>Responsive case: max. PIF is 5 units<br>Oxygen decreases anti-inflammatory factor secretion (secretes AIF in hypoxic conditions) |
| PD-1hi CD137lo CD8 T Cell | Extremely low killing ability compared to other CD8 T Cells | Contact with PD-L1hi tumor cell decreases migration speed ("sticks") |
| PD-1lo CD137lo CD8 T Cell | Good at killing tumor cells<br>Better at killing PD-L1lo than PD-L1hi by a factor of 10<br>Does not make its own PIF<br>Chemotaxes to PIF | Anti-inflammatory factor decreases tumor attack rate<br>Pro-inflammatory factor increases tumor attack rate<br>Contact with PD-L1hi tumor decreases migration speed<br>AIF decreases migration speed |
| PD-1hi CD137hi CD8 T cell | Not good at tumor killing<br>Makes PIF<br>Chemotaxes toward PIF | Contact with PD-L1hi tumor decreases migration speed |
| PD-1lo CD137hi CD8 T Cell | Good at killing tumor<br>Better at killing PD-L1lo than PD-L1hi by a factor of 10<br>Makes PIF<br>Chemotaxes toward PIF | Anti-inflammatory factor decreases tumor attack rate<br>Pro-inflammatory factor increases tumor attack rate<br>Contact with PD-L1hi tumor decreases migration speed<br>AIF decreases migration speed |
| PD-1hi CD4 T Cell | Makes PIF | AIF decreases migration speed |
| PD-1lo CD4 T Cell | Makes PIF | AIF decreases migration speed |

#### References


1. Sluka, J.P., Shirinifard, A., Swat, M., Cosmanescu, A., Heiland, R.W., and Glazier, J.A. (2014). The cell behavior ontology: describing the intrinsic biological behaviors of real and model cells seen as active agents. Bioinformatics *30*, 2367-2374. 10.1093/bioinformatics/btu210.
2. Friedman, S.H., Anderson, A.R.A., Bortz, D.M., Fletcher, A.G., Frieboes, H.B., Ghaffarizadeh, A., Grimes, D.R., Hawkins-Daarud, A., Hoehme, S., Juarez, E.F., et al. (2016). MultiCellDS: a community-developed standard for curating microenvironment-dependent multicellular data. bioRxiv **[preprint]** *090456*. 10.1101/090456.
3. Sundus, A., Kurtoglu, F., Konstantinopoulos, K., Chen, M., Willis, D., Heiland, R., and Macklin, P. (2022). PhysiCell training apps: Cloud hosted open-source apps to learn cell-based simulation software. bioRxiv **[preprint]** *10.1101/2022.06.24.497566*. 10.1101/2022.06.24.497566.
4. Jenner, A.L., Smalley, M., Goldman, D., Goins, W.F., Cobbs, C.S., Puchalski, R.B., Chiocca, E.A., Lawler, S., Macklin, P., Goldman, A., and Craig, M. (2022). Agent-based computational modeling of glioblastoma predicts that stromal density is central to oncolytic virus efficacy. iScience *25*, 104395. 10.1016/j.isci.2022.104395.
5. Islam, M.A., Getz, M., Macklin, P., and Versypt, A.N.F. (2022). An agent-based modeling approach for lung fibrosis in response to COVID-19. bioRxiv **[preprint]**. 10.1101/2022.10.03.510677.
6. Wang, Y., Brodin, E., Nishii, K., Frieboes, H.B., Mumenthaler, S.M., Sparks, J.L., and Macklin, P. (2021). Impact of tumor-parenchyma biomechanics on liver metastatic progression: a multi-model approach. Sci Rep *11*, 1710. 10.1038/s41598-020-78780-7.
7. Rocha, H.L., Godet, I., Kurtoglu, F., Metzcar, J., Konstantinopoulos, K., Bhoyar, S., Gilkes, D.M., and Macklin, P. (2021). A persistent invasive phenotype in post-hypoxic tumor cells is revealed by fate mapping and computational modeling. iScience *24*, 102935. 10.1016/j.isci.2021.102935.
8. Getz, M., Wang, Y., An, G., Asthana, M., Becker, A., Cockrell, C., Collier, N., Craig, M., Davis, C.L., Faeder, J.R., et al. (2021). Iterative community-driven development of a SARS-CoV-2 tissue simulator. bioRxiv **[preprint]**. 10.1101/2020.04.02.019075.
9. Risner, K.H., Tieu, K.V., Wang, Y., Bakovic, A., Alem, F., Bhalla, N., Nathan, S., Conway, D.E., Macklin, P., and Narayanan, A. (2020). Maraviroc inhibits SARS-CoV-2 multiplication and s-protein mediated cell fusion in cell culture. bioRxiv **[preprint]**. 10.1101/2020.08.12.246389.
10. Ozik, J., Collier, N., Heiland, R., An, G., and Macklin, P. (2019). Learning-accelerated discovery of immune-tumour interactions. Mol Syst Des Eng *4*, 747-760. 10.1039/c9me00036d.
11. Letort, G., Montagud, A., Stoll, G., Heiland, R., Barillot, E., Macklin, P., Zinovyev, A., and Calzone, L. (2019). PhysiBoSS: a multi-scale agent-based modelling framework integrating physical dimension and cell signalling. Bioinformatics *35*, 1188-1196. 10.1093/bioinformatics/bty766.
12. Ozik, J., Collier, N., Wozniak, J.M., Macal, C., Cockrell, C., Friedman, S.H., Ghaffarizadeh, A., Heiland, R., An, G., and Macklin, P. (2018). High-throughput cancer hypothesis testing with an integrated PhysiCell-EMEWS workflow. BMC Bioinformatics *19*, 483. 10.1186/s12859-018-2510-x.
13. Ghaffarizadeh, A., Heiland, R., Friedman, S.H., Mumenthaler, S.M., and Macklin, P. (2018). PhysiCell: An open source physics-based cell simulator for 3-D multicellular systems. PLoS Comput Biol *14*, e1005991. 10.1371/journal.pcbi.1005991.
14. Juarez, E.F., Lau, R., Friedman, S.H., Ghaffarizadeh, A., Jonckheere, E., Agus, D.B., Mumenthaler, S.M., and Macklin, P. (2016). Quantifying differences in cell line population dynamics using CellPD. BMC Systems Biology *10*. 10.1186/s12918-016-0337-5.
15. Macklin, P., Mumenthaler, S., and Lowengrub, J. (2013). Modeling Multiscale Necrotic and Calcified Tissue Biomechanics in Cancer Patients: Application to Ductal Carcinoma In Situ (DCIS). In Multiscale Computer Modeling in Biomechanics and Biomedical Engineering, pp. 349-380. 10.1007/8415_2012_150.
16. Majno, G., and Joris, I. (1995). Apoptosis, oncosis, and necrosis. An overview of cell death. Am J Pathol *146*, 3-15.

17. Krysko, D.V., Vanden Berghe, T., D'Herde, K., and Vandenabeele, P. (2008). Apoptosis and necrosis: detection, discrimination and phagocytosis. Methods *44*, 205-221. 10.1016/j.ymeth.2007.12.001.
18. Kerr, J.F., Winterford, C.M., and Harmon, B.V. (1994). Apoptosis. Its significance in cancer and cancer therapy. Cancer *73*, 2013-2026. 10.1002/1097-0142(19940415)73:8<2013::aid-cncr2820730802>3.0.co;2-j.
19. Hengartner, M.O. (2000). The biochemistry of apoptosis. Nature *407*, 770-776. 10.1038/35037710.
20. Garland, J.M., and Halestrap, A. (1997). Energy metabolism during apoptosis. Bcl-2 promotes survival in hematopoietic cells induced to apoptose by growth factor withdrawal by stabilizing a form of metabolic arrest. J Biol Chem *272*, 4680-4688. 10.1074/jbc.272.8.4680.
21. Macklin, P., Edgerton, M.E., Thompson, A.M., and Cristini, V. (2012). Patient-calibrated agent-based modelling of ductal carcinoma in situ (DCIS): from microscopic measurements to macroscopic predictions of clinical progression. J Theor Biol *301*, 122-140. 10.1016/j.jtbi.2012.02.002.
22. Ghaffarizadeh, A., Friedman, S.H., and Macklin, P. (2016). BioFVM: an efficient, parallelized diffusive transport solver for 3-D biological simulations. Bioinformatics *32*, 1256-1258. 10.1093/bioinformatics/btv730.
23. Mirams, G.R., Arthurs, C.J., Bernabeu, M.O., Bordas, R., Cooper, J., Corrias, A., Davit, Y., Dunn, S.J., Fletcher, A.G., Harvey, D.G., et al. (2013). Chaste: an open source C++ library for computational physiology and biology. PLoS Comput Biol *9*, e1002970. 10.1371/journal.pcbi.1002970.
24. Kang, S., Kahan, S., McDermott, J., Flann, N., and Shmulevich, I. (2014). Biocellion: accelerating computer simulation of multicellular biological system models. Bioinformatics *30*, 3101-3108. 10.1093/bioinformatics/btu498.
25. Hoehme, S., and Drasdo, D. (2010). A cell-based simulation software for multi-cellular systems. Bioinformatics *26*, 2641-2642. 10.1093/bioinformatics/btq437.
26. Abbasi, A., Amjad-Iranagh, S., and Dabir, B. (2022). CellSys: An open-source tool for building initial structures for bio-membranes and drug-delivery systems. J Comput Chem *43*, 331-339. 10.1002/jcc.26793.
27. Cytowski, M., Szymańska, Z., Umiński, P., Andrejczuk, G., and Raszkowski, K. (2017). Implementation of an Agent-Based Parallel Tissue Modelling Framework for the Intel MIC Architecture. Scientific Programming *2017*, 1-11. 10.1155/2017/8721612.
28. Mathias, S., Coulier, A., Bouchnita, A., and Hellander, A. (2020). Impact of Force Function Formulations on the Numerical Simulation of Centre-Based Models. Bulletin of Mathematical Biology *82*. 10.1007/s11538-020-00810-2.
29. van Leeuwen, I.M.M., Mirams, G.R., Walter, A., Fletcher, A., Murray, P., Osborne, J., Varma, S., Young, S.J., Cooper, J., Doyle, B., et al. (2009). An integrative computational model for intestinal tissue renewal. Cell Proliferation *42*, 617-636. 10.1111/j.1365-2184.2009.00627.x.
30. Meineke, F.A., Potten, C.S., and Loeffler, M. (2001). Cell migration and organization in the intestinal crypt using a lattice-free model. Cell Proliferation *34*, 253-266. 10.1046/j.0960-7722.2001.00216.x.
31. Segovia-Juarez, J.L., Ganguli, S., and Kirschner, D. (2004). Identifying control mechanisms of granuloma formation during M. tuberculosis infection using an agent-based model. J Theor Biol *231*, 357-376. 10.1016/j.jtbi.2004.06.031.
32. Wang, Y., Bergman, D., Trujillo, E., Pearson, A.T., Sweis, R.F., and Jackson, T.L. (2023). Mathematical Model Predicts Tumor Control Patterns Induced by Fast and Slow CTL Killing Mechanisms. bioRxiv [preprint] *2023.07.19.548738*. 10.1101/2023.07.19.548738.
33. Osinska, I., Popko, K., and Demkow, U. (2014). Perforin: an important player in immune response. Cent Eur J Immunol *39*, 109-115. 10.5114/ceji.2014.42135.
34. Farhood, B., Najafi, M., and Mortezaee, K. (2019). CD8(+) cytotoxic T lymphocytes in cancer immunotherapy: A review. J Cell Physiol *234*, 8509-8521. 10.1002/jcp.27782.
35. Raskov, H., Orhan, A., Christensen, J.P., and Gogenur, I. (2021). Cytotoxic CD8(+) T cells in cancer and cancer immunotherapy. Br J Cancer *124*, 359-367. 10.1038/s41416-020-01048-4.

36. Norton, K.A., Gong, C., Jamalian, S., and Popel, A.S. (2019). Multiscale Agent-Based and Hybrid Modeling of the Tumor Immune Microenvironment. Processes (Basel) *7*. 10.3390/pr7010037.
37. Anderson, A.R.A. (2007). A Hybrid Multiscale Model of Solid Tumour Growth and Invasion: Evolution and the Microenvironment. In Single-Cell-Based Models in Biology and Medicine, pp. 3-28. 10.1007/978-3-7643-8123-3_1.
38. Hoehme, S., Friebel, A., Hammad, S., Drasdo, D., and Hengstler, J.G. (2017). Creation of Three-Dimensional Liver Tissue Models from Experimental Images for Systems Medicine. Methods Mol Biol *1506*, 319-362. 10.1007/978-1-4939-6506-9_22.
39. Finley, S.D., and Popel, A.S. (2013). Effect of tumor microenvironment on tumor VEGF during anti-VEGF treatment: systems biology predictions. J Natl Cancer Inst *105*, 802-811. 10.1093/jnci/djt093.
40. Swan, A., Hillen, T., Bowman, J.C., and Murtha, A.D. (2018). A Patient-Specific Anisotropic Diffusion Model for Brain Tumour Spread. Bull Math Biol *80*, 1259-1291. 10.1007/s11538-017-0271-8.
41. Chaplain, M.A., Graziano, L., and Preziosi, L. (2006). Mathematical modelling of the loss of tissue compression responsiveness and its role in solid tumour development. Math Med Biol *23*, 197-229. 10.1093/imammb/dql009.
42. Alarcon, T., Byrne, H.M., and Maini, P.K. (2003). A cellular automaton model for tumour growth in inhomogeneous environment. J Theor Biol *225*, 257-274. 10.1016/s0022-5193(03)00244-3.
43. Scott, J.G., Basanta, D., Anderson, A.R., and Gerlee, P. (2013). A mathematical model of tumour self-seeding reveals secondary metastatic deposits as drivers of primary tumour growth. J R Soc Interface *10*, 20130011. 10.1098/rsif.2013.0011.
44. Kaznatcheev, A., Vander Velde, R., Scott, J.G., and Basanta, D. (2017). Cancer treatment scheduling and dynamic heterogeneity in social dilemmas of tumour acidity and vasculature. Br J Cancer *116*, 785-792. 10.1038/bjc.2017.5.
45. Poleszczuk, J., Hahnfeldt, P., and Enderling, H. (2014). Biphasic modulation of cancer stem cell-driven solid tumour dynamics in response to reactivated replicative senescence. Cell Prolif *47*, 267-276. 10.1111/cpr.12101.
46. Powathil, G.G., Adamson, D.J., and Chaplain, M.A. (2013). Towards predicting the response of a solid tumour to chemotherapy and radiotherapy treatments: clinical insights from a computational model. PLoS Comput Biol *9*, e1003120. 10.1371/journal.pcbi.1003120.
47. Hamis, S., Nithiarasu, P., and Powathil, G.G. (2018). What does not kill a tumour may make it stronger: In silico insights into chemotherapeutic drug resistance. J Theor Biol *454*, 253-267. 10.1016/j.jtbi.2018.06.014.
48. Fortuna, I., Perrone, G.C., Krug, M.S., Susin, E., Belmonte, J.M., Thomas, G.L., Glazier, J.A., and de Almeida, R.M.C. (2020). CompuCell3D Simulations Reproduce Mesenchymal Cell Migration on Flat Substrates. Biophys J *118*, 2801-2815. 10.1016/j.bpj.2020.04.024.
49. Dunn, S.J., Appleton, P.L., Nelson, S.A., Nathke, I.S., Gavaghan, D.J., and Osborne, J.M. (2012). A two-dimensional model of the colonic crypt accounting for the role of the basement membrane and pericryptal fibroblast sheath. PLoS Comput Biol *8*, e1002515. 10.1371/journal.pcbi.1002515.
50. Glen, C.M., Kemp, M.L., and Voit, E.O. (2019). Agent-based modeling of morphogenetic systems: Advantages and challenges. PLoS Comput Biol *15*, e1006577. 10.1371/journal.pcbi.1006577.
51. Schubert, M., Dokmegang, J., Yap, M.H., Han, L., Cavaliere, M., and Doursat, R. (2021). Computational modelling unveils how epiblast remodelling and positioning rely on trophectoderm morphogenesis during mouse implantation. Plos One *16*. 10.1371/journal.pone.0254763.
52. Camacho-Gómez, D., García-Aznar, J.M., and Gómez-Benito, M.J. (2022). A 3D multi-agent-based model for lumen morphogenesis: the role of the biophysical properties of the extracellular matrix. Engineering with Computers *38*, 4135-4149. 10.1007/s00366-022-01654-1.
53. Cess, C.G., and Finley, S.D. (2020). Multi-scale modeling of macrophage—T cell interactions within the tumor microenvironment. PLOS Computational Biology *16*. 10.1371/journal.pcbi.1008519.
54. Ruiz-Martinez, A., Gong, C., Wang, H., Sove, R.J., Mi, H., Kimko, H., and Popel, A.S. (2022). Simulations of tumor growth and response to immunotherapy by coupling a spatial agent-based

model with a whole-patient quantitative systems pharmacology model. PLoS Comput Biol *18*, e1010254. 10.1371/journal.pcbi.1010254.
55. Ni, C., and Lu, T. (2022). Individual-Based Modeling of Spatial Dynamics of Chemotactic Microbial Populations. ACS Synth Biol *11*, 3714-3723. 10.1021/acssynbio.2c00322.
56. Hellweger, F.L., and Bucci, V. (2009). A bunch of tiny individuals—Individual-based modeling for microbes. Ecological Modelling *220*, 8-22. 10.1016/j.ecolmodel.2008.09.004.
57. Hastings, J., Owen, G., Dekker, A., Ennis, M., Kale, N., Muthukrishnan, V., Turner, S., Swainston, N., Mendes, P., and Steinbeck, C. (2016). ChEBI in 2016: Improved services and an expanding collection of metabolites. Nucleic Acids Res *44*, D1214-1219. 10.1093/nar/gkv1031.
58. Degtyarenko, K., de Matos, P., Ennis, M., Hastings, J., Zbinden, M., McNaught, A., Alcantara, R., Darsow, M., Guedj, M., and Ashburner, M. (2008). ChEBI: a database and ontology for chemical entities of biological interest. Nucleic Acids Res *36*, D344-350. 10.1093/nar/gkm791.
59. Cook, D.L., Mejino, J.L., Neal, M.L., and Gennari, J.H. (2008). Bridging biological ontologies and biosimulation: the ontology of physics for biology. AMIA Annu Symp Proc *2008*, 136-140.
60. Gkoutos, G.V., Mungall, C., Dolken, S., Ashburner, M., Lewis, S., Hancock, J., Schofield, P., Kohler, S., and Robinson, P.N. (2009). Entity/quality-based logical definitions for the human skeletal phenome using PATO. 2009 Annual International Conference of the IEEE Engineering in Medicine and Biology Society.
61. PhysiCell (2023). PhysiCell Version 1.12.0. https://github.com/MathCancer/PhysiCell/releases/tag/1.12.0.
